# The Dynamics of Adaptive Genetic Diversity During the Early Stages of Clonal Evolution

**DOI:** 10.1101/170589

**Authors:** Jamie R. Blundell, Katja Schwartz, Danielle Francois, Daniel S. Fisher, Gavin Sherlock, Sasha F. Levy

## Abstract

The dynamics of genetic diversity in large clonally-evolving cell populations are poorly understood, despite having implications for the treatment of cancer and microbial infections. Here, we combine barcode lineage tracking, sequencing of adaptive clones, and mathematical modelling of mutational dynamics to understand diversity changes during experimental evolution. We find that, despite differences in beneficial mutational mechanisms and fitness effects between two environments, early adaptive genetic diversity increases predictably, driven by the expansion of many single-mutant lineages. However, a crash in diversity follows, caused by highly-fit double-mutants fed from exponentially growing single-mutants, a process closely related to the classic Luria-Delbruck experiment. The diversity crash is likely to be a general feature of clonal evolution, however its timing and magnitude is stochastic and depends on the population size, the distribution of beneficial fitness effects, and patterns of epistasis.

---

In large clonally-evolving populations, lineages harboring beneficial “driver” mutations expand, compete with one another, and acquire further beneficial mutations, shaping genetic diversity [1–3]. Recent studies employing deep genomic sequencing have shown that large laboratory [4–6] and clinical [7–13] cell populations harbor high levels of genetic diversity that changes through time. In disease-relevant scenarios, such as cancer [7–11] and within-host microbial dynamics [13], the timescale over which diversity builds up is often short, such that dominant clones only accumulate a handful of driver mutations. When the supply of driver mutations is low, evolution is characterized by successive selective sweeps, wherein a single adaptive lineage periodically purges genetic diversity [14, 15]. However, when the supply of driver mutations is high, evolution is characterized by clonal interference, with multiple adaptive lineages expanding and competing through time [5, 16–19]. In the clonal interference regime, mutations often rise and fall together in cohorts [5, 11, 20]. However, due to the limited ability to detect low frequency mutations via genomic sequencing, it remains unclear what controls diversity changes through time in this regime, whether large purges of diversity might also occur as in the case of a selective sweep, and whether these diversity crashes are predictable across replicates and environments [21].

To address these questions, we introduced ~500,000 unique DNA barcodes into *S. cerevisiae* [22] and evolved populations of ~5 × 10^8^ cells in triplicate for 304 generations under two well-mixed nutrient-limited environments: nitrogen-limitation (N-lim) and carbon-limitation (C-lim). Lineage abundances were tracked by barcode sequencing every ~8-24 generations. Lineages that harbor adaptive mutations were identified by finding those trajectories that deviate from a neutral expectation [22]. Tracking all adaptive lineages through time reveals the changing levels of adaptive lineage diversity (Figure 1). Initially, adaptive diversity expands, driven by thousands of independent mutations. This expansion is quantitatively different between environments; in N-lim, the expansion is slower and fewer lineages reach high frequencies. Later however, similarities between environments emerge: a handful of lineages begin to dominate the population, causing a crash in adaptive lineage diversity.

**Figure 1.**
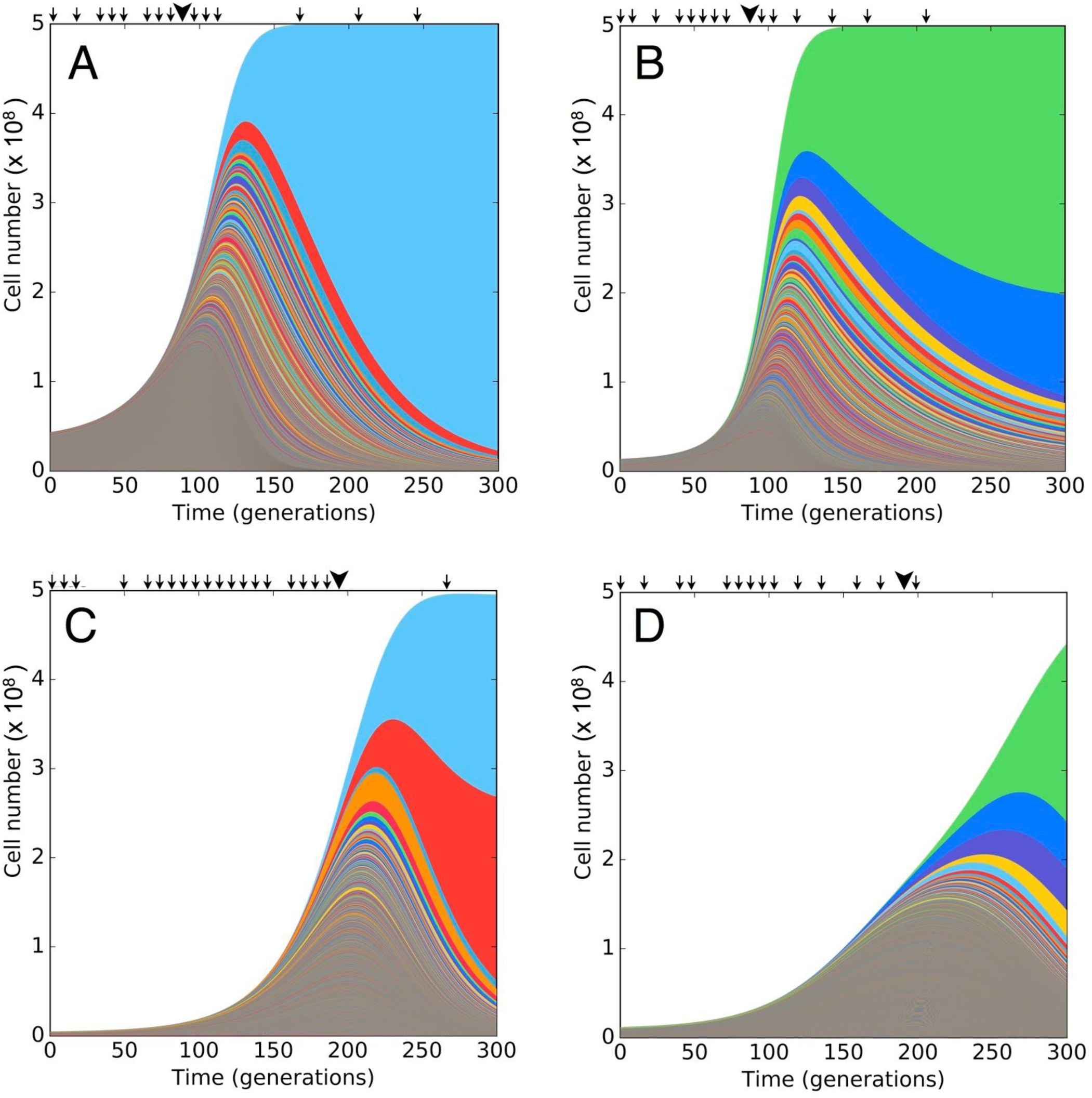
“Muller” plots of adaptive lineages. The cell number of all adaptive lineages (colors) inferred from barcode sequencing (arrows) and whole genome sequencing of picked clones (large arrowheads) of replicate evolutions in carbon-limited (A and B) and nitrogen-limited media (C and D). Colors are for visualization purposes only, and do not represent lineages harbouring specific mutations.

We suspected that differences in lineage diversity dynamics between the two environments could be attributed, in part, to differences in the <u>m</u>utational <u>D</u>istribution of <u>F</u>itnesses <u>E</u>ffects (mDFE), defined as the distribution of mutation rates over fitness effects for single beneficial mutations arising on the ancestor. We have previously shown that high-resolution lineage tracking over short times can be used to infer the mDFE [22]. In C-lim, the mDFE results in ~10^4^ beneficial mutations with fitness effects (s) above 3% entering the population over the first ~100 generations. This initially produces a quasi-deterministic expansion in diversity because the low fitness effect beneficial mutations that dominate early occur at a high rate. Later, the diversity expansion becomes more stochastic because the high fitness effect beneficial mutations that dominate during this time occur at a lower rate (i.e. the effective beneficial mutation rate decreases, SM2.1). To test whether these features might generalize to other environments we inferred the mDFE for N-lim (SM1). We find that the shape of the mDFE in N-lim is qualitatively different from that in C-lim (Figure 2). Most strikingly, the rates of mutation to higher fitness effects (s>5%) are ~3-fold lower in N-lim, and fall off rapidly, resulting in no detectable fitness effects above 8%. With time, the lineage dynamics become exponentially more sensitive to these differences at the higher fitness effects (expanded region Figure 2 and SM2). In C-lim, these mutations establish (escaping stochastic loss), expand, and compete over shorter timescales (Figure 1 A vs. C). In N-lim, the lower rate of mutation to higher fitness effects results in even more stochastic dynamics: the highly fit mutations occur in smaller numbers, causing larger variations between replicates (Figure 1 C vs. D, SM2.1).

**Figure 2.**
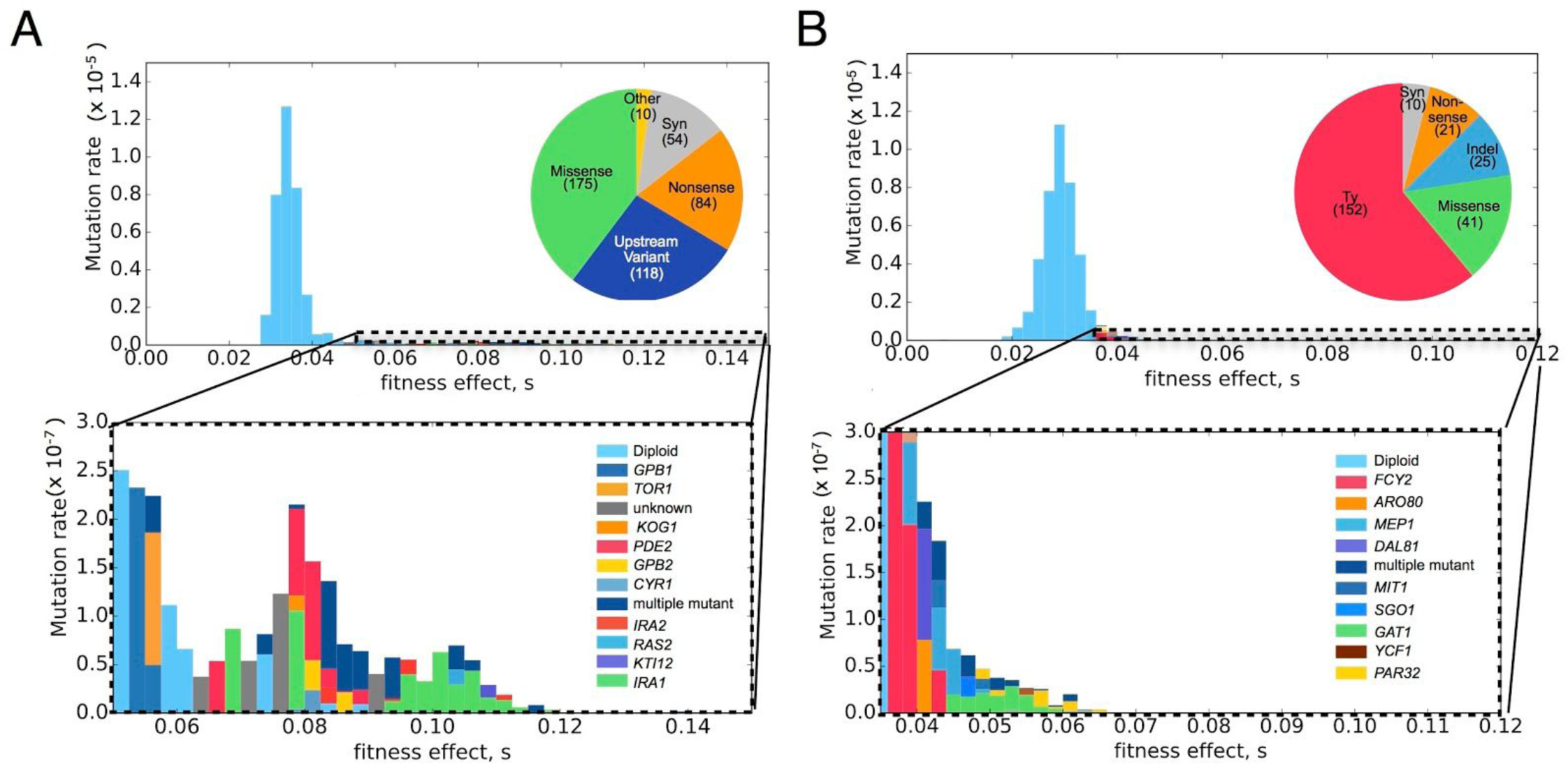
Using barcode-directed whole-genome sequencing of adaptive clones to find the mutational targets underlying the distribution of beneficial fitness effects (mDFE) in (A) carbon- and (B) nitrogen-limitation. Most adaptive events are diploidizations (top, light blue), but high fitness effects are caused by mutations in or near genes (below, color key). Each fitness bin is colored according the to the estimated rate of mutation of verified single-mutants to each gene in that bin (SM3.3). The color key is roughly ordered from lowest to highest fitness effect of a mutational event. Pie charts indicate the mutational mechanisms of adaptive mutations.

To verify that single beneficial mutations are the determinants of the early diversity dynamics, we used barcode-directed whole-genome sequencing to identify hundreds of unique adaptive mutations that span a wide range of fitness effects. In C-lim, we previously found a near-comprehensive spectrum of adaptation-driving single-mutations by sequencing 418 adaptive clones isolated from generation 88 [23]. Here, we repeated this for N-lim, sequencing 291 adaptive clones picked from generation 192 and re-measuring fitness (SM3.2). In both environments, the majority of sequenced adaptive clones contain a single adaptive mutation (>75%), consistent with single-mutants being the determinants of early diversity dynamics. Lineages containing two adaptive mutations are crucial to the later-time diversity dynamics and we return to these below. Focusing on single-mutant clones, we find major differences in mutational mechanisms and the mutational targets of adaptation (Figure 2). Surprisingly, Ty transposition events play a major role in driving adaptation in N-lim but not C-lim. In both environments, adaptation is driven first by cells that undergo a frequent diploidization (Dip) event, and later by cells that acquire mutations in a small set of nutrient sensing pathway genes (Figure 2) and [16]. The majority of these recurrently mutated genes are putative loss-of-function (LoF) mutations, with a minority being putative gain-of-function (GoF) mutations (SM3.4). Additionally, there are a significant number of adaptive mutations in singly mutated genes that do not appear to function in nutrient sensing pathways, suggesting cells have many pathways to increase fitness in N-lim (Figure 2B).

In each evolution, we observed a lineage diversity crash whereby a handful of lineages outcompete all others (Figure 1). One possibility is that multiple adaptive clones within each lineage contribute to a lineage’s dominance. Alternatively, a single large clone within each lineage may be primarily responsible for this dominance. However, it is unclear how such large clones would arise. To investigate these questions, we simulated the diversity dynamics using the mDFE inferred in each environment, removing any lineages which were found to contain two adaptive mutations occurring in the same cell (double-mutants), and tracked both the lineage and clone diversity (SM4). To make comparisons of genetic diversity across a large number of simulations, we plot the Shannon entropy [24] of adaptive lineages and clones through time, which track one another closely until the time at which triple- and quadruple-mutant clones first expand within barcoded lineages (SM2.5). Simulations accounting only for the stochastic occurrence of, and competition between, single-mutants (single mutant model) predict that diversity should crash slowly, at odds with observations (Figure 3A). We therefore reasoned that the diversity crash is caused by the emergence of double-mutants. To test this, we modified our simulations to allow for multiple-mutants drawn from the single-mutant mDFE and whose fitness effects combine additively (additive model, Figure 3B). Additive model simulations produce diversity crashes that are caused by a handful of lineages that each contain a single dominant double-mutant clone (typically <5 lineages are >90% of the population at times beyond 150 generations, SM4).

**Figure 3.**
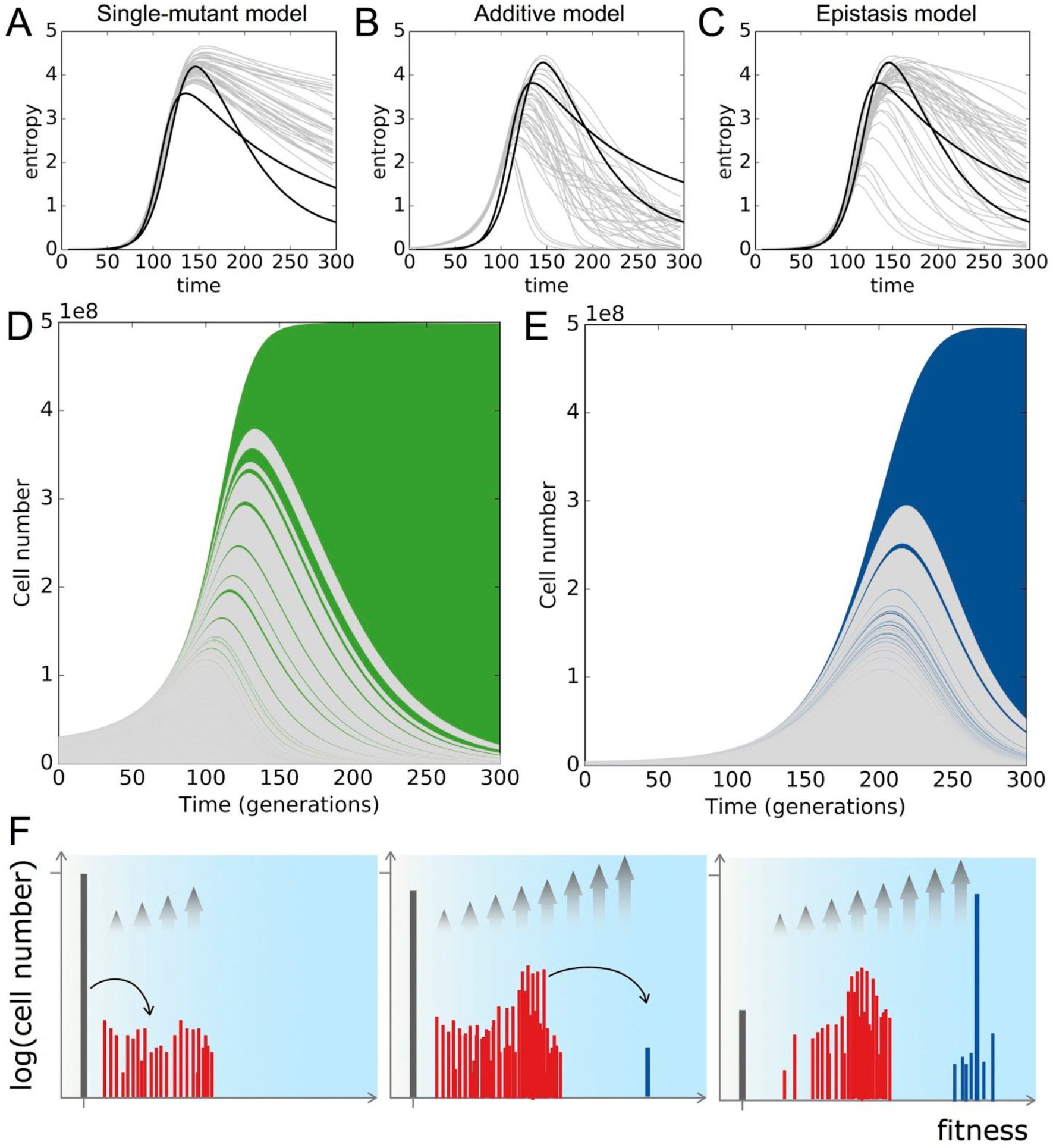
Exponential feeding of double mutants causes a diversity crash. (A-C) Shannon entropy of adaptive lineages from replicate experiments in carbon-limitation (black lines) and stochastic simulations (grey lines) using a single-mutant model (A), additive model (B), or epistasis model (C). The entropy of adaptive clones (not shown) closely tracks the entropy of adaptive lineages (*SM 2.5*). (D,E) Muller plots from Figure 1 recolored to depict single-mutant (grey) and early double-mutant (green and blue) adaptive lineages in (D) carbon- and (E) nitrogen-limitation. (F) Schematic showing the statistical behavior of the ancestor (grey), first (red), and second (blue) mutants. Grey arrows show the relative growth rates due to selection.

To understand why a handful of large double-mutant clones cause a diversity crash we considered a model of the mutation acquisition process (SM2) and [17]. Single-mutants establish in large numbers because the product of the population size (N = 5 × 10^8^) and the effective beneficial mutation rate (U_b_>~10^−7^, see SM2.1) is large (NU_b_>50). However, since the probability of two mutations occurring concurrently is small (NU_b_^2^<<1), double mutants must enter stochastically on the background of exponentially growing single-mutants. By considering when an expanding single-mutant clone will typically give rise to a double-mutant (following similar arguments presented in [15, 17]), we find that the distribution of sizes (n) of double-mutant clones is a power law ~1/n^(2+Δ)/(1+Δ)^, where Δ is the fitness effect of the second mutation over the fitness effect of the first mutation (s_2_/s_1_). If double-mutants are no fitter than single-mutants (Δ=0), this collapses to the classic Luria-Delbruck distribution of clone sizes, ~1/n^2^ [25], whereby many double-mutant cells belong to a large number of small clones. However, when double-mutants are significantly fitter than single-mutants (Δ>1), most double-mutant cells belong to a handful of large clones (SM2). In this second case, a few lucky double-mutants with the earliest occurrence times go on to dominate the population, but stochasticity in their occurrence times results in variability in the timing and depth of the diversity crash between replicates (Figure 3F, SM4).

To validate that a handful of double-mutants cause the diversity crashes observed in our experiments, we examined the trajectories of lineages containing a confirmed double-mutant and verified that they do indeed go on to dominate (data in Figure 3D & E). Surprisingly, however, the sequencing of clones revealed that dominant double-mutants were not composed of two high fitness effect mutations (e.g. LoF+LoF) as would be predicted by our additive model simulations (SM4). Instead, dominant clones that were sequenced were Dip+GoF double-mutants (Dip+*ras2* in C1; Dip+*mep1* in N1), despite neither GoF mutation occurring at a high rate. We reasoned that this could be caused by epistasis: some classes of beneficial mutations combine non-additively and these interactions must play a crucial role in determining which mutations go on to dominate. To test this, we modified our additive model simulations (above) to ban second mutations that are implausible or unobserved (Dip+Dip, Dip+LoF, LoF+LoF, or LoF+GoF, SM4). Simulations using this “epistasis model” produce diversity crashes and lineage trajectories that are more consistent with observations (Figure 3C). Importantly, the epistasis model predicts that clones driving the diversity crash will usually be Dip+GoF, if the crash is deep, and LoF+Dip if the crash is shallow.

Lineage trajectories alone have limited power to distinguish between the additive and epistasis models (Figures 3B & C). To further test our models, we therefore asked if the dynamics of mutations, rather than lineages, are consistent with predictions of either model. We measured the abundance of diploids in the population every 8-24 generations using a colony-growth assay (SM6) and [23], not only for the 4 evolutions described above, but for two additional evolutions (1 in C-lim, 1 in N-lim) not characterized by lineage tracking (Figure 4B-E). Consistent with our observations, both models predict that replicate diploid trajectories will track each other closely, first, as large numbers of ancestral cells diploidize and expand, and second, as diploids begin to be out-competed by haploids that have acquired fitter LoF and GoF mutations. At later times, however, the models deviate. In C-lim, the additive model predicts that LoF+LoF or LoF+GoF double-mutants drive the continuing decline of diploids (Figure 4B). In N-lim, the additive model predicts that Dip+LoF mutations should expand fast enough that diploid trajectories never dip (Figure 4D). However, consistent with observations, the epistasis model predicts that the diploid trajectory will dip and subsequently recover, driven by LoF and GoF haploids that diploidize and by diploids that acquire GoF mutations (Figures 4C & E). Since this diploid recovery is driven by rare double-mutants, its timing and depth are predicted to be highly stochastic, resulting in large variations between replicates, in agreement with our data. In cases where diploids recover early (e.g. C1, N1 and N3), our model correctly predicts that the recovery is likely driven by Dip+GoF double-mutants, largely because this event has the highest Δ value (SM2).

**Figure 4.**
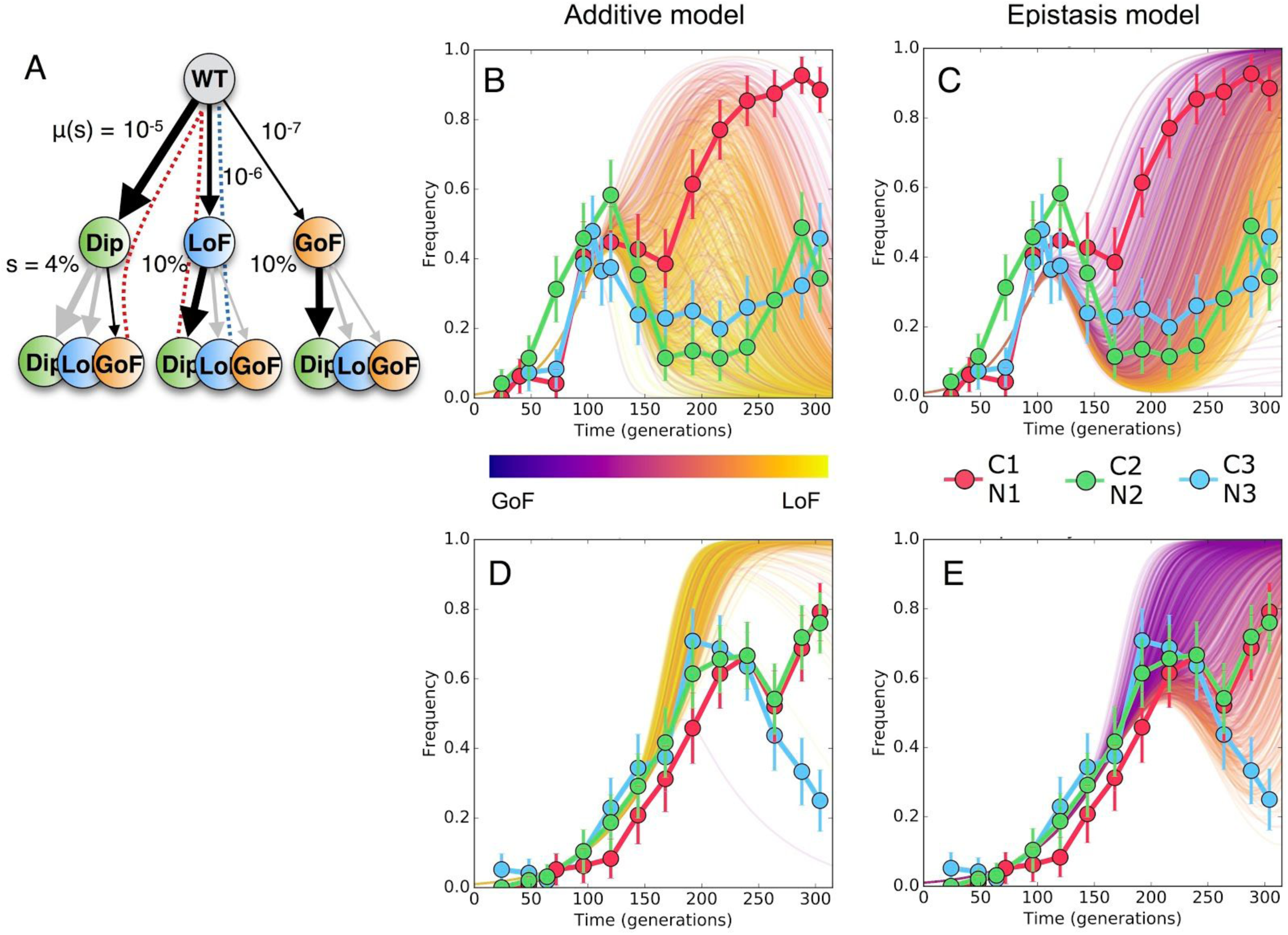
Simulations of diploid dynamics using the additive and epistasis models. (A) The simplified fitness landscape used for simulations in C-lim, where u(s) indicates the mutation rates and s indicates the fitness effects to Diploidy (Dip), Loss-of-Function (LoF), and Gain-of-Function (GoF) mutations. Greyed arrows are paths disallowed by the epistasis model. Dashed lines indicate the paths taken by the dominant clones in the additive (blue) epistasis models (red). (C-E) The diploid trajectories in C-lim and N-lim predicted by the additive model (B and D, respectively) and the epistasis model (C and E, respectively) compared to the measured diploid trajectories (data points) from the three replicate populations in each condition. Color scale indicates the extent to which the diploid rescue is driven by Dip+GoF (purple) vs. LoF+Dip or Dip+LoF (yellow) mutants, with early rescue being more likely to be driven by Dip+GoF mutations.

We have shown that the genetic diversity in our experiments first increases quasi-deterministically, caused by a large number of single-mutants, and later crashes stochastically, caused by competition from a handful of highly-fit double-mutants that occur anomalously early. While the expansion rate and the timing and depth of the crash are influenced by the population size, the mDFE, and the pattern of epistasis between adaptive mutations, they are likely to be general features of clonal evolution. In environments such as our experiments where the mDFE lacks extremely high fitness effect mutations, mutations that cause the diversity expansion are being fed from a large, effectively-constant population, while mutations that cause the crash are being fed from a small, exponentially-growing population. In environments where extremely high fitness effect mutations are possible, such as in the presence of a growth-inhibiting drug [26], a diversity expansion and crash will still occur, but the crash will sometimes be driven by the expansion of a highly fit single-mutant (SM2.6 and SM2.7). That is, a diversity crash is likely to occur, whether driven by a well-described selective sweep of a single mutant [14, 15] or by a multiple mutant that occurs anomalously early. While we have focused on well-mixed populations, a qualitatively similar phenomenon has been described during the expansion of spatially structured populations [27]. More generally, our work highlights that while genetic diversity evolves in a highly stochastic way and depends on rare events, the statistics of these processes provide a means by which to forecast evolution [21].

While in our experiments the deterministic to stochastic transition occurs between first and second mutations, in small populations (NU_b_<1), first adaptive mutations are stochastic, and diversity crashes (of neutral mutations) will occur at nearly every adaptive event. In even larger populations (NU_b_^2^/s> 1 >NU_b_^3^/s^2^), double-mutants will occur deterministically, but triple-mutants stochastically, and therefore the diversity crash will be caused by a few triple-mutants. More generally, for mDFEs that lack a small supply of relatively large fitness effects (whose dynamics can be characterized by a single “predominant fitness class” [17]), the diversity crash will be driven by clones harbouring *k* beneficial mutations, where *k* is the smallest integer for which Ns(U_b_/s)^k^<1, a parameter which also controls the steady-state range of fitnesses present in an adapting population [17]. Previous work has found that “beneficial cohorts”, multiple beneficial mutations co-occurring in clones that are at frequencies below the detection limit of whole-genome or whole-exome sequencing, commonly drive laboratory [5, 20] and clinical [11] clonal evolution. Our results suggest that, at least during the early stages of clonal evolution, these cohorts are expected, with cohort size being determined by *k* (SM2). Furthermore, theoretical work predicts that beneficial cohorts and fluctuations in genetic diversity should occur throughout evolution, driven by the stochastic occurrence of anomalously early and/or fit multiple-mutants [28, 29]. Experimental validation of these predictions, however, will likely require the development of double-barcoding technologies [30] or barcodes that continuously generate diversity through time [31–33].

## Methods

### Experimental evolutions

Details of strain construction, media and equipment used in the evolutions can be found in the supplementary information of Levy et. al. [14].

### Barcode sequencing

Details of the DNA preps, PCR reactions and sequencing protocols can be found in the supplementary information of Levy et. al. [14] for the Carbon condition. Further relevant details for the Nitrogen condition can be found in Section 1.1 and 1.2 of the supplementary information accompanying this manuscript.

### Identifying adaptive lineages

Adaptive lineages were identified using the same methods and code as outlined in the supplementary information of Levy et. al. [14]

### Isolating and Sequencing of adaptive clones

All clones, including those from the Nitrogen conditions, were isolated and sequenced following the protocol outlined in detail in the “Star methods” section of Venkataram et. al. [16]. For further details of the sequencing of the Nitrogen clones we refer the reader also to Sections 3.1 and 3.2 of the supplementary information accompanying this manuscript.

### Simulated lineage dynamics

Details of the simulations can be found in Sections 2 and 4 of the supplementary information accompanying this manuscript.

## Acknowledgements

We wish to thank all members of the Levy, Sherlock and Fisher labs for useful discussions and comments. J.R.B. is supported NSF PHY-1545840, Stand Up 2 Cancer and by the Louis and Beatrice Laufer Center; D.S.F. by NSF PHY-1305433, NSF PHY-1545840, Stand Up 2 Cancer and NIH R01 HG003328; G.S by NIH grants R01 HG003328 and GM110275; and S.F.L by NIH grant R01 HG008354 and by the Louis and Beatrice Laufer Center. All data available on request.

## SUPPLEMENTARY MATERIALS

### 1 N-lim experiment and DFE

#### 1.1 Media used for N-lim condition

The barcoded yeast library from [1], containing ∼500,000 barcodes, was evolved by serial batch culture under nitrogen limitation in 100 ml of 5x Delft media [2] with 0.04% ammonium sulfate and 4% dextrose. Cells were grown in 500 ml Delong flasks (Bellco) at 30° C and 223 RPM for 48 hours between each bottleneck. Bottlenecks were performed by adding 400 *μ*l of the evolution to fresh media. Cell counts were performed at each bottleneck to estimate the generation time. Contamination checks for bacteria or other non-yeast microbes were performed regularly. Barcode sequencing and counting was performed, as described [1].

#### 1.2 Inferring the DFE in N-lim

##### Here we outline how we infer the DFE in N-lim and compare its features with those measured previously in C-lim

To construct the DFE in N-lim we followed the procedure outlined in detail in [1]. Because adaptation is slower in N-lim we used trajectories for ∼200 generations with read depths given in the following table:

**Table 1:** Read depths across time points for the two replicate evolutions in nitrogen limitation (N-lim). Only these time points are used in the analysis.

| Generation | 0 | 8 | 16 | 48 | 72 | 80 | 96 | 104 | 112 | 120 | 128 | 136 | 144 | 160 | 168 | 176 | 184 | 192 | 200 |
| --- | --- | --- | --- | --- | --- | --- | --- | --- | --- | --- | --- | --- | --- | --- | --- | --- | --- | --- | --- |
| Depth N1<br>($\times 10^6$) | 222 | 36 | 30 | 36 | 30 | 33 | 39 | 39 | 36 | 35 | 32 | 24 | 23 | 23 | 34 | 39 | 20 | 39 | – |
| Depth N2<br>($\times 10^6$) | 222 | – | 30 | 38 | 34 | 50 | 34 | 29 | – | 39 | – | 37 | – | 29 | – | 37 | – | 25 | 47 |

To identify lineages with beneficial mutations, using the same null model as outlined in [1], each barcode trajectory was assigned a posterior probability of harboring an established beneficial mutation with fitness effect *s* and establishment time **τ** based on its abundance change between subsequent time points. Lineage trajectories in both N-lim replicates (N1 and N2) are shown in Figure 1 alongside the original C-lim trajectories (C1 and C2) (previously published in [1]) for comparison. The more rapid adaptation observed in C-lim relative to N-lim can be quantified by plotting the mean-fitness trajectory (mean fitness of cells in the population relative to the ancestor), shown in the right-hand panel of Figure 1. These mean fitness trajectories are inferred using the fitness estimates and interpolated trajectories for each adaptive lineage identified in each environment.

**Figure 1:**
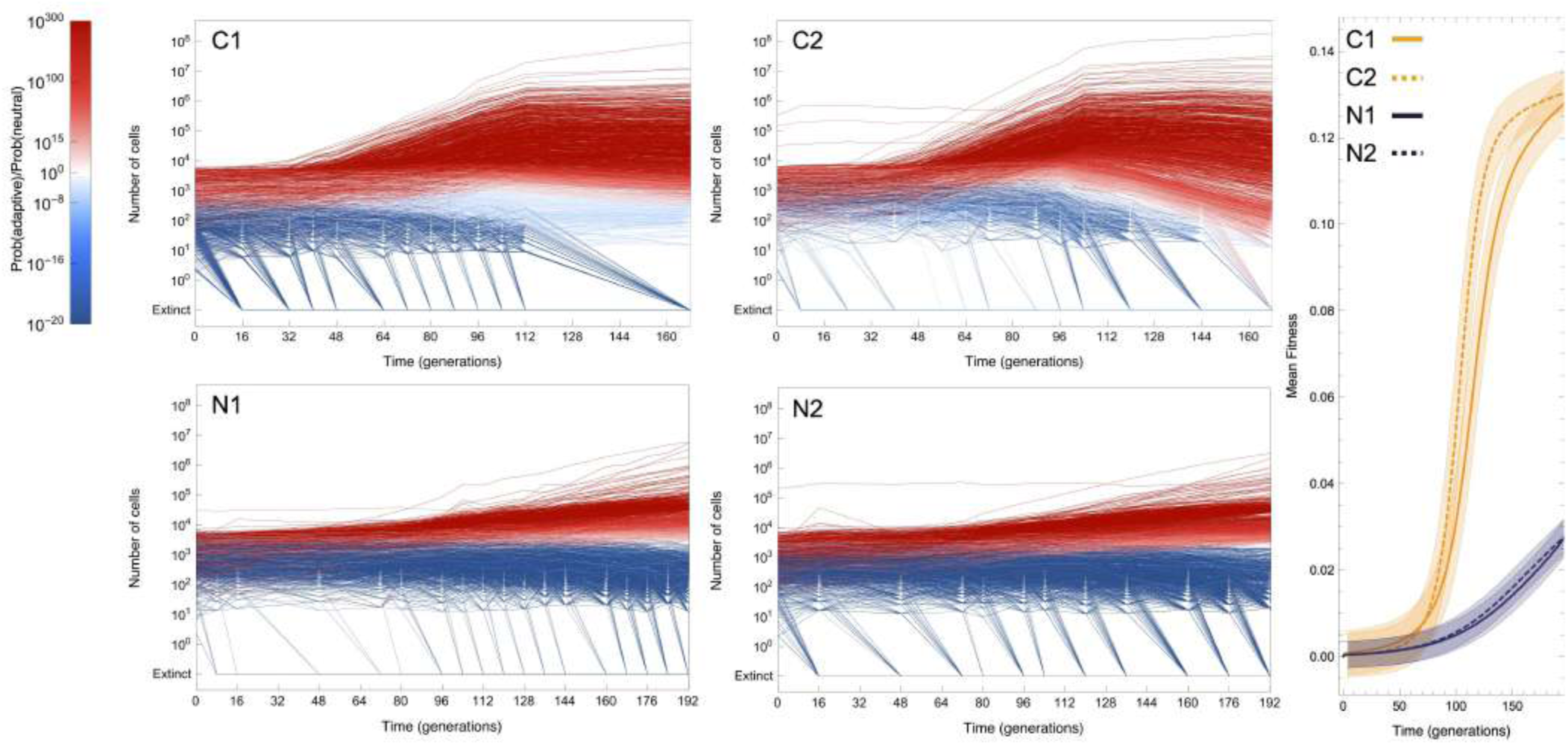
(Left) Lineage trajectories from each of the four replicates (C1, C2 performed in C-lim and N1, N2 performed in N-lim) for which lineage tracking was performed colored by probability of containing an adaptive mutation. (Right) The mean fitness of cells in each experiment over time relative to the wild-type. Replicates within the same environment are consistent with one another although do show some variation, particularly at later times, due to the stochastic occurrence time of mutations [1]. Shaded region indicates approximate error on the mean fitness inference.

To infer the DFE in N-lim, we first filtered out adaptive lineages that were identified as adaptive in both replicates (purple points in Figure 2) as these were likely pre-existing mutations that arose prior to the beginning of the experimental evolutions [1]. Figure 2 shows all beneficial mutations identified as adaptive colored according to whether they are pre-existing (purple) or not (green) in both N-lim replicates and, for comparison, the previously published C-lim replicates.

**Figure 2:**
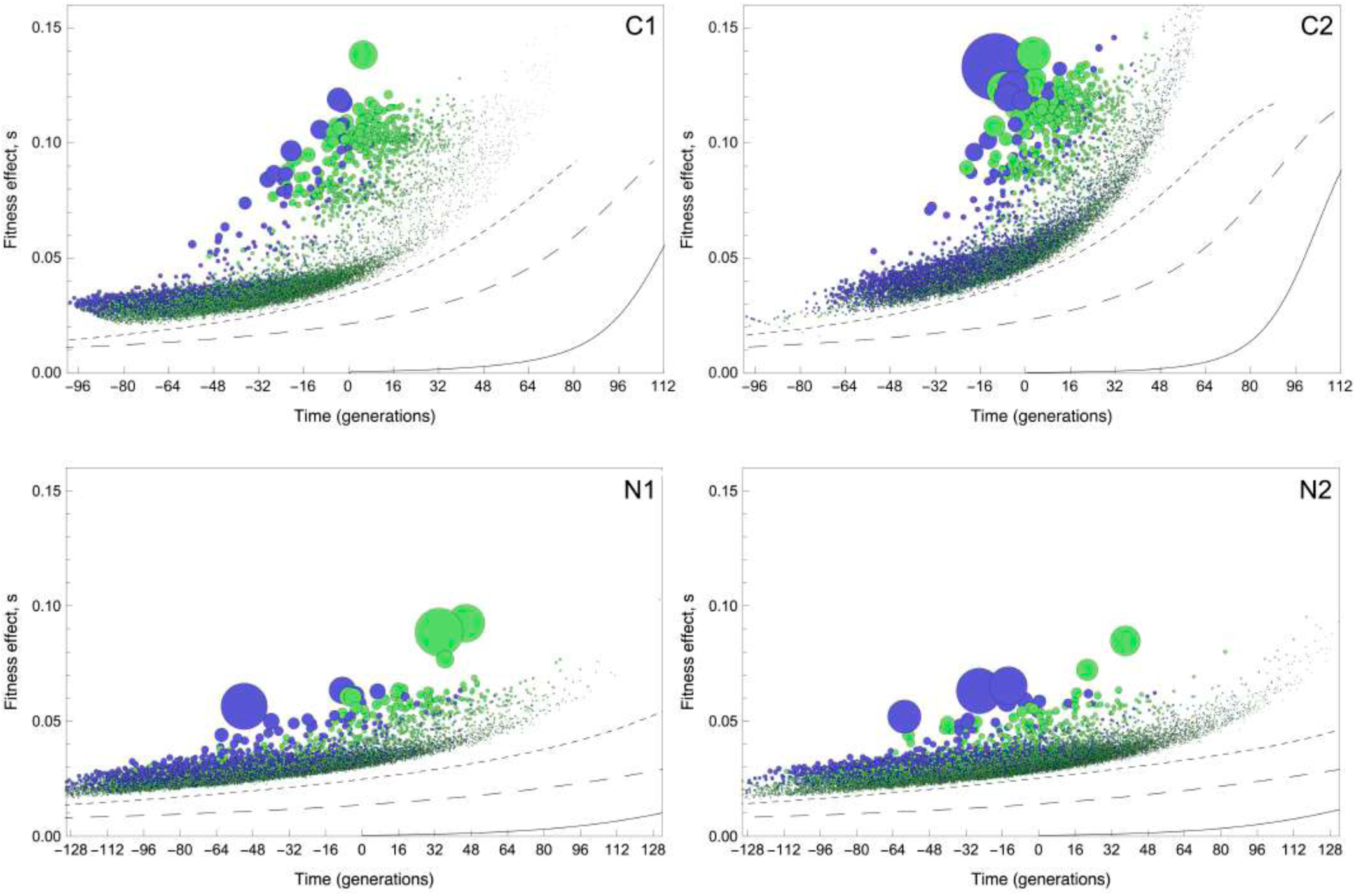
Expansion of lineages can be used to estimate the fitness effect *s* (y-axis) and establishment time, *τ*, (x-axis) of all detectable single beneficial mutations that enter the population. Each circle is an independently occurring mutation, the size indicates its abundance in the population at generation 88. Color indicates whether it entered prior to the separation of the replicates (purple, “pre-existing”) or after the separation (green, “not pre-existing”). The top panel is from previously published data in [1] and is shown here for comparison.

Next we used the estimates of *s* inferred for each adaptive lineage together with the deterministic approximation outlined in [1] (that relates the mutation rate density *μ*(*s*) to fitness effects in the range [*s*, *s* + *ds*] to the measured fraction *f*(*ds*, *t*) of cells in the population expanding at rates between [*s*, *s* + *ds*] at time *t*) to estimate the rate of mutation to each fitness interval via:

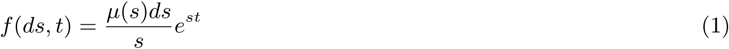

which is independent of the variance in offspring number [1]. The result of this procedure gives one an estimate for the mutation rate density as a function of fitness effect as shown here in Figure 3 and also in the main text Figure 2. The lower two panels are inferences for the N-lim replicates while the upper two panels show previously published DFE estimates for the two C-lim replicates. It should be noted that the y-axes is log-scaled to enable visualizing the shape of the high-fitness nose of the DFE which is critical in determining the dynamics (see Section 2).

**Figure 3:**
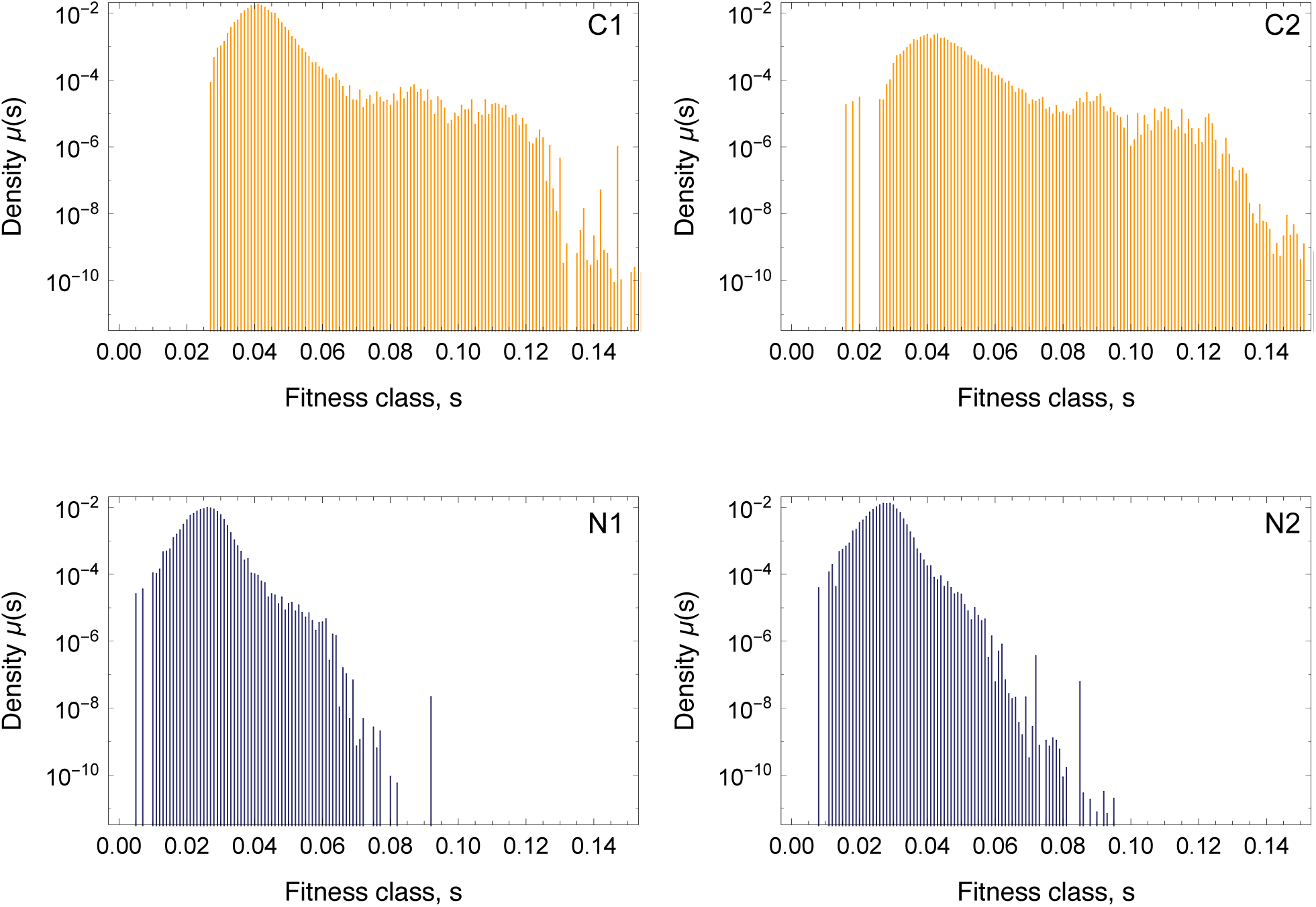
The distribution of fitness effects of all detected single beneficial mutations arising in each of the four replicates. Rates to each fitness effect are inferred by counting the number of cells in a given fitness range over time and using a deterministic approximation to infer *μ*(*s*) (see [1]).

### 2 The DFE and population size determine early lineage- and genotype-size statistics

Here we outline how the DFE influences the size statistics of single-mutants and, subsequently, multiple-mutants and how these together with neutral drift affect lineage sizes through time. We show that single-mutants determine the diversity over short timescales but multiple-mutants determine the diversity over longer times scales. We outline that the entropy of all adaptive genotypes in the population undergoes an expansion followed by a crash. This expansion and crash can be measured by tracking the entropy of adaptive lineages.

#### 2.1 Distribution of single-mutant sizes over time and effective *U_b_*

The arguments in this section follow similar arguments presented in the supplemental methods of [1] and [3]. Consider a mutation at a given site in the genome *i* with fitness advantage *s*, which occurs at a constant rate *R_i_* = *NU_i_* from a pool of *N* ancestral feeding cells. The total number of single-mutant cells with this mutation is the convolution of the single mutant size distribution (see [1]) with the distribution of times at which the mutations enter (uniform). This yields a distribution of rescaled clone sizes *ν* = *n*/*ñ* = *n*/((*c*/*s*)*e^st^*):

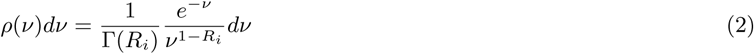

The rate of mutating a given site is *μ* ≈ 3 × 10^−10^ per bp per generation [1] hence *R_i_* ≪ 1 and the distribution of sizes of unique SNPs is approximately *NUe^−ν^*/*ν* with the largest single-mutants being those that arose immediately reaching sizes *n* ∼ (*c*/*s*)*e^st^* and the smallest being those that arose in the previous generation and that are of size ∼ 1.

In general there are a range of possible *s* with different mutation rates *μ*(*s*) hence the distribution of single-mutant abundances is obtained by summing all of these:

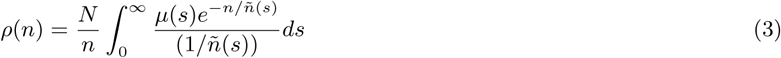

Thus the exact form of the single-mutant distribution will depend on the shape of *μ*(*s*) and in general will not retain the characteristic 1/*n* form even at low frequencies unless *μ*(*s*) is sharply peaked over a narrow range.

##### Note on clonal interference

The above expression assumes no competition between lineages. This is a good approximation at early times, but for times after the single-mutant class has reached a significant fraction of the entire population clonal interference cannot be ignored. In the simplest model however, clonal interference — which results from the total population size of cells having to remain ≈ *N* — is captured by considering a genotype’s fitness advantage over the (increasing) mean-fitness of the population, *x̄*(*t*). Thus, while the previous expression will still accurately capture relative frequencies of mutants, the absolute numbers of the different genotypes will be modified by a factor

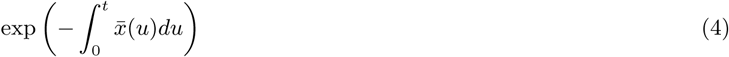

Consider now an entire class of mutations with a given fitness effect range *s*, *s* + *ds* rather than a specific mutation. The mutation rates to this range of fitness effects is given by *μ*(*s*)*ds*. Provided the population size *Nμ*(*s*)*ds* ≫ 1, (true for the majority of the range we consider, but see below for when this does not hold) single mutants occur in large numbers for a given fitness range (since *Nμ*(*s*)*ds* = *R* ≫ 1) and therefore, as a class, behave quasi-deterministically. The fraction of single-mutant cells in a given fitness range *f*(*s*)*ds* is simply

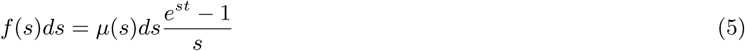

Since the timescale over which mutations with fitness effects *s* will contribute is 1/*s* and the number establishing per generation is *Nμ*(*s*)*ds*, the number of mutations contributing significantly to the genetic diversity in each fitness class is of order *Nμ*(*s*)*ds*. The small lineage size ensures that the majority of single-mutants will occur inside independent lineages, hence the number of lineages that expand at a rate between *s*, *s* + *ds* is also *Nμ*(*s*)*ds*.

##### Effective fitness effects and effective mutation rates

For times *st* > 1, the differences in fitness effects between mutations enable some lineages to expand more relative to others. The range of fitness effects dominating all adaptive cells are those for whom *f*(*s*)*ds* = *μ*(*s*)*ds*(*e^st^* – 1)/*s* is maximized, which corresponds to fitness classes in the region of *s̃*, where

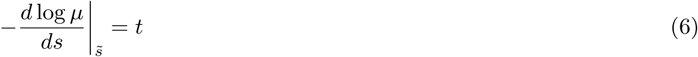

the region having a width *δs̃* is given by 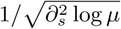 evaluated at *s̃*. Mutations in the fitness range [*s̃*(*t*), *s̃*(*t*) + *δs̃*(*t*)] dominate the population at *t* and hence they can be considered an effective *s*. Similarly, the rate of mutation to these fitness effects *Ũ_b_* = *μ*(*s̃*(*t*))*δs̃*(*t*) can be considered an effective mutation rate. Together these can be used to determine the number of expanding unique single-mutants contributing to the diversity at time *t*:

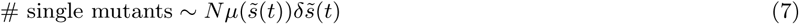

##### Stochastic occurrence of highly fit single mutants

The fitness of the predominant fitness class *s̃* typically increases over time because the size of each fitness class has an exponential weighting *e^st^*. Since *μ*(*s*) in both environments we analyze decreases with increasing *s*, this results in a smaller *Ũ_b_* at later times. This has important consequences for stochasticity. The single mutant genetic diversity is determined by an exponential expansion of types, but where the number of types is ever decreasing (see “top-hat” DFE example). For late enough times, the predominant fitness class *s̃*(*t*) with associated width *δs̃*(*t*) has an associated mutation rate *Ũ_b_*(*t*) = *μ*(*s̃*(*t*))*δs̃*(*t*), which can be small enough that *NŨ_b_*(*t*) ≲ 1. At this point, the number of mutations contributing to the adaptive single-mutant population becomes small since the population is dominated by the small number of high fitness clones. This results in lower genetic diversity and increased stochasticity. At times *t̃* ≈ (1/*s̃*)In (*s̃*/*Ũ_b_*) single-mutants comprise a significant fraction of all cells. Around this time they begin to interfere with one another, slowing their exponential expansion. The predictions resulting from considering a “predominant” class are checked in the section exploring the dynamics that result from a “top-hat” DFE.

#### 2.2 Distribution of multiple-mutant sizes over time

Here we calculate the distribution of sizes of multiple-mutant clones, which are fed by mutation from exponentially growing single mutant clones. We note that a closely related calculations, for neutral mutations with possibly large mutation rate, has been performed in [4]). Consider a single-mutant (or mutant-class, or indeed a multiple-mutant) that, early in its trajectory is expanding in the population as

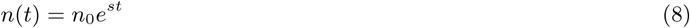

(if this were an class of single mutants fed from the ancestor at rate *U* then *n*_0_ ≈ *NU*). Suppose that second mutants with fitness advantage *s* × Δ occur at rate *U*′. If fitness effects combine additivity, the fitness of the second-mutant relative to the ancestor is *s*(1 + Δ). Provided *n*_0_*U*′ = *NUU*′/*s* ≪ 1 the occurrence of double mutants will be rare and thus highly stochastic, with most of the stochasticity coming from the occurrence time of the mutant, rather than drift. The expected time for the ith double-mutant to establish is *t_i_* where

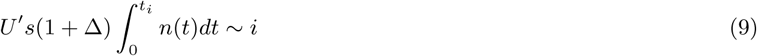

since this mutant will grow as *n′* ≈ (*ν*/*s*(1 + Δ))*e*^*st*(1+Δ)^ (with *ν* a random variate from an exponential distribution with mean 1 stemming from stochasticity of drift), the typical size 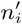 of the *i*th double mutant is

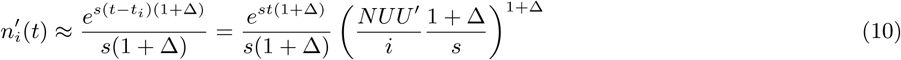

The typical relative sizes of subsequent mutations therefore follows the series

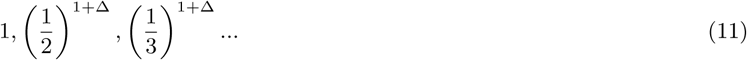

hence the number of terms that contribute significantly to this series (and thus the typical number of mutations that contribute significantly to the total number of double-mutant cells) is controlled by the ratio of the fitness effects of double mutants compared to single mutants Δ. For Δ ≳ 1 one can consider the sum of this series (evaluated via an integral) to estimate of the number of terms (i.e. mutations) contributing to the sum and is found to be

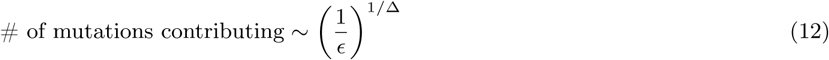

where 1 – *ϵ* controls how much of the sum one accounts for (i.e. an *ϵ* = 0.1 corresponds to the resulting number of double-mutants accounting for ~ 90% of the population). The result of which is that the number of terms contributing to the total population rapidly approaches 1 as Δ increases and even for small Δ = 1 is a modest, meaning only a handful of double mutants dominate the population. The mean and standard deviation in the number of double-mutants comprising ≥ 90% of the total population of double-mutants, as a function of increasing Δ is shown in Table 2. We note that as Δ → 0 the number of terms contributing to the sum diverges, hence the dominance of a handful of clones is a feature of beneficial mutations i.e. those with a significant fitness advantage over the feeding population and thus a Δ which is not very close to zero.

**Table 2:**
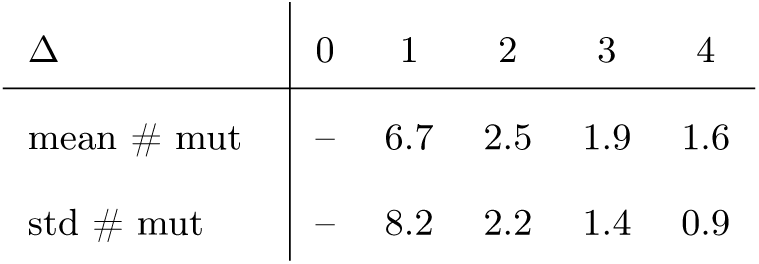
The mean and standard deviation of the number of second mutants that will comprise ≥ 90% of the total doublemutant population as a function of the growth advantage of the double mutant (Δ). In case of a neutral mutation Δ = 0 the number of mutations contributing to the total diverges over time, never reaching a fixed number as newer mutations keep contributing.

| $\Delta$ | 0 | 1 | 2 | 3 | 4 |
| --- | --- | --- | --- | --- | --- |
| mean # mut | – | 6.7 | 2.5 | 1.9 | 1.6 |
| std # mut | – | 8.2 | 2.2 | 1.4 | 0.9 |

For Δ > 1 the number of mutants that contribute is essentially only the first few and thus there is a rather general prediction that that fit double mutants should always cause a diversity crash independent of the details of the shape of the DFE, though the extent and timing of the crash depends on the DFE. The previous argument is been based on “typical” sizes. We note that the exact timing of the mutant is stochastic and hence there are fluctuations around these typical values for all of the *i*. The only fluctuations that really matter though are for the first few mutants *i* = 1, 2, 3…, the first being particularly important, which can be seen by considering sub-integer *i* in the above expressions. *i* = 0.1 will occur 10% of the time *i* = 0.01 occurs 1% of the time. These result in double-mutants that are 10^1+Δ^ or 100^1+Δ^ times larger. For large Δ, this can produce a double-mutant (the first one that occurs) that dominates the entire population, almost completely wiping out genetic diversity. Hence, the relatively rare events with an early and fit double mutant are predicted to be able to wipe out diversity.

We can use similar arguments — i.e. using the probability of when the first established second-mutant will enter the population ‗ to predict the distribution of second-mutant sizes (as for single-mutants in the previous section). We find the distribution over sizes of the earliest second-mutant to occur is

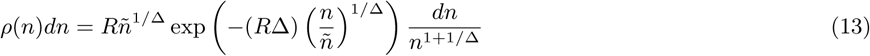

where *ñ* = (*c*/(1 + Δ)*s*)*e*^*s*(1+Δ)*t*^. This expression holds out to a size *n* ≈ *ñ* beyond which it develops exponential tail (mutations above this size occurred immediately and their distribution of sizes above this result from the stochasticity associated with genetic drift). As a sanity check, this expression recovers the classic Luria-Delbruck form of ~ 1/*n*^2^ for neutral mutations (Δ = 0) and approaches 1 ~ /*n* form for Δ ≫ 1 which is the same as if the feeding population were constant and feeding a rare beneficial mutation. The discussion around Eq. 7 in the text above (regarding clonal interference and its effect on the absolute sizes of single-mutants) applies similarly to double-mutants i.e. the above expression will capture the scaling relation between relative frequencies of double-mutants but the absolute size of each in the population will be modified by the changing mean fitness term.

Below we explore two concrete examples of DFEs to highlight how the shape of the DFE affects mutation statistics and diversity.

#### 2.3 Example: a delta-function DFE

##### Single mutants

Consider that the DFE is very sharply peaked such that every mutation confers the same advantage, *s*, and that mutations to this fitness occur at rate *U_b_*. We begin by restricting analysis to single mutations only (we will later ask when this assumption breaks down to understand the effect of multiple-mutants). At early times, the mean fitness will be negligible and the probability of a mutation establishing in the population in the interval (*t*′, *t*′ + *dt*′) is simply

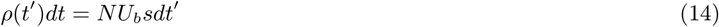

meaning that mutants enter at a constant rate. Early on, each of these mutants grows exponentially

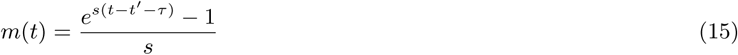

where *t*′ is the time of occurrence of the mutation and *τ* is a stochastic variable called the “establishment time” which accounts for differences in the time it takes for a mutation to climb out of the stochastic behavior that dominates at low frequencies and begin to grow exponentially. (The distribution of *τ* is peaked at zero with variations of ±1/*s* either side). Because mutants grow exponentially and come in at a constant rate, the total number of single-mutant cells is dominated by the earliest few mutations that occur. More precisely, because the growth rate of mutants is *s*, only mutants that enter in the first ~ 1/*s* generations contribute significantly to the expansion and hence the total number of single-mutant cells increases

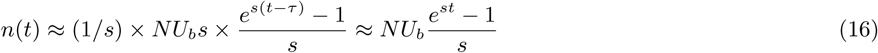

This enables us to estimate how the mean fitness of the population increases:

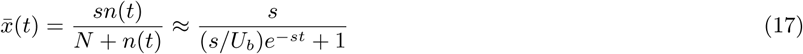

This correctly captures the behavior that the mean fitness, *x̄*(*t*) is small for *t* ≪ (1/*s*)In(*s*/*U_b_*) then asymptotes to *x̄*(*t*) ≈ *s* as *t* ≫ (1/*s*)In(*s*/*U_b_*) (Figure 4 a & b, 2nd row). The timescale for the mean fitness to increase appreciably

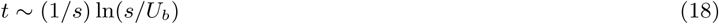

is particularly important in understanding the diversity. The feeding of new mutants remains roughly constant until *t* ≈ (1/*s*)In(*s*/*U_b_*) at which time time the feeding population declines rapidly and the fitness advantage of new mutations is diminished, causing the rate of establishing new mutations to decline and eventually stop altogether. These observations can be used to calculate the entropy

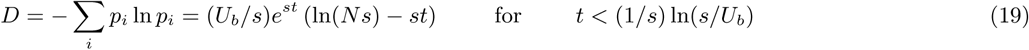

**Figure 4:**
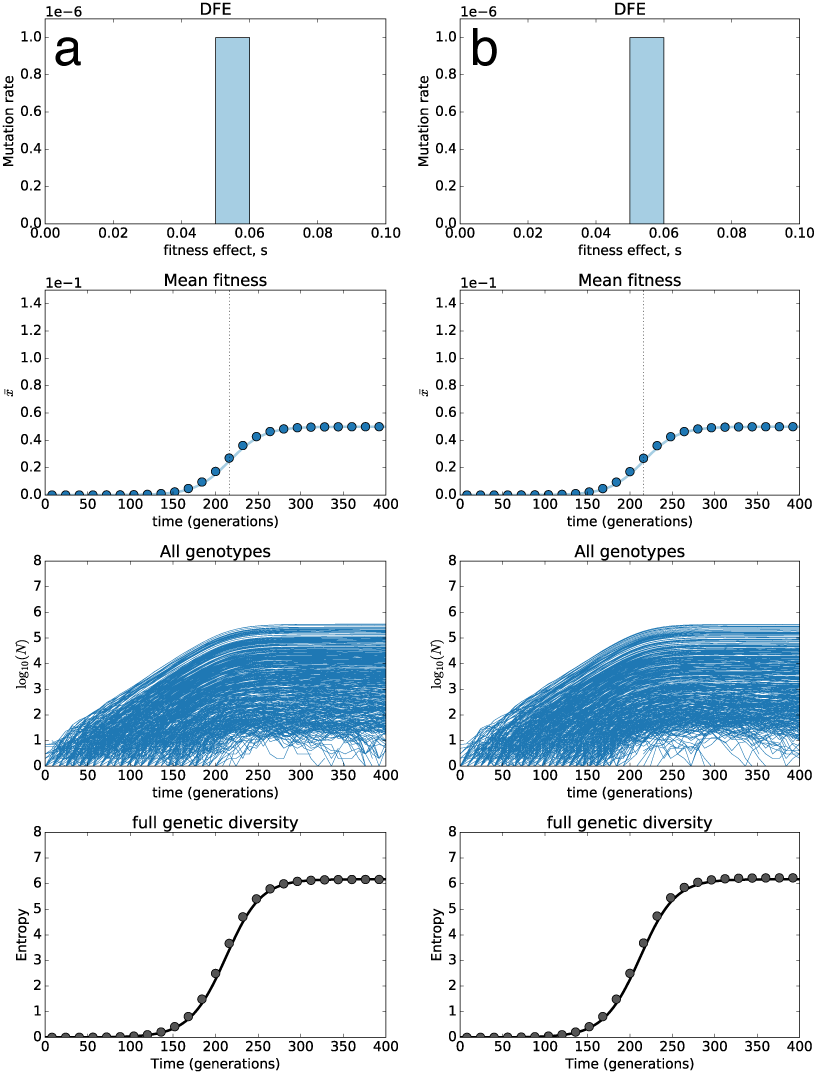
Diversity dynamics for a delta-function DFE with only single-mutants. Two replicate simulations (a, b) using a single-mutant model with a delta-function DFE. Top row, the DFE, *ρ*(*s*) = *δ*(*s* – *s*_0_) where *s*_0_ = 0.05 and *U_b_* = 10^−6^ the bottleneck population size is 5 × 10^7^ which grows by a factor of 256 between bottlenecks resulting in an effective population size of ≈ 4 × 10^8^. Second row, the mean fitness plateaus over a timescale of the “sweep” time (1/*s*)ln(*s*/*U_b_*) ≈ 216 for the numbers in this simulation, and highlighted by the dashed vertical gridline. Third row, the genotype trajectories of all unique genotypes that arise in the simulation. Bottom row: the entropy trajectory (see Section 5.2) of all genotypes in the population (data points) compared to the theoretical prediction from using Eq. 22 in the text.

The entropy will plateau at *t* ~ (1/*s*)ln(*s*/*U_b_*) obtaining its maximum diversity

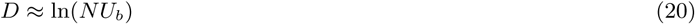

These features are indeed born out in simulations (Figure 4 a & b 4th row). Note that a slightly more accurate form for the entropy can be obtained by accounting for the diminishing fitness advantage of clones as they expand rather than using a sharp cut-off. This more accurate approximation shows good quantitative agreement with the measured entropy of the distribution.

##### Multiple mutants

How do the previous observations change if we allow multiple mutants to enter? In this delta-function DFE model multiple mutants allow genotypes to have different fitnesses, which will typically reduce diversity as some are fitter than others. At the same time, however, there is a far larger space of genotypes available to multiple mutations which will act to increase diversity. We start by considering double-mutants that have a fitness advantage *s*_1_ + *s*_2_. These are fed from the expanding class of 1st mutants which is increasing as

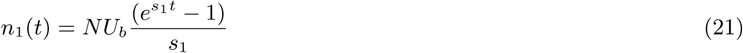

The probability that a mutation, destined to establish, enters the population in the interval (*t*′,*t*′ + *dt*′) is now

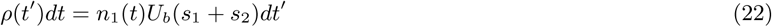

which is different to before since *n*_1_(*t*) is increasing exponentially. Each of the mutants therefore grows exponentially as

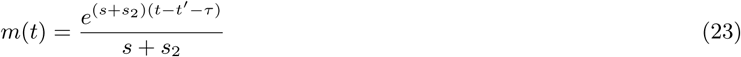

Provided *n*_1_(*t* = 1/*s*_1_)*U_b_* ≪ 1 as is the case in these simulations and for the multiple mutants in the experiments, the single mutants do not feed double mutants immediately, but must grow up exponentially for an appreciable fraction of the sweep time first. Considering the typical time the second mutants will enter, the size of the expanding double-mutant class will be

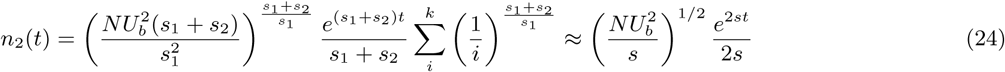

where *i* refers to the *i*th double-mutant that enters and *k* is the index of the most recent double mutant to occur. This is very different if *n*_1_(*t* = 1/*s*_1_)*U_b_* ≫ 1, in which case many double-mutants occur before the single mutant class grows exponentially and hence the expansion is dominated by the 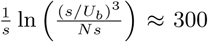 double-mutants that occur in the first 1/(*s*_1_ + *s*_2_) generations, leading to the double-mutant class expanding as

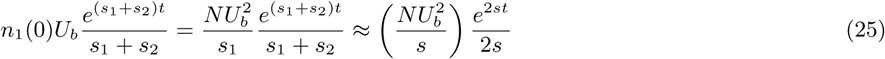

which, if 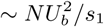 is much larger than in Eq. 24. If 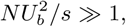 the entire class of second mutants is dominated by the first few mutations to occur since

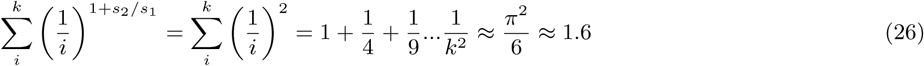

The fact that the diversity of second-mutant is dominated by the first handful is also borne out in simulated data (Figure 5C, red lines). Only these double-mutants will contribute significantly to the entropy, and since there are already *NU_b_* ≫ 1 single-mutants expanding and driving the early-time entropy, the handful of double-mutants does not significantly increase it. However, the double mutants do expand and eventually outcompete the single mutants. This effect causes a diversity “crash” resulting in a significant reduction in entropy (Figure 5, bottom row). The magnitude and timing of this drop in diversity is determined by the relative diversities of the single versus double-mutant classes and the timing of when the double mutants out-compete the single mutants i.e. when *n*_2_(*t*) ≫ *n*_1_(*t*) which yields

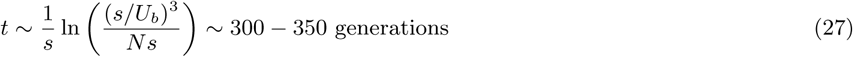

**Figure 5:**
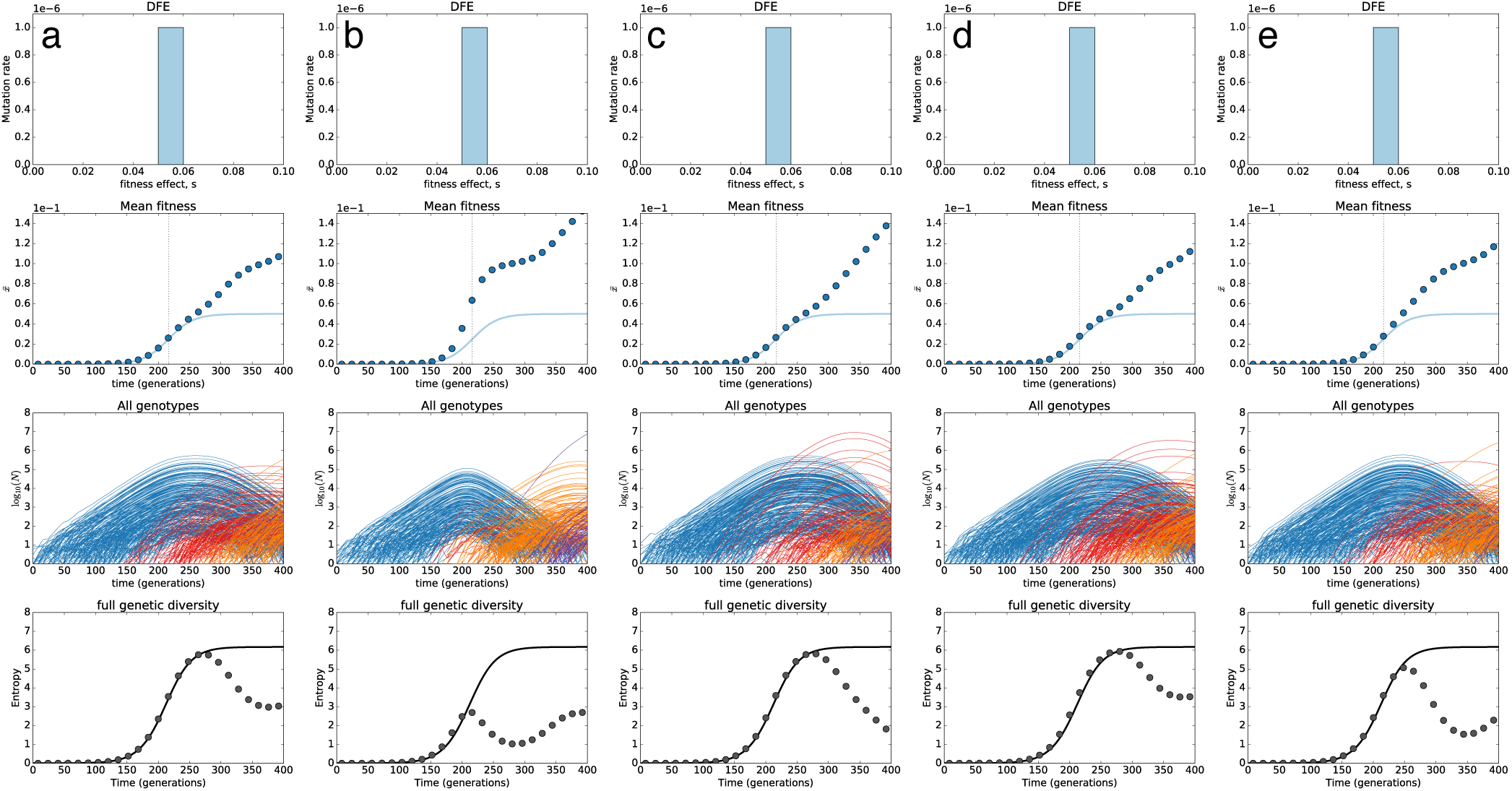
Diversity dynamics for a delta-function DFE with multiple-mutants that combine additively. Top row, the DFE, *ρ*(*s*) = *δ*(*s* – *s*_0_) where *s*_0_ = 0.05 and *U_b_* = 10^−6^ the bottleneck population size is 5 × 10^7^ which grows by a factor of 256 between bottlenecks resulting in an effective population size of ≈ 4 × 10^8^. Second row, multiple mutants increase the mean fitness significantly when they comprise an appreciable fraction of the population. The double-mutants will be a significant fraction of the single mutants after a time 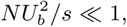 generations leading to large deviations from the single mutant predictions (solid blue line, second row). Third row, the genotype trajectories of all unique genotypes that arise in the simulation (single-mutants blue, double-mutants red, triple-mutants orange, quadruple-mutants purple). Note the emergence of a small number of very large double mutants (a, c, d, e) and occasionally of a anomalously early triple mutants (b). Bottom row: the entropy trajectory (see Section 5.2) of all genotypes in the population (data points) compared to the theoretical prediction from using Eq. 22 in the text that assumes single mutants only. The expansion of the double-mutants causes the diversity to “crash” over a similar timescale as when the double mutants comprise a significant fraction of the population. The small number of large double-mutant clones eventually outcompete the single mutants over this timescale.

At this time, the single mutants have been outcompeted and are therefore at a total abundance < 0.5, hence their contribution to the entropy is reduced from ~ In(*NU_b_*) at the maximum (Figure 5D where In(*NU_b_*) ≈ 6), to ~ 1. The contribution of the second-mutants to the entropy will also be *O*(1) and should largely be independent of *N* provided 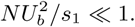

##### Stochasticity

While Eq. 24 correctly captures the typical size of the ith mutant, there are fluctuations due to variation in the occurrence time of the first few double-mutants. This stochasticity can be seen in Figure 5a–e showing 5 independent simulated evolutions with the same DFE.

#### 2.4 Example: a “top-hat” DFE

##### Single mutants

Consider a “top-hat” distribution with density *μ*(*s*)*ds* = 2 × 10^−5^*ds*, such that the total beneficial mutation rate is again *U_b_* = 10^−6^ but it is now uniformly spread out over the interval [0.025, 0.075]. In the deterministic approximation, which holds well for single-mutants, the total number of cells across time binned by fitness will be

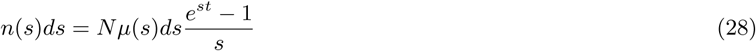

The mean fitness over time is therefore

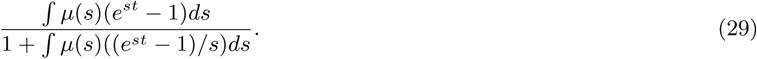

Typically, only a narrow range of fitness classes contribute to the total number of adaptive cells and hence one can normally find an effective fitness effect *s̃* and range of fitness effects *δs̃* and an effective rate to these fitness effects *U_b_*. For the case of the “top-hat” these are straightforward to work out: fitness classes grow exponentially relative to one another so only a range of fitnesses within 1/*t* of the maximum fitness effect, *s_max_* will contribute. Hence the effective selection coefficients will lie in the range

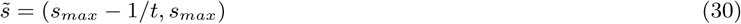

and since the rate to fitness effects is constant, the effective beneficial mutation rate becomes

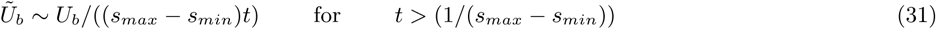

Substituting *s̃* and *Ũ* into the expressions for the mean fitness and entropy from the delta-function distribution successfully predicts the behavior of both the mean fitness over time (Figure 6 a & b, 2nd row) as well as the genetic diversity (Figure 6 a and b, 4th row).

**Figure 6:**
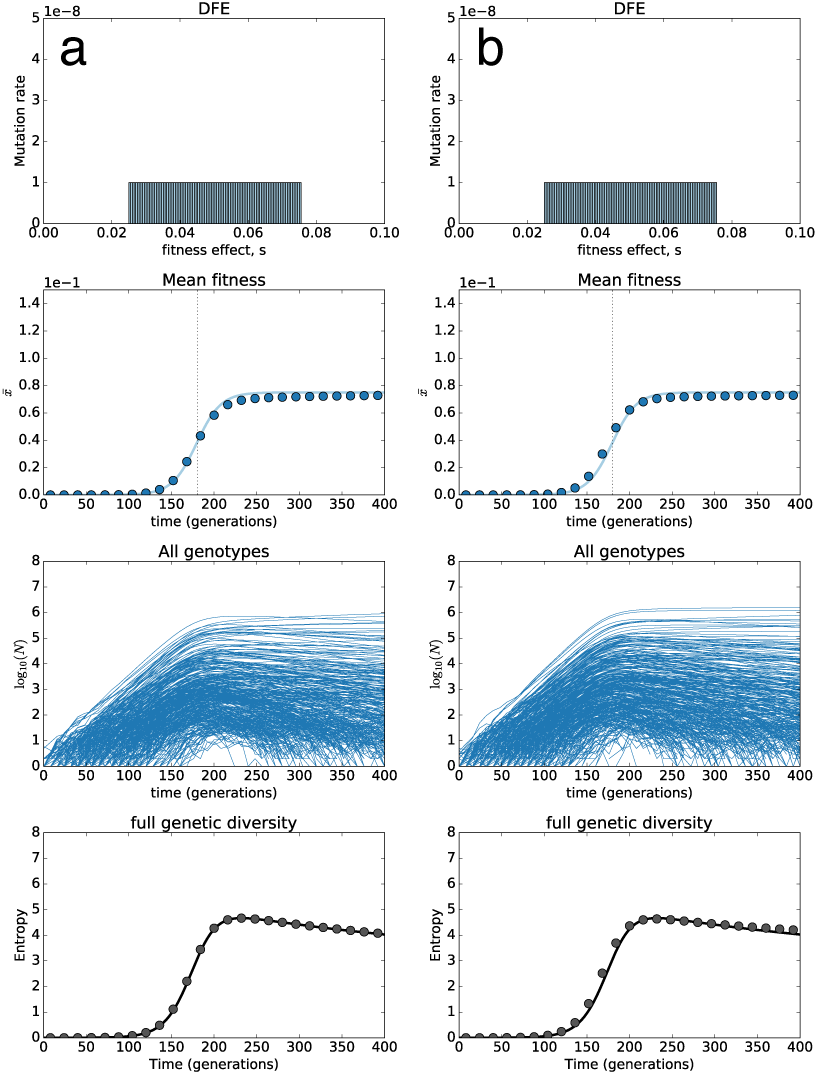
Diversity dynamics for a top-hat DFE allowing only single mutants. Simulations using a single-mutant model with a top-hat DFE. Top row, the DFE, *ρ*(*s*) = 2 × 10^−5^ between *s* = 0.025 and *s* = 0.075 and *U_b_* = 10^−6^ the bottleneck population size is 5 × 10^7^ which grows by a factor of 256 between bottlenecks resulting in an effective population size of ≈ 4 × 10^8^. Second row, the mean fitness plateaus over a timescale of the “sweep” time (1/*s*)In(*s*/*U_b_*) ≈ 170 since the “predominant” *s* = 0.075. Third row, the genotype trajectories of all unique genotypes that arise in the simulation. Bottom row: the entropy trajectory (see Section 5.2) of all genotypes in the population (data points) compared to the theoretical prediction from using Eq. 35 in the text. The slow decline of the diversity is due to clonal competition between single-mutants with different fitness effects.

The effect of changing the DFE to a top-hat distribution with the same mean fitness effect as the delta function results in the mean fitness increasing more rapidly because the increased effective fitness effect *s̃* has a larger effect the rate of mean fitness increase than the reduced effective mutation rate *Ũ_b_* does. This results from the timescale for the mean to increase depending inversely on the fitness effect and only logarithmically on the mutation rate via Eq. 18. Similarly, the genetic diversity increases more rapidly as it is being driven by larger fitness effect mutants that expand faster. However it peaks at a smaller value because the number of unique mutants contributing to the expansion is smaller (and decreases over time) due to the lower rate of mutating to the specific “predominant” fitness class *s̃*. This effect produces decline in the genetic diversity, even in the absence of further mutations, due to the larger fitness effect mutants out-competing the others. After the initial exponential expansion, the entropy at time *t* is effectively given by In(*NŨ_b_*) which in this case decreases as – ln(*t*) due to *Ũ_b_* ~ 1/*t*.

##### Multiple mutants

The effect of multiple mutants is qualitatively similar for a top-hat DFE as it is for the delta-function DFE. The emergence of a handful of large double mutants causes the entropy trajectory to crash at roughly the time when double mutants dominate the population. This subsequently partially recovers with the emergence of triple- and quadruple-mutants. The stochasticity in the size distribution of double-mutants resulting from exponential feeding is clear in Figure 7 a-e which are 5 replicate simulations. Panels a-c show more typical simulations where there are a handful of large double-mutants. Panels d-e show examples of anomalously early multiple mutants that drive an even more dramatic diversity crash.

**Figure 7:**
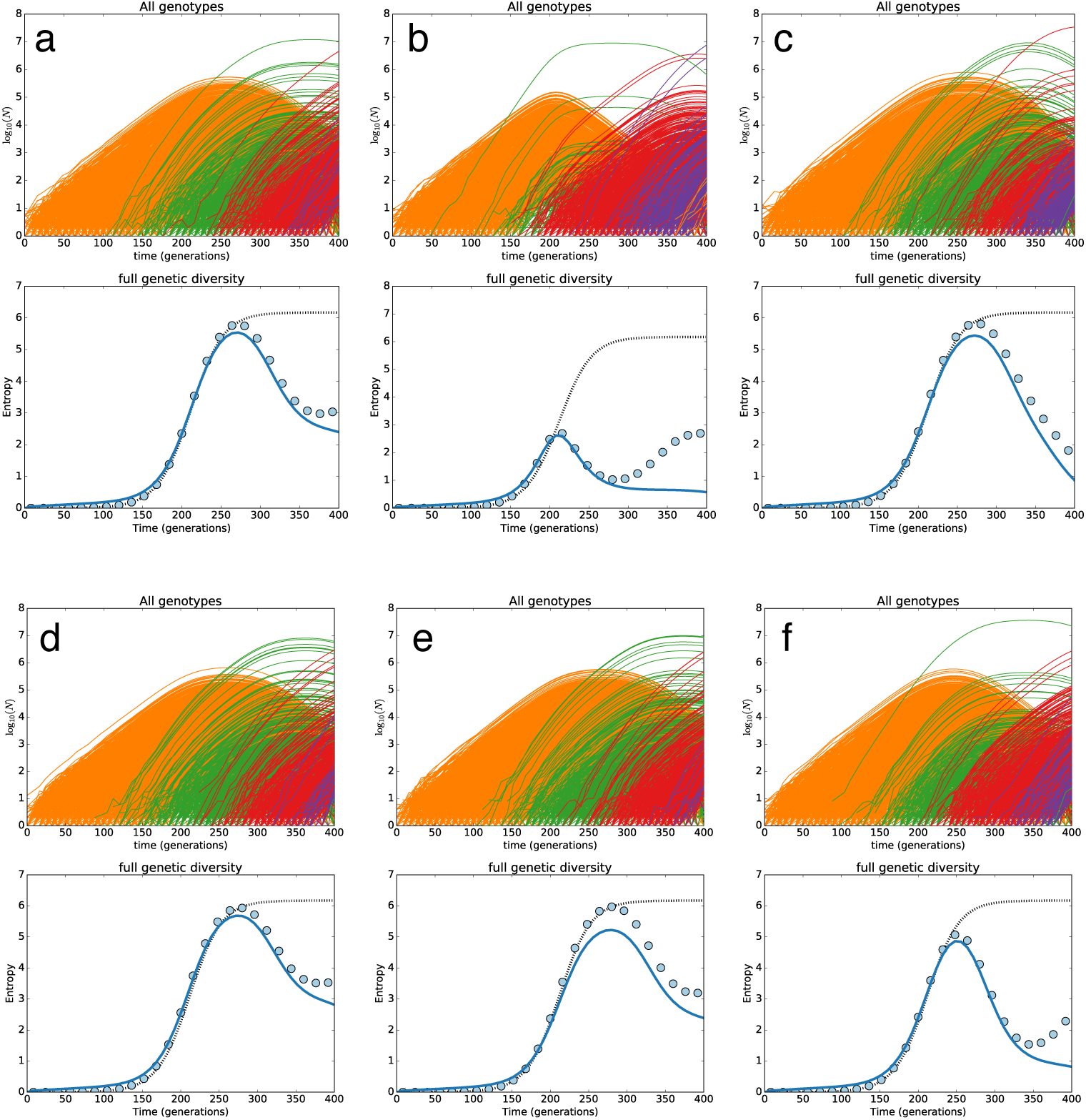
Diversity dynamics for a top-hat DFE with multiple-mutants that combine additively. Top row, the DFE, *ρ*(*s*) = 2 × 10^−5^ between *s* = 0.025 and *s* = 0.075 and *U_b_* = 10^−6^ the bottleneck population size is 5 × 10^7^ which grows by a factor of 256 between bottlenecks resulting in an effective population size of ≈ 4 × 10^8^. Second row, multiple mutants increase the mean fitness significantly when they comprise an appreciable fraction of the population. The double-mutants will be a significant fraction of the single mutants after a time 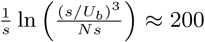 (with predominant *s* = 0.075) generations leading to large deviations from the single mutant predictions (solid blue line, second row). Third row, the genotype trajectories of all unique genotypes that arise in the simulation (single-mutants blue, double-mutants red, triple-mutants orange, quadruple-mutants purple). Note the emergence of a small number of very large double-mutants (b, e) and occasionally of a anomalously early triple mutants (a, d). Bottom row: the entropy trajectory (see Section 5.2) of all genotypes in the population (data points) compared to the theoretical prediction from using Eq. 35 in the text that assumes single mutants only. The expansion of the double-mutants causes the diversity to “crash” over a similar timescale as when the double mutants comprise a significant fraction of the population. The small number of large double-mutant clones eventually outcompete the single mutants over this timescale.

#### 2.5 Barcode lineage diversity tracks genotype diversity into the crash

In the previous sections, genetic diversity is measured via the entropy of all *unique genotypes* in the population. We however can only measure the abundance of *unique lineages.* A natural question is then: **does the adaptive lineage diversity (measured via entropy of all adaptive lineages) match the adaptive genetic diversity (measured via entropy of all adaptive genotypes)?**

Somewhat surprisingly, our simulations show that the diversity of adaptive barcodes tracks true genotypic diversity well beyond the emergence of single mutations, robustly capturing the diversity “crash” caused by double-mutants outcompeting single-mutants (Figure 8). Top panels plot the log-abundance of almost all genotypes that enter the population colored by the number of adaptive mutations: single mutants (orange), double mutants (green), triple mutants (red), quadruple mutants (purple). The bottom panel plots the entropy of all adaptive genotypes (data points) and, for comparison, the predicted entropy of single mutants (hashed line) and entropy of all adaptive barcode lineages (blue line). In all simulations performed the diversity of adaptive barcode lineage closely tracks the true genetic diversity of adaptive mutations out to the point of the minimum diversity (the “crash”), which in all cases is caused by the expansion of a small number of double-mutants.

**Figure 8:**
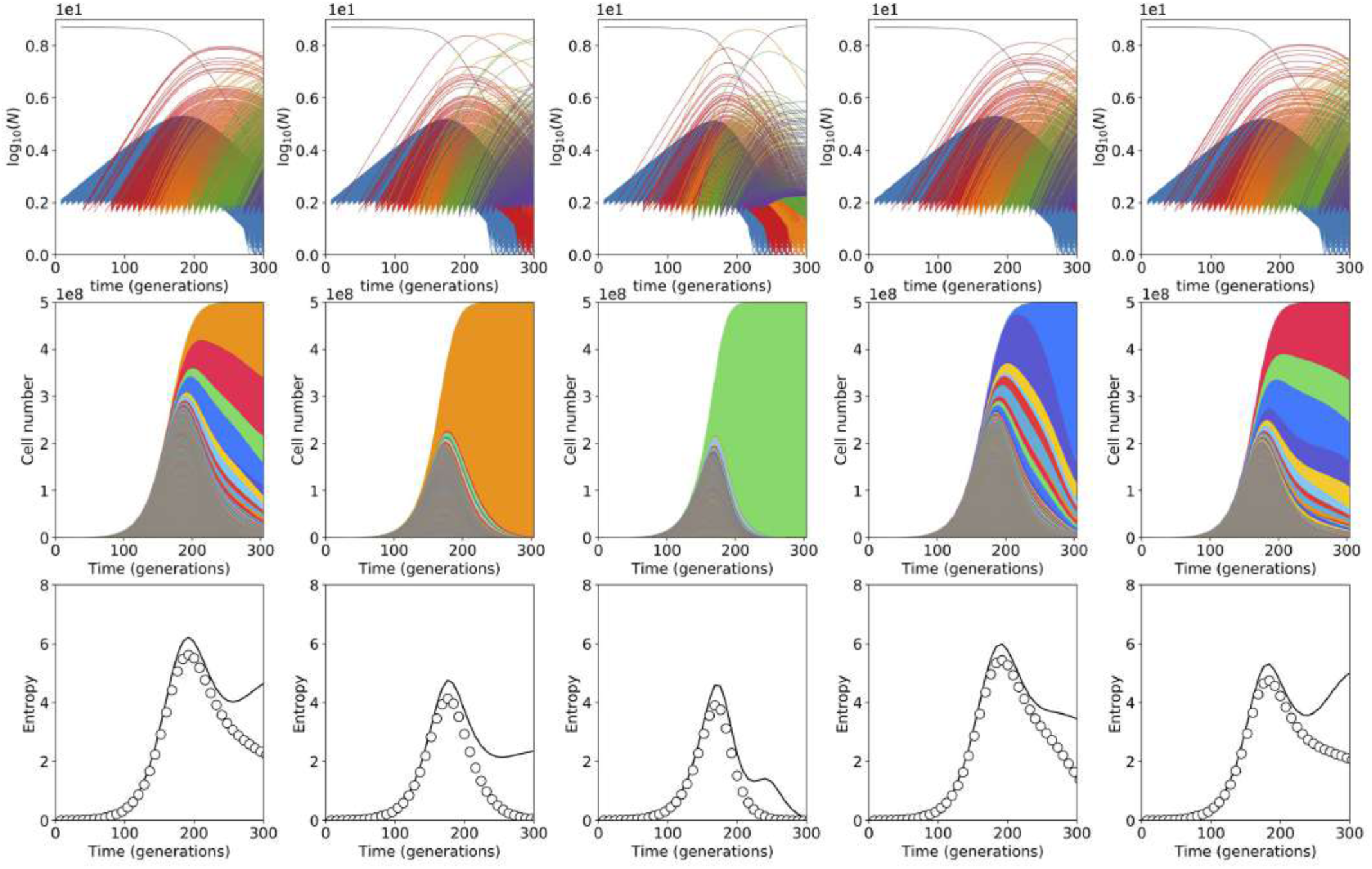
Barcode diversity tracks true diversity. (a-f) stochastic simulations of the evolutionary dynamics that result from a delta function DFE peaked at *s* = 5%, with *U_b_* = 10^−6^ and *N* = 4 × 10^8^. Top panel lines are all genotypes in the population through time colored by the number of beneficial mutations harbored by that genotype: single-mutants (orange), double-mutants (green), triple-mutants (red) and quadruple mutants (purple). As predicted, double-mutants are typically dominated by between 1 and 10 genotypes, in contrast to single-mutants in which hundreds of distinct genotypes contribute. Bottom panels show the entropy of all beneficial genotypes calculated from the true genetic diversity (data points), predicted assuming single-mutants only (hashed line) and inferred using the abundance of adaptive barcoded lineages (blue lines). Barcode abundance can be used to robustly infer true beneficial diversity out to when double mutants expand to cause the diversity “crashΔ.

The reason adaptive lineages provide useful information about double-mutant expansions is because only a handful of double-mutants contribute significantly to diversity and these can occur inside ~ 10^3^ different adaptive barcode lineages containing an expanding single-mutant. It is therefore likely that all of the large double mutants occur in different barcode lineages and, because they occur anomalously early, quickly become the dominant types in those lineages. Tracking the abundance of the largest adaptive barcode lineages at late times therefore provides a robust method to infer the diversity of adaptive mutations in a population and can be used to test whether double-mutants cause a diversity “crash” that is predicted in both theory and simulation.

Extending the above arguments explains why barcode diversity does not track true genetic diversity for triple mutants. It is predicted that ~ 10 triple-mutants contribute significantly to diversity. However, these are all likely to occur in the largest few double mutant lineages, meaning these largest few lineages will harbor “sub-clonal” diversity. Adaptive barcode diversity will therefore underestimate true adaptive diversity beyond the point of minimum diversity causing the diversity trajectories to diverge (Figure 8).

#### 2.6 Growth-inhibiting drug DFE

Here we model how adaptive genetic diversity evolves under a DFE with a small supply of large-effect mutations, as might occur in the presence of a growth-inhibiting drug. As an example of what happens in such a scenario, we use a model DFE where small-fitness-effect mutations (*s_α_* = 0.05) occur at a relatively high rate (*U_α_* = 10^−5^ per cell division) while large fitness-effect mutations (*s_β_* = 0.20) occur at a lower rate (in the range 10^−11^ < *U_β_* < 10^−9^ per cell division). In this context we take “large” fitness effects to mean that *s_β_* > *s_α_* × (ln(*Ns_α_*)/ln(*s_α_*/*U_α_*)). That is, large-effect mutations destined to establish will typically sweep regardless of the fitness of the genotype on which they fall (since the fitness effect is larger than the range of fitnesses expected in the population at any time). In this limit the the dynamics of adaptive genetic diversity depends most sensitively on the supply of large-effect mutations *U_β_*.

Large effect mutants enter typically every ~ 1/*NU_β_s_β_* generations. The early time dynamics of the modest-effect mutations studied here takes place over timescales *τ_sw_* = (1/*s_α_*)In(*s_α_*/*U_α_*) called the “sweep time” (~ 170 generationsfor the parameters above and those in Figures 9 - 11). It is the relative magnitudes of these timescales that can be used to assess the impact of the large-effect mutants on the adaptive genetic diversity.

**Figure 9:**
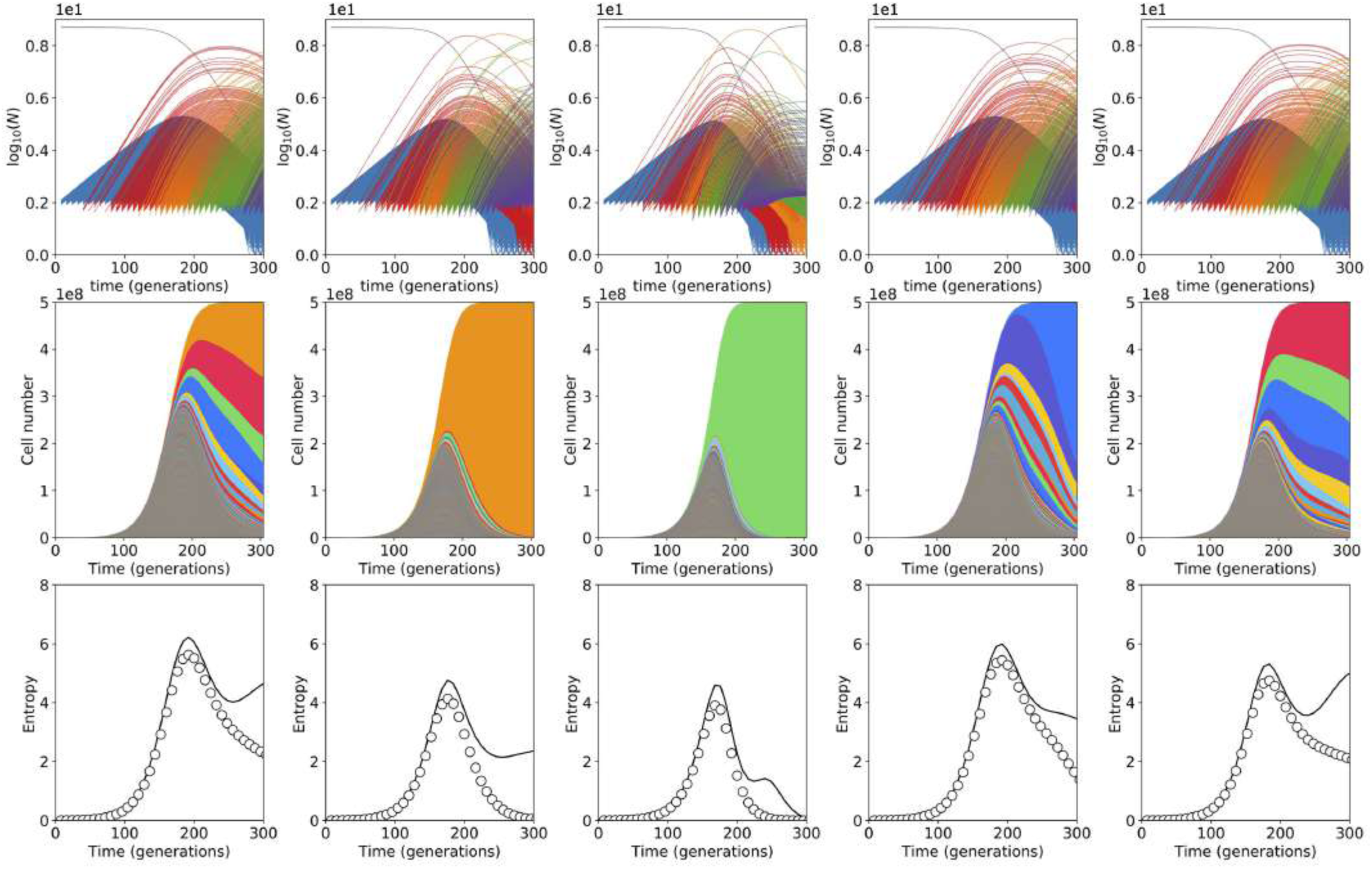
Diversity dynamics for DFE modeling growth-inhibiting drug. Here the DFE is composed of two delta-functions (at *s_α_* = 0.05 with rate *U_α_* = 10^−5^ and *s_β_* = 0.20 with rate *U_β_* = 10^−11^). Top row, the genotype trajectories of all unique genotypes that arise in the simulation (single-mutants blue, double-mutants red, triple-mutants orange, quadruple-mutants purple). Large crashes in diversity (column 2 and 3) are caused by anomalously early double mutants comprised of two small-effect mutations. Bottom row: the entropy trajectory (see Section 5.2) of all genotypes in the population (data points) compared to the theoretical prediction from using Eq. 35 in the text that assumes single mutants only.

When the supply of large-effect mutations is low enough, large-effect mutations have little impact on the early adaptive diversity dynamics (Figure 9, where *U_β_* = 10^−11^). In this case the typical time it takes for a large-effect mutation to enter is ~ 10^3^ generations and hence the probability of large-effect mutation entering during the early time dynamics (<170 generations) is small. Thus, when large diversity crash does occur, it is driven by anomalously early double mutants as can be seen in Figure 9 and as is the case for the empirical DFEs observed in the experimental evolutions.

As one increases the supply of large-effect mutations (Figure 10, where *U_β_* = 10^−10^) the timing of the first large-effect mutation and the sweep time become comparable. For parameters used here, this is predicted to occur at *U_β_* ~ 10^−10^, and Figure 10 indeed shows that, in a substantial fraction of simulations, a large-effect mutation enters and causes a single-mutant driven diversity crash. As the supply of large-effect mutants in increased further (Figure 11 where *U_β_* = 10^−9^), large-effect single mutants sweep in almost all simulations and the diversity does not have enough time to build up before a large-effect mutant sweeps.

**Figure 10:**
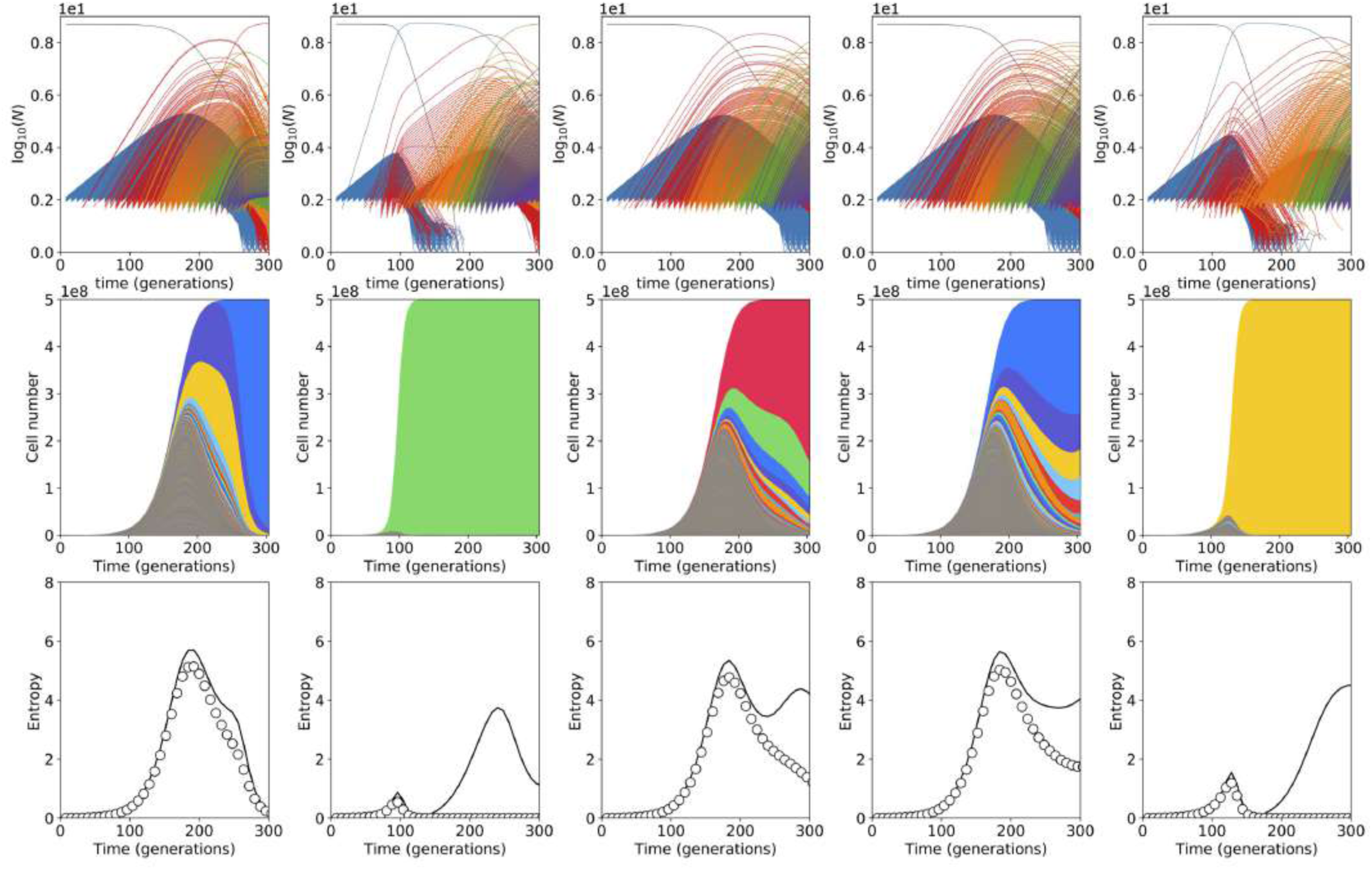
Diversity dynamics for DFE modeling growth-inhibiting drug. Here the DFE is composed of two delta-functions (at *s_α_* = 0.05 with rate *U_α_* = 10^−5^ and *s_β_* = 0.20 with rate *U_β_* = 10^−10^). Top row, the genotype trajectories of all unique genotypes that arise in the simulation (single-mutants blue, double-mutants red, triple-mutants orange, quadruple-mutants purple). Here, because the supply of large-effect single-mutants is higher, large crashes in diversity (e.g. column 1 and 5) can be caused large-effect single-mutants which sweep. Bottom row: the entropy trajectory (see Section 5.2) of all genotypes in the population (data points) compared to the theoretical prediction from using Eq. 35 in the text that assumes single mutants only. In cases where large-effect single-mutants enter early, they drive a diversity crash before single-mutant diversity has much time to build up.

**Figure 11:**
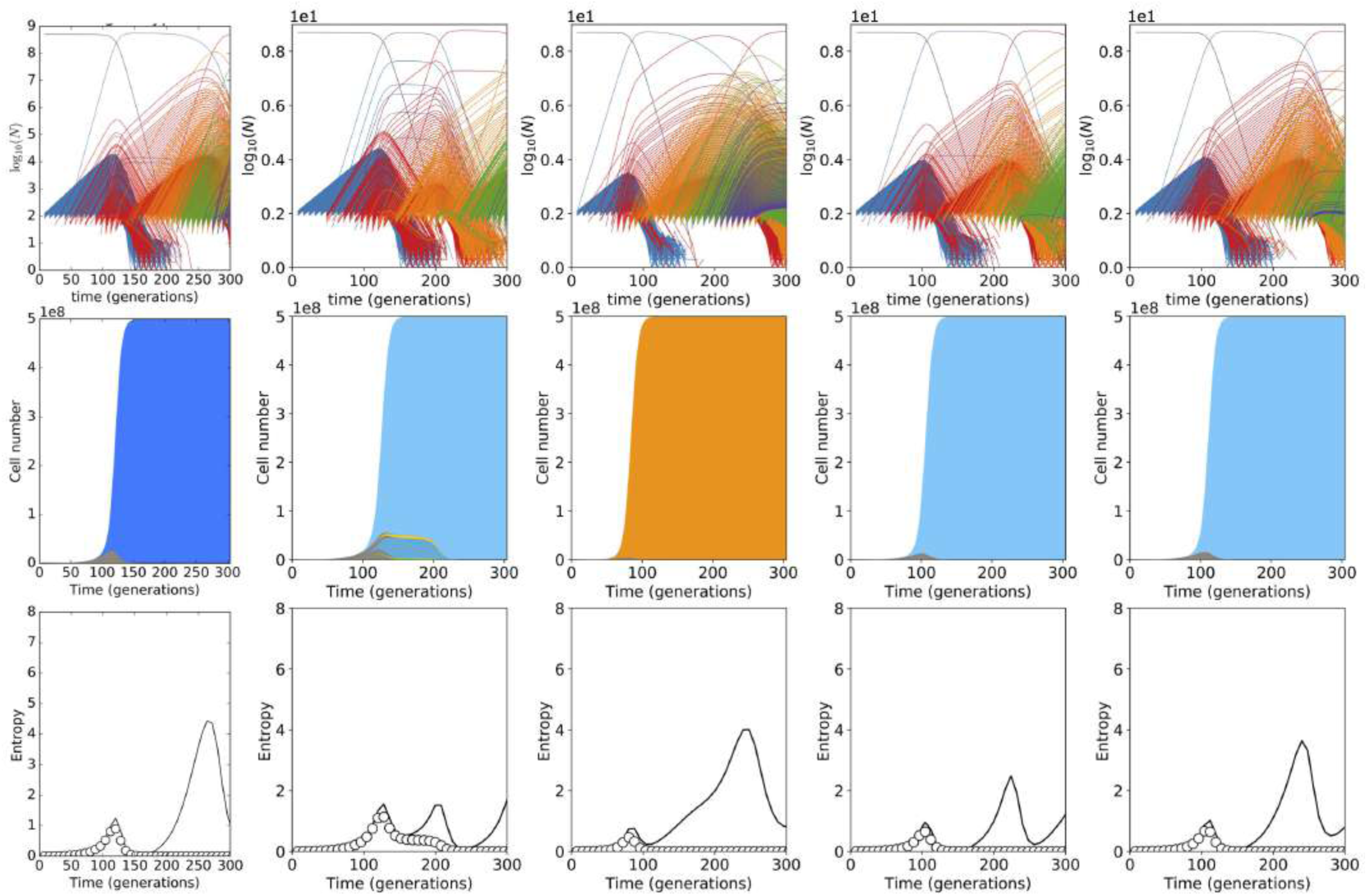
Diversity dynamics for DFE modeling growth-inhibiting drug. Here the DFE is composed of two delta-functions (at *s_α_* = 0.05 with rate *U_α_* = 10^−5^ and *s_β_* = 0.20 with rate *U_β_* = 10^−9^). Top row, the genotype trajectories of all unique genotypes that arise in the simulation (single-mutants blue, double-mutants red, triple-mutants orange, quadruple-mutants purple). With a larger supply of large-effect single-mutants almost all simulations exhibit a diversity crash driven by large-effect single-mutants which sweep. Bottom row: the entropy trajectory (see Section 5.2) of all genotypes in the population (data points) compared to the theoretical prediction from using Eq. 35 in the text that assumes single mutants only. In all cases a diversity crash caused by large-effect single-mutants occurs before diversity can build up substantially. This is followed by large oscillations in diversity occurring over timescales at which large-effect mutations enter and sweep i.e. every 1/(*NU_β_s_β_*) ~ 100 generations.

The presence of a growth-inhibiting drug could therefore be expected to alter the early adaptive diversity dynamics since stochastically occurring large-effect single-mutants could drive a crash, which would be quantitatively different from the case where the diversity crash is driven by a small number of fit multiple-mutants. It should be noted however, that in this case, the evolutionary dynamics changes more generally: it is driven by successive sweeps of large-effect mutants and is no longer truly in the clonal-interference multiple-mutation limit studied here.

#### 2.7 Power-law DFEs

In this section we explore how the dynamics of adaptive genetic diversity would be expected to evolve when the DFE has a supply of beneficial mutations spread over roughly two orders-of-magnitude. We consider DFEs where *μ*(*s*) ~ *s*^−*γ*^ with a lower limit of *s* = 0.005 (set since this mutations below this effect size would not be able to establish before the end of the observation period ~ 300 generations) and with an upper limit of *s* = 0.3 (set to remain biologically plausible, given the observed fitness effects). The total rate of beneficial mutations is set to be *U_b_* = 10^−6^.

Figure 12 shows the early time dynamics of beneficial clones for *γ* = 2. In this case, the dynamics effectively enters a successive sweep regime whereby stochastically occurring large-fitness-effect clones effectively fix before the occurrence of the *n* + 1th mutant clone. In this case, diversity has a limited time to build up and is typically purged causing characteristic humps in the entropy (last row, Figure 12).

**Figure 12:**
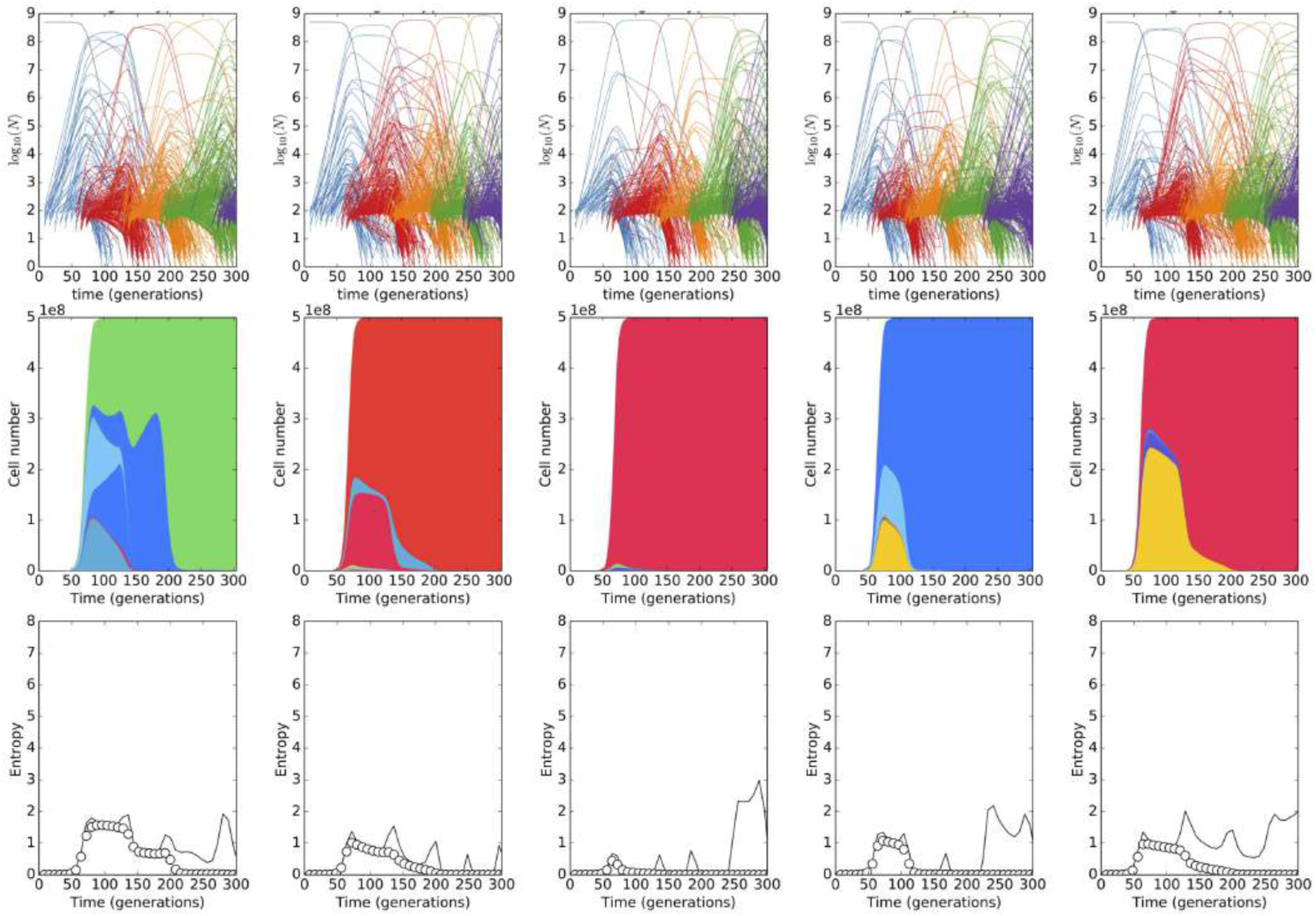
Diversity dynamics for a power-law DFE. In these simulations *μ*(*s*) ~ *s*^−2^ for 0.005 > *s* < 0.3. The total beneficial mutation rate is normalized to *U* = 10^−6^. As in previous simulations the bottleneck population size is 7 × 10^7^ which grows by a factor of 256 between bottlenecks resulting in an effective population size of ≈ 5 × 10^8^. Top row, the genotype trajectories of all unique genotypes that arise in the simulation (single-mutants blue, double-mutants red, triple-mutants orange, quadruple-mutants purple). Middle row: Muller plots of adaptive lineages. Bottom row: the entropy trajectory (see Section 5.2) of all genotypes in the population (data points) compared to the theoretical prediction from using Eq. 35 in the text that assumes single mutants only. This is followed by large oscillations in diversity occurring over timescales at which large-effect mutations enter and sweep i.e. every 1/(*NU_β_s_β_*) ~ 100 generations.

Power law DFEs thus produce qualitatively different adaptive diversity dynamics than we observe in the experimental data, though we note that a full quantitative analysis of the dynamics expected under a power-law DFE is beyond the scope of the present work [3].

### 3 Sequencing of clones, remeasuring fitness and “coloring” the DFE

#### 3.1 C-lim

In total 475 clones were picked from the C-lim evolution (369 clones from C1, blue disks in 13, 106 clones from C2, green disks in 13). A large proportion of these clones have been described previously in [5] and we refer readers to this reference for clone picking and sequencing details. Figure 13 shows the top 100 clones ranked by re-measured fitness. Anomalously large double mutant clones are highlighted by barcode numbering on the plot.

**Figure 13:**
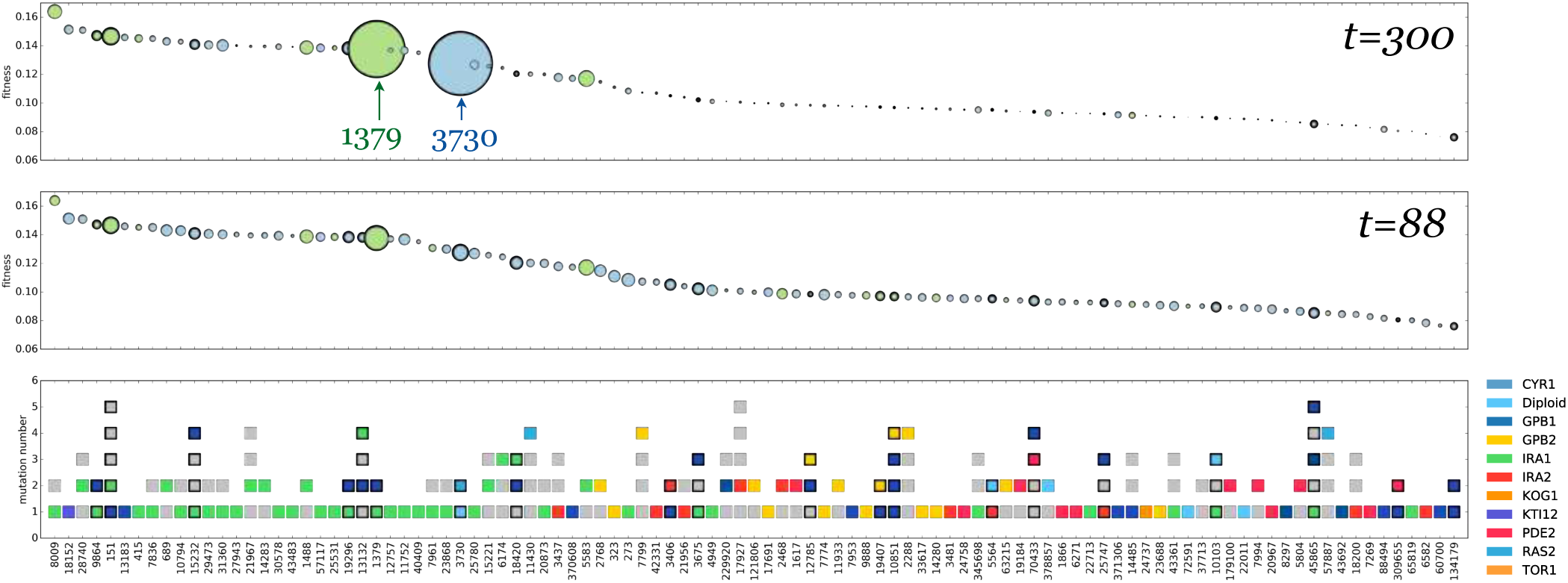
C-lim sequenced clones. The top 100 clones ranked (left to right) in order of decreasing re-measured fitness relative to the ancestor in C-lim. Each column is a clone (picked from the population at generation 88). The top panel shows the extrapolated abundance – indicated by disk area – at generation 300, the middle panel the measured abundance – indicated by disk area – at time of sampling (generation 88) and the bottom panel shows the barcode ID, and number of mutations identified in each clone (squares) with colors indicating which gene the mutation landed in. Gray indicates “other” mutations that appear only once, a substantial fraction of which are likely neutral. Multiple mutants are denoted by bold outlines and sometimes also by a dark blue square. The dark blue square indicates an adaptive mutation that did not fall into one of the commonly mutated genes listed in the key on the right-hand side.

##### Adaptive mutations

We used the following criteria to determine whether mutations were adaptive “driver” mutations (those causing the non-neutral fitness) or neutral passenger mutations.

1. If a gene was mutated multiple times in clones with distinct barcodes, mutations in that gene were designated as `adaptive`. A conservative estimate of the probability that two independently occurring neutral mutations in a given gene would be identified as adaptive via such an approach is small (~ 0.005), as the number of genes mutated twice in independent lineages is 26 and there are ~ 5, 000 possible genes. This estimate is conservative because (i) it does not use functional information and the vast majority of these mutations indicate they are functional (e.g. missense, frameshift, upstream indel) and (ii) because there is often only one mutation observed in that clone, making it more likely to be the variant causing the fitness difference.
2. If a mutation in a gene was only observed once, but that clone was clearly non-neutral (mean re-measured fitness>0.01), and no other mutations were identified in the clone, then that gene was labeled as adaptive `adaptive`.

##### Multiple-mutants

A clone that contains two or more mutations that are identified as “adaptive” via the above criteria was classified as a multiple mutant. In replicate C1 the largest clone (barcode 3730) is a confirmed double-mutant with mutations Dip+ *RAS2*. C2 is more difficult to assess as the number of clones sampled was smaller and thus a larger fraction of the high abundance barcodes at late times were not sampled at generation 88 (see Figure 23). The largest lineage in C2 — 1379 — is indeed identified as a double-mutant composed of an *IRA1* (mutated 47 times across both replicates) and *YIL169C* (mutated 6 times across both replicates). However a note of caution is warranted here: 5 of the 6 mutations in *YIL169C* are at exactly the same position. While this can occur by chance at the population sizes we use, it is more likely due to a neutral pre-existing mutation that arose prior to barcoding and was barcoded multiple times. This is especially likely in this case since the same SNP is sampled into both replicates (C1 and C2) yet is not adaptive in C1. Hence in C2 there is little evidence for early anomalously fit double-mutants. This lack of early double-mutants is consistent with the diploid trajectory in C2 (Figure 4C of main text, green data points) where the diploid trajectory is outcompeted for a long time and is rescued late, indicating a later double-mutant in this replicate.

#### 3.2 N-lim

We isolated clones from generation 192 of N1, and remeasured the fitness of the 291 clones within that pool, whose trajectories indicated they were adaptive as previously described [5]. We whole genome sequenced all clones from this pool to a mean coverage of 30x. Variants were called using the same pipeline as outlined in [5]. Details of the top 100 clones ranked by re-measured fitness is shown in Figure 14.

**Figure 14:**
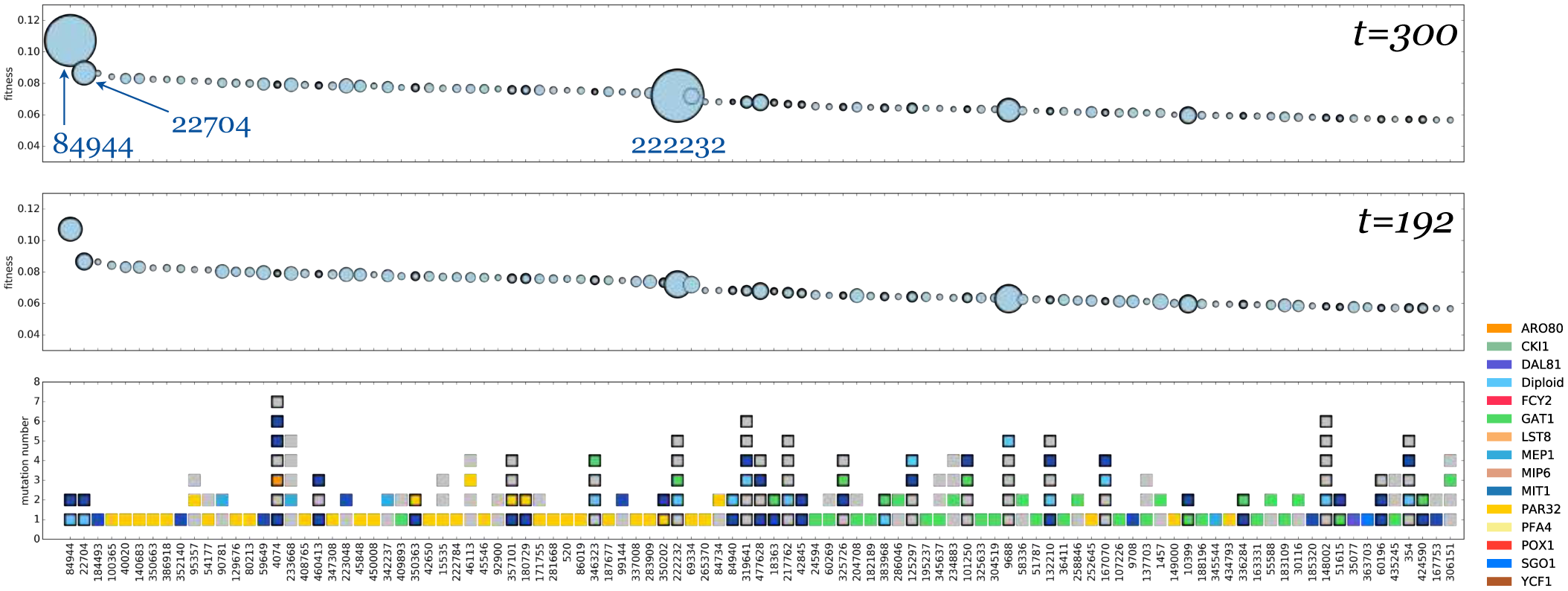
N-lim sequenced clones. The top 100 clones ranked (left to right) in order of decreasing re-measured fitness relative to the ancestor in N-lim. Each column is a clone (picked from the population at generation 192). The top panel shows the extrapolated abundance – indicated by disk area – at generation 300, the middle panel the measured abundance – indicated by disk area – at time of sampling (generation 192) and the bottom panel shows the barcode ID, and number of mutations identified in each clone (squares) with colors indicating which gene the mutation landed in. Gray indicates “other” mutations that appear only once, a substantial fraction of which are likely neutral. Multiple mutants are denoted by bold outlines and sometimes also by a dark blue square. The dark blue square indicates an adaptive mutation that did not fall into one of the commonly mutated genes listed in the key on the right-hand side.

From the whole genome sequencing data, we identified SNPs, small indels, larger deletions and insertions, Ty transposition events, and CNVs, including aneuploidy and segmental aneuploidy and annotated the genes within which those mutations fell. Among these genes, there were 39 that had more mutations than would be expected by chance including: *MEP1*, *GAT1*, *PAR32*, *FCY2*, *DAL81* and *MIT1.* In addition, we saw mutations in *MEP2* and *MEP3.* The *MEP* genes encode the ammonium permeases, suggesting that the mutations that we observe are likely to be gain of function, increasing the cell’s ability to scavenge ammonium, which is the limiting nutrient. We also observe mutations in *GAT1*, which has been previously identified as mutation in nitrogen limited chemostats [6], and encodes a GATA transcription factor that activates the nitrogen catabolite repression regulon.

##### Adaptive mutations

Adaptive mutations were identified in the same way as in the C-lim replicates (see above).

##### Multiple-mutants

In the sequenced N-lim replicate, 28 of the top 100 clones ranked by fitness are double-mutants (defined as above for C-lim). Of these, two lineages go onto dominate the population 84944 (composed of Dip+ *MEP1*) and 222232 (composed of Dip+ *GAT1*) both indicating Dip+GoF structure since the gene variants are beneficial as heterozygotes.

#### 3.3 Coloring the DFE by gene / mutation type

Combining the whole genome sequencing of adaptive clones with the lineage tracking data enables one to assign which mutations in which genes contribute to various regions of the DFE. This is shown for both N-lim and C-lim environments in Figure 2 of the main text. To “color” the beneficial mutation rate spectrum, by gene like this, we divided up the fitness range ([0, 0.15] in C-lim and [0, 0.12] in N-lim) into bins of width *δx* = 0.002. For each of the replicates with large numbers of sequenced clones (C1 and N1) we then assigned each adaptive barcode to a bin if its fitness estimate (from the maximum likelihood approach using the early time trajectory in Section 1) fell within that fitness bin. For each adaptive barcode in each fitness bin, we then asked whether a clone from this barcode was whole genome sequenced and if it was sequenced, which genes were mutated.

The contribution of a given barcode lineage to the bin rate was determined by estimating the rate using the inferred fitness effect and establishment time via

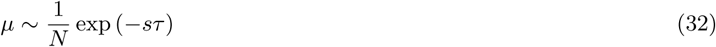

We scaled the rate of each gene in a given bin up by a factor such that the rate to all genes in a given bin added up to the total rate inferred for that bin. For example if 2 barcoded lineages each with a different mutated gene contributed to a bin, but the estimates of the rates to each were in the ratio 2:1, and the total rate to the bin was 10^−6^, then the first gene would be colored to a height of 0.67 × 10^−6^ and the second to 0.33 × 10^−6^. Bins in which no clones were sampled for sequencing were marked as unknown. This method will work well for bins in which many unique lineages that were found to be mutated, and will work less well as the number genes contributing to a bin decreases, since in the latter case there is a good chance we have under-sampled. This effect however does not affect any subsequent results, only the coloration of the DFEs in Figure 2 of the main text. Only single-mutants, which were not pre-existing (e.g. were adaptive replicate C1 (N1) and not in C2 (N2)), and which contained mutations in verified adaptive genes were counted. Verified adaptive genes are defined in section 3.1.

Multiple mutants, i.e. lineages which contain clones that upon sequencing had 2 or more mutations in genes that were independently verified as adaptive (using the above criteria) were colored differently (dark blue) so that the (small) contribution of multiple mutant lineages to the DFE could be assessed.

#### 3.4 Loss-of-Function vs. Gain-of-Function mutations

In the simulations described in section 4, we classify high-fitness mutations as either Loss-of-Function (LoF) or Gain-of-Function (GoF). This classification is inspired by observations that, in both environments, a large fraction of adaptive mutations disrupt gene function (putative LoF mutations) while a smaller fraction are likely to have modified the gene’s function (putative GoF mutations). **It should be stressed, however, that our classification of mutations as being either LoF or GoF are not definitive, and none of our conclusions depend on assigning individual clones as harboring LoF or GoF mutations. However multiple lines of evidence do point to these two broad classes of mutation being important to the diversity dynamics**. Specifically:

- **Putative LoF mutations typically occur in negative regulators**. We defined clones as harboring LoF mutations if the adaptive mutation was identified as (i) nonsense, (ii) frameshift or (iii) a large deletion or (iv) missense but in a gene in which at least one other clone harbored a mutation of type (i)-(iii) and was also identified as adaptive. Examples in C-lim of genes which are adaptive under a LoF mutation include *IRA1*, *IRA2*, *GPB2*, *PDE2* [5], examples in N-lim include *PAR32*, *GAT1*, *MIT1*, *FCY2*.
- **Putative GoF mutations often occur in positive regulators**. We defined clones as harboring GoF mutations if the adaptive mutation was identified as (i) missense and in which no other type of mutation was found to be adaptive in that gene or, in the case of N-lim, occasionally a Ty transposition if further biology could justify it as GoF (below). In C-lim, for example, mutations in positive regulators of the *RAS* pathway are exclusively missense mutations in genes such as *RAS2*, *CYR1*, *TFS1* strongly indicating they are GoF mutations. In N-lim, mutations in *MEP1*, *MEP2*, *and MEP3* genes were identified as putative GoF. The MEP genes encode ammonium permeases, suggesting that the mutations that we observe are likely to be gain of function, increasing the cell’s ability to scavenge ammonium, which is the limiting nutrient. Mutations in the MEP genes are either very specific missense mutations (sometimes hitting the same exact nucleotide) or Ty insertions downstream of the gene.
- **Putative LoF mutations are more common than GoF mutations**. Based on the mutations in Figure 4 of [5] these classes occur in the ratio of 66:16 or roughly 4:1. Making a quantitative assessment in N-lim is more challenging as definitely assigning LoF or GoF to a given mutation is harder. Thus we elected to use the same ratio 4:1 for N-lim simulations too.
- **Putative LoF mutations partially or totally recessive**. In C-lim, as predicted by our theory, the LoF mutations are almost never seen in diploids. There were however three clones which were sampled which did have LoF mutation in diploids. An *IRA1* homozygote (barcode ID 10103, re-measured fitness 9.6%), an *IRA2* homozygote (barcode ID 5564, re-measured fitness 10.0%) and an *IRA2* heterozygote (barcode ID 3577, re-measured fitness 2.8%). This suggests that high-fitness effect LoF mutations are recessive since the homozygotes retain their high fitness while the heterozygote does not.
- **Putative GoF mutations at least partially dominant**. In contrast to the paucity of LoF mutations in diploids, GoF mutations are observed in diploids and are exclusively seen in heterozygotes. Missense heterozygote mutations in *ACF2* (barcode ID 2039, re-measured fitness 8.3%), *RAS2* (barcode ID 3730, re-measured fitness 13.7%), *PSE1* (barcode ID 9689, re-measured fitness 5.7%), *VPS3* (barcode ID 15337, re-measured fitness 6.9%) and a Chromosomal amplification of Chr11 (barcode ID 3577, re-measured fitness 8.6%) are all clearly supplying a fitness benefit above and beyond the fitness effect of being a diploid (4%±1%).

These features become important when we outline the epistasis model. Since self-diploidization is a common mutation the distinction between LoF and GoF is likely to be important when considering double-mutants. Indeed, the high fitness double-mutants we observe are typically Dip+GoF mutations and are heterozygote for the GoF mutation since it occurred second. In the epistasis model outlined below we use this biological insight to reason that the order of acquiring mutations is important and profoundly changes the dynamics of genetic diversity. If LoF mutations were able to occur at the same rate on the diploid background as they do on the haploid, for example, the additive model predicts they would dominate the population of double-mutants.

### 4 Simulated lineage dynamics with inferred DFEs under 3 models

In section 2 we used theory and simulations using simple DFEs to determine genotype and lineage dynamics. Here we perform similar simulations using more realistic DFEs: those directly inferred in both N-lim and C-lim. We show that the insights from simple DFEs hold for these more realistic DFEs: single-mutants drive an expansion in genetic diversity that subsequently crashes due to the emergence of double-mutants.

To simulate the genotype and lineage dynamics with measured single-mutant DFEs, we started with the DFEs inferred in each environment (outlined in Section 1) and made the following changes:

1. Lineages that contained multiple-mutants verified via sequencing of clones (see Section 3) were removed from the DFE
2. The de-novo rate of diploidization (fitness classes below the dashed line in Figure 15) used was 3 × 10^−6^ per generation. In our previous work [5], we discovered that the large class of mutants in the 4% range in C-lim and 3% range in N-lim were diploids. Based on the high initial fraction (~ 1%) of the population which harbored a diploid mutation, it was also clear that many of these must have been induced during the transformation of barcodes rather than arising de-novo. To estimate the rate of de-novo diploidization we therefore asked what de-novo rate was most consistent with the measured diploid trajectories (Section 6.1). We found that a de-novo rate of ~ 10^−6^ was consistent with the rescue time for diploid trajectories. This rate is broadly consistent with the measured the fraction of diploids present in the population in the same experimental conditions but which did not undergo a barcode transformation. After ~ 120 generations of growth an estimated fraction of ~ 10% ± 3% of cells were diploid (Lucas Herrissant, *unpublished data*). Using the fact that this class will behave deterministically, the rate can be estimated using

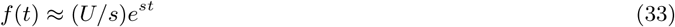

resulting in an estimated rate between 3 × 10^−6^ – 3 × 10^−5^ per generation. Uncertainties in this number should be stressed too: the exact number of diploids depends on their growth advantage in the growth conditions prior to the serial batch transfer, which are not known. **Figure 15:**
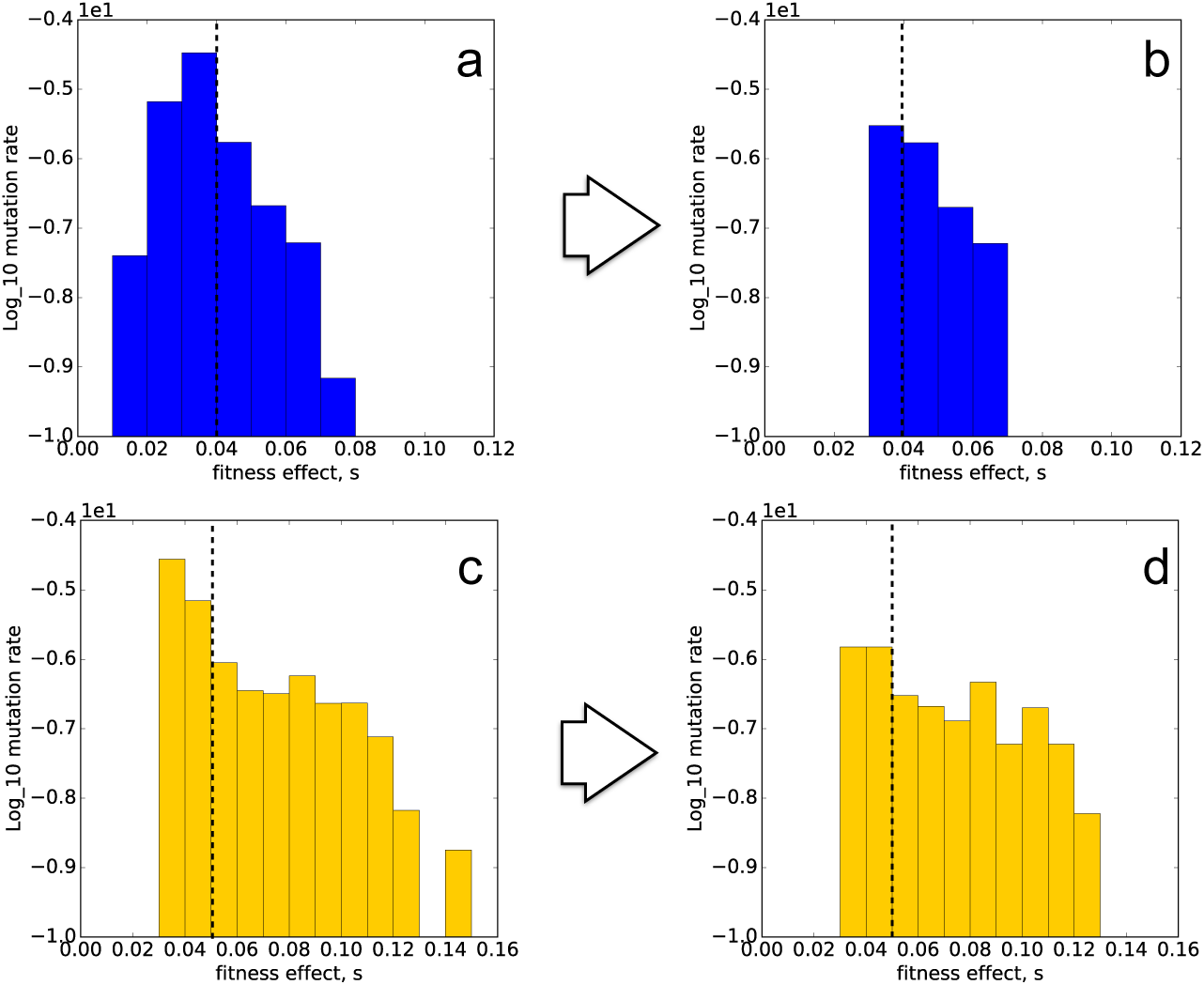
Caricature DFEs. (a) The inferred DFE in N-lim from all adaptive lineages and (b) the DFE resulting from excluding known multiple mutant lineages (from sequencing of clones) and revising the rate to Dip mutations (those below dashed line). (c) and (d) the equivalent plots for the C-lim environment. Note the logarithmic scale on the y-axis.

The results of these two adjustments are shown in Figure 15 (a → b for N-lim and c → d for C-lim). The mutations below the dashed line are all diploidizations (Dip). In both C-lim and N-lim loss-of-function (LoF) mutations occur at a higher rate than gain-of-function (GoF) mutations. As such for our caricature DFEs we assumed for simplicity that each fitness class above the dashed line has both LoF and GoF mutations which occur with relative rates of 4:1. This is broadly consistent with the number of LoF to GoF mutations seen in each environment (see Figure 4 of [5]).

#### Simulation details

Simulations were performed using a custom written python code (available on request) that simulated the fate of mutations and lineages and plotted the associated diversity measures (entropy) through time. The simulation closely follows the experimental procedure, briefly:

- A population of *N* = 5 × 10^8^ “cells” are barcoded with *L* = 500, 000 tags. The initial population of cells that gets barcoded contains 99% ancestor and 1% Dip. The distribution of initial abundances is Poisson distributed with a mean of 1000.
- Each lineage contains a dynamically updated number of genotypes (unique sets of independently occurring mutations), each of which is initially either the ancestor (WT) or the ancestor + a diploidization (WT+Dip).
- Genotypes give rise to further genotypes via acquiring mutations drawn from the DFE. The acquisition of new mutations occurs stochastically and in two steps (i) a mutation is generated (ii) a stochastic variable determines whether it establishes, and if it does, the new genotype created starts at establishment size (1/fitness advantage over the mean).
- Each generation genotypes increase in abundance according to their fitness advantage over the “mean fitness” (calculated each generation). Genetic drift (arising from small number fluctuations) is modelled implicitly in the establishment process and after this (i.e. at frequencies large enough that drift is irrelevant) lineages are modeled deterministically. Initially for example a (WT+Dip) genotype would have a fitness advantage of 4% in C-lim or 3% in N-lim since this is the fitness effect of Dip in each environment.
- The abundance of each lineage is simply the sum of the abundances of all genotypes within that lineage (which can be multiple). Genotype and lineage abundances are recorded every 8 generations.

##### 4.1 Single-mutant model

In the single-mutant-model simulations, any cell that has acquired a mutation can no longer mutate and thus only single-mutant genotypes exist and competition between these determines the entire lineage and genotype dynamics. In both environments (C-lim, Figure 16; N-lim, 17) this model produces lineage and diversity dynamics that are not in agreement with observations. The single-mutant model predicts too many lineages that remain at intermediate-high frequencies at the later time points (3rd row, “Muller” plot) as compared with the experimental lineage data. This produces elevated levels of diversity (as measured via the entropy of all adaptive lineages, 4th row) again above observations (Figure 3 of main text).

**Figure 16:**
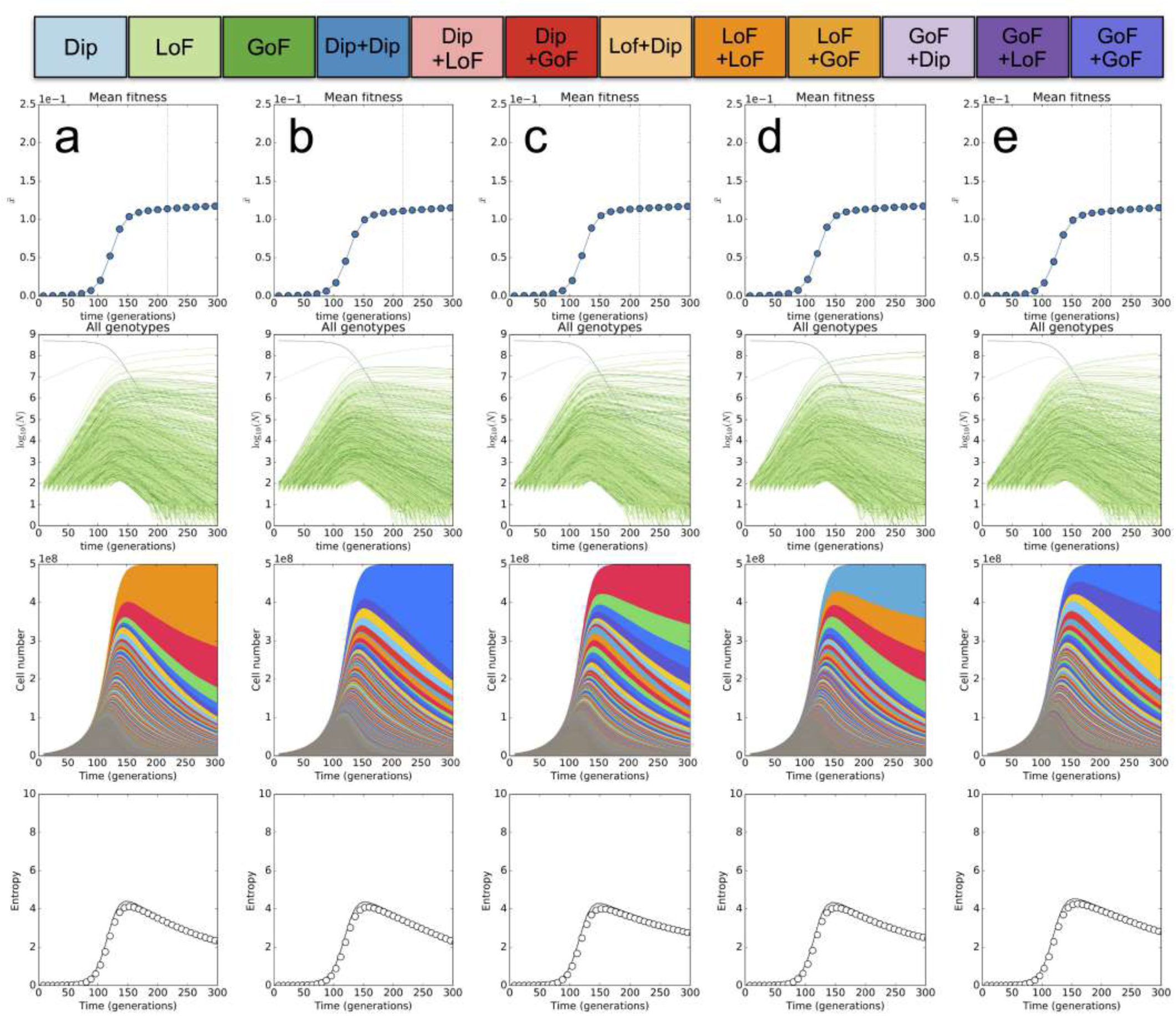
C-lim Single-mutants. (a–e) 5 randomly sampled simulated replicates of the dynamics in C-lim under a singlemutant only model. Each column is a replicate and shows the mean fitness of cells relative to the ancestor over time (1st row), the abundance of every genotype in the population through time on a log-scale (2nd row, genotypes colored according to color-key on top of figure), the abundance of each adaptive lineage (different colors are here chosen arbitrarily for visualization purposes) ranked top to bottom by lineage size at generation 300 (third row) and the entropy of all adaptive lineages through time (4th row) as measured via all genotypes (solid line) and via adaptive lineages (data points).

**Figure 17:**
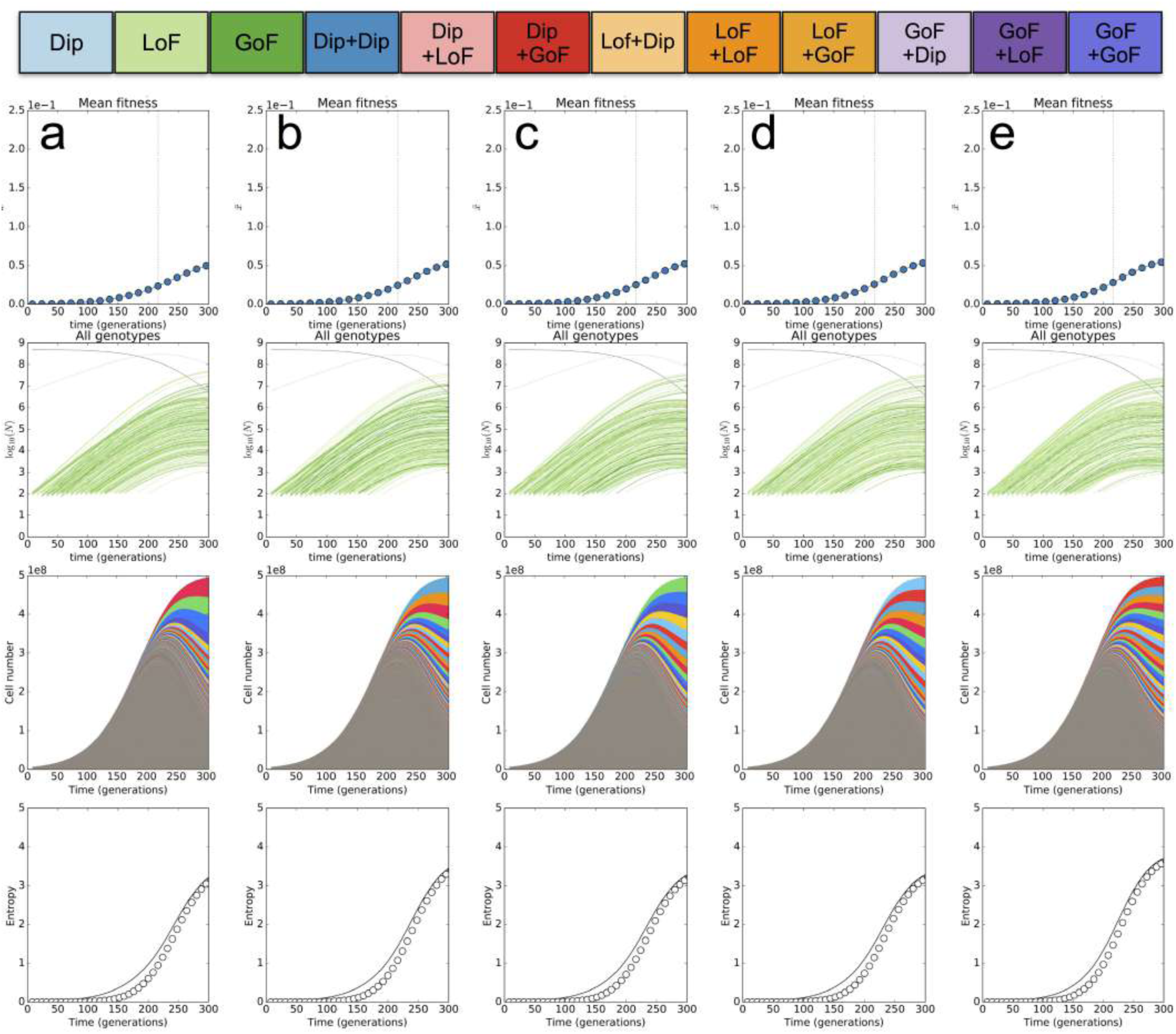
N-lim Single-mutants. (a–e) 5 randomly sampled simulated replicates of the dynamics in N-lim under a singlemutant only model. Each column is a replicate and shows the mean fitness of cells relative to the ancestor over time (1st row), the abundance of every genotype in the population through time on a log-scale (2nd row, genotypes colored according to color-key on top of figure), the abundance of each adaptive lineage (different colors are here chosen arbitrarily for visualization purposes) ranked top to bottom by lineage size at generation 300 (third row) and the entropy of all adaptive lineages through time (4th row) as measured via all genotypes (solid line) and via adaptive lineages (data points).

As expected most of the adaptation in the single-mutant model is driven by the high-fitness LoF and GoF mutations (light and dark green lines in 2nd row of both Figure 16 and 17)

There are a number of reasons this model is not accurately capturing the correct lineage and diversity dynamics. First, is that the whole genome sequencing of clones gives direct evidence that very fit double mutants are the most abundant genotypes in the population at late times in both C-lim and N-lim. Second, the adaptive lineage dynamics do not match the observed ones: competition between single mutants alone is not enough to explain the crash in genetic diversity observed across both environemtns.

##### 4.2 Additive model

In the additive simulations, cells can continue to stochastically acquire multiple adaptive mutations (waves of different colors in row 2 of Figures 18 and 19 (color key on top of plot). New mutations are drawn from the same unchanging DFE (i.e. the DFE is independent of genotype) and fitness effects of mutations combine additively. This model would be appropriate if epistasis were rare, and cells were able to acquire many different LoF and GoF mutations that do not interact.

**Figure 18:**
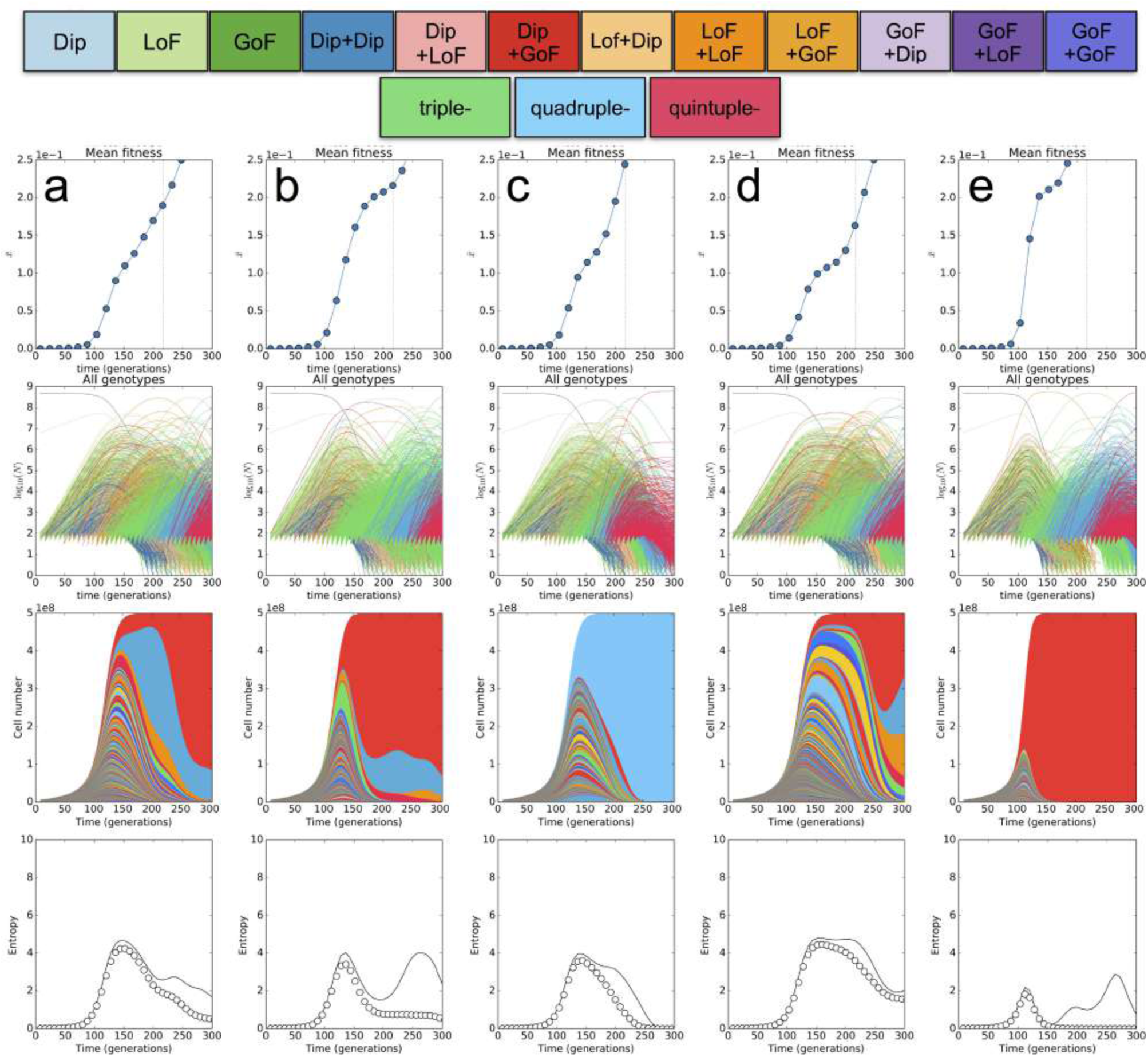
C-lim Multiple-mutants. (a–e) 5 randomly sampled simulated replicates of the dynamics in C-lim under the additive model. Each column is a replicate simulation and shows the mean fitness of cells relative to the ancestor over time (1st row), the abundance of every genotype in the population through time on a log-scale (2nd row, genotypes colored according to color-key on top of figure), the abundance of each adaptive lineage (different colors are here chosen arbitrarily for visualization purposes) ranked top to bottom by lineage size at generation 300 (third row) and the entropy of all adaptive lineages through time (4th row) as measured via all genotypes (solid line) and via adaptive lineages (data points).

**Figure 19:**
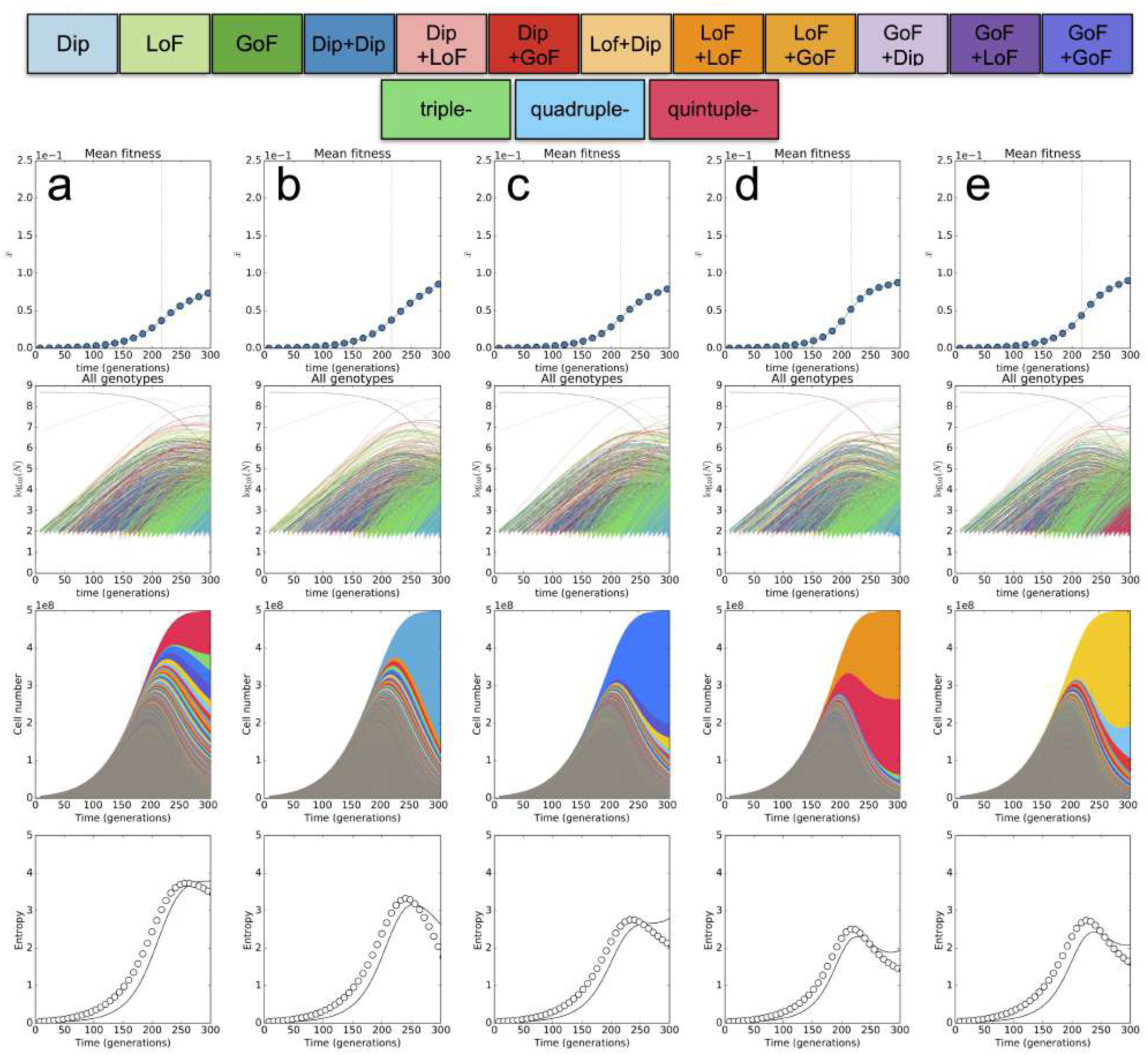
N-lim Multiple-mutants. (a–e) 5 randomly sampled simulated replicates of the dynamics in N-lim under the multiple-mutant no epistasis model. Each column is a replicate and shows the mean fitness of cells relative to the ancestor over time (1st row), the abundance of every genotype in the population through time on a log-scale (2nd row, grey is the ancestor, blue are single mutants), the abundance of each adaptive lineage (different colors) ranked top to bottom by lineage size at generation 300 (third row) and the entropy of all adaptive lineages through time (4th row) as measured via all genotypes (solid line) and via adaptive lineages (data points).

For the C-lim simulations (Figure 18) this model also produces lineage and diversity dynamics that are not in agreement with observations. First, the lineage dynamics (Figure 18) predicted by such a model are qualitatively different than those observed, whereby at intermediate-late times lineages are predicted to undergo large-scale fluctuations over long timescales (~50 generations) which we never observe in the experimental data (see columns b, d and e in Figure 18 as examples). These lineage dynamics produce trajectories of the adaptive lineage entropy which are in disagreement with observations: diversity either does not crash as dramatically and remains high even out to 300 generations as the competition between a few fit lineages continues, or it crashed too dramatically due to a very early LoF+LoF mutant (columns a and c in Figure 18 show examples). While these look qualitatively similar to the crashed observed in the data (Figure 1 of main text, and entropy plots in Figure 3 of main text) they are typically sharper: the entropy falling off more rapidly between 150-200 generations than is observed in experiment.

Another key difference between the predictions of the additive model and the observations comes from comparing the types of clones that should dominate the population at intermediate and late times. The additive model predicts that the dominant double-mutant clones over generations 150 - 300 in the 50 simulations performed are LoF+LoF (18/50), Dip+LoF (14/50), early triple mutants (11/50) GoF+LoF (4/50) or Dip+GoF (3/50). This is at odds with the sequencing of clones where we observe Dip+GoF double-mutants to be the most dominant. Furthermore, this additive model predicts that diploid trajectories (see main text Figure 4 and supplementary section 6.1) will undergo a double-dip whereby the rescue, driven by Dip+LoF clones will be outcompeted by LoF+LoF clones, which also disagrees with our data.

For N-lim simulations (Figure 19) using the additive model produces lineage dynamics which are qualitatively similar to those observed. The entropy adaptive lineage trajectories predicted by this model (see 4th row of Figure 19) are also in reasonable agreement with observed trajectories. However model predictions are in disagreement with observations in two important ways. First, the dominant intermediate/late time genotype (the one driving the diversity crash) is predicted to be Dip+LoF, which is not observed in our data. Second, the more modest fitness advantage of the haploid LoF and GoF mutations in N-lim coupled with the the expansion of Dip+LoF mutants, predict that the diploid trajectories in N-lim (see Figure 4 main text) should sweep to fixation almost every time with no dip. This is again in disagreement with data.

##### 4.3 Epsitasis model

The whole genome sequencing of hundred of adaptive clones (see Section 3) and the Diploid trajectories (see section 6.1 and main text Figure 4) produced a number of somewhat surprising results including:

- In both environments no example of a double-mutant composed of two fit single mutant SNPs was found e.g. (LoF+LoF) or (LoF+GoF) or (GoF+LoF) or (GoF + GoF).
- The double mutants observed (and that dominate the population at late times) were either Dip+GoF (and thus heterozygote for the GoF mutation since it occurred after diploidization) or (LoF+Dip) (and thus homozygote for the LoF mutation since it preceded diploidization).
- Diploid abundance in all replicate evolutions undergoes an expansion-contraction-expansion dynamics.
- Cells do not undergo further genome duplications (e.g. to become tetraploid).
- The fitnesses (those from the original lineage tracking data and the fitness re-measurements) of double mutants (WT+Dip+GoF) and (WT+LoF+Dip) combine approximately additively

Together, these observations suggested a simple model for epistasis in which, while there are 3 classes of single mutants (Dip, LoF, GoF) there are only three possible double mutants as shown Figure 4A of the main text. The key features of the epistasis model we use are:

- if first mutation is Dip then second mutations can only be GoF drawn from rates to GoF mutations in the DFE for that environment (~ 10^−8^ per generation).
- if first mutation is LoF then second mutations can only be Dip which occurs at rate of 10^−6^/gen independent of LoF mutation
- if first mutation is GoF then second mutations can only be Dip which occurs at rate of 10^−6^/gen independent of GoF mutation

The predictions for this model from 5 replicate simulations are shown in Figure 20 (C-lim) and 21 N-lim. For C-lim simulations the epistasis model produces lineage dynamics and diversity trajectories (adaptive lineage entropy trajectories) in close agreement with observations. While there remains considerable stochasticity, the dominance of a handful of doublemutant genotypes (red trajectories, 2nd row Figure 20) and corresponding dominance of a handful of lineages (3rd row) is consistent with data. Importantly, this model also predicts that overly dominant double-mutants are more likely to be caused by (Dip+GoF) and that these are most often the cause of the diversity crash.

**Figure 20:**
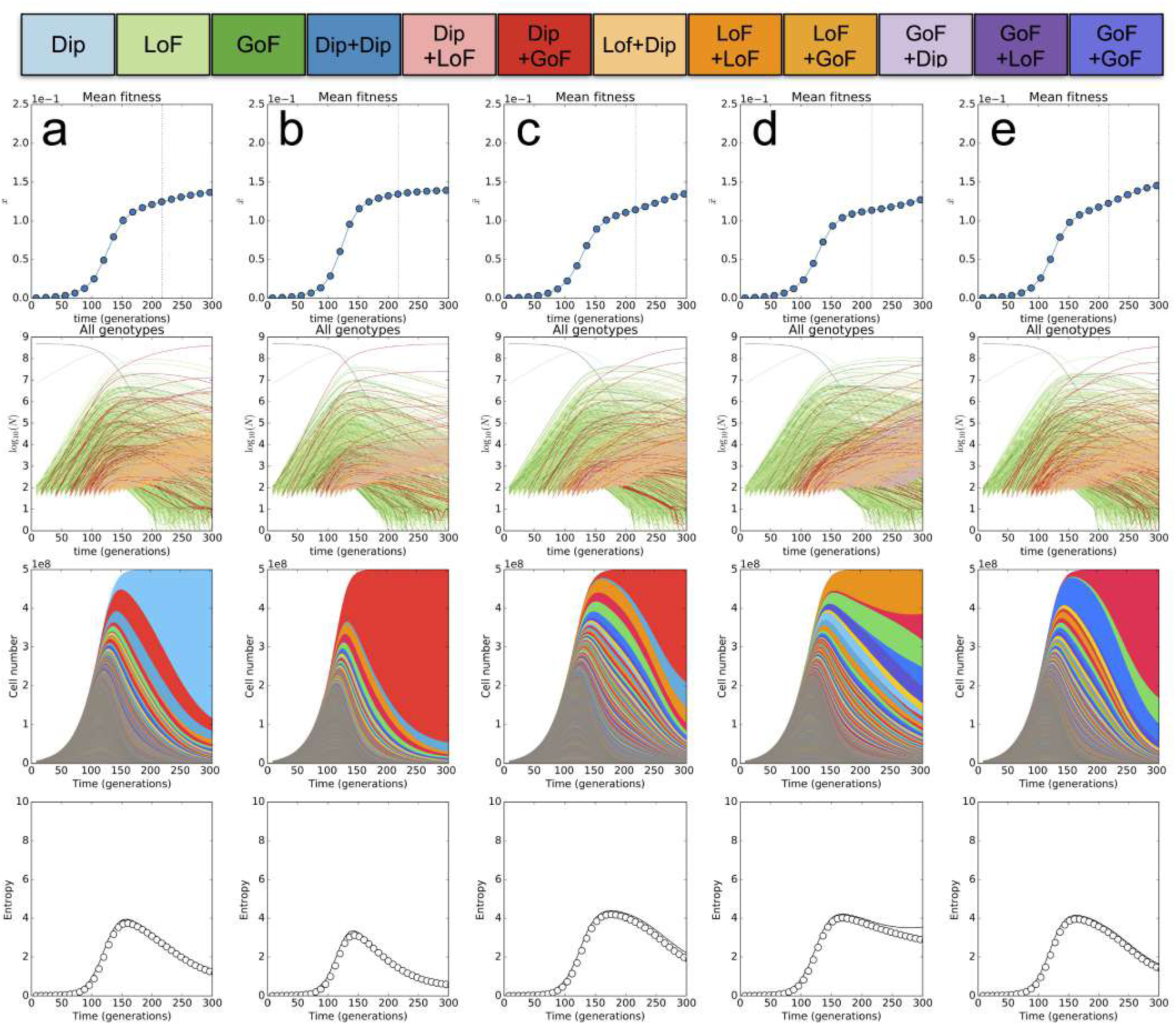
C-lim Epistasis model. (a–e) 5 randomly sampled simulated replicates of the dynamics in C-lim under the epistasis model. Each column is a replicate and shows the mean fitness of cells relative to the ancestor over time (1st row), the abundance of every genotype in the population through time on a log-scale (2nd row, grey is the ancestor, blue are single mutants), the abundance of each adaptive lineage (different colors) ranked top to bottom by lineage size at generation 300 (third row) and the entropy of all adaptive lineages through time (4th row) as measured via all genotypes (solid line) and via adaptive lineages (data points).

**Figure 21:**
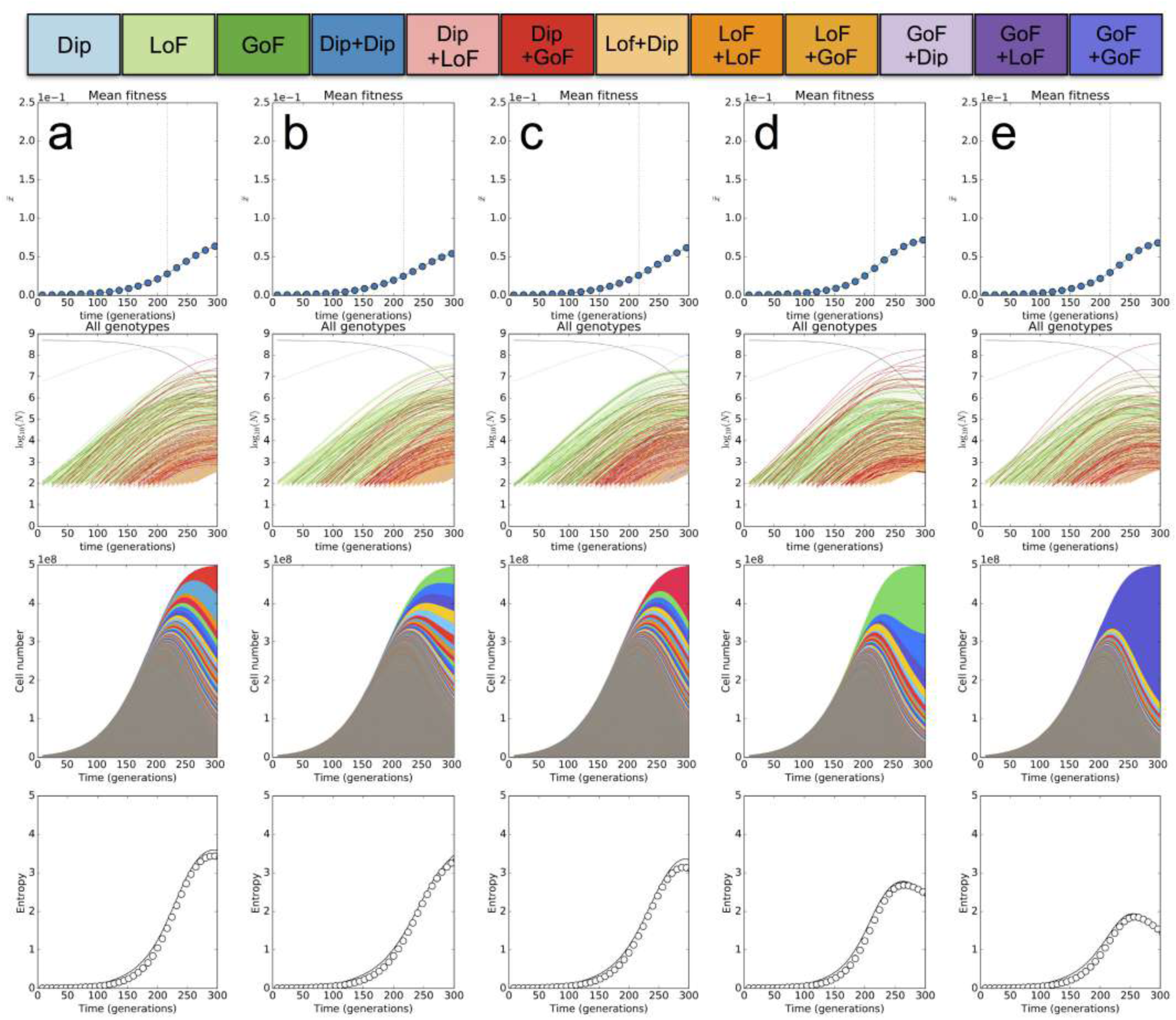
N-lim Epistasis model. (a–e) 5 randomly sampled simulated replicates of the dynamics in N-lim under the epistasis model. Each column is a replicate and shows the mean fitness of cells relative to the ancestor over time (1st row), the abundance of every genotype in the population through time on a log-scale (2nd row, grey is the ancestor, blue are single mutants), the abundance of each adaptive lineage (different colors) ranked top to bottom by lineage size at generation 300 (third row) and the entropy of all adaptive lineages through time (4th row) as measured via all genotypes (solid line) and via adaptive lineages (data points).

For N-lim simulations, lineage trajectories alone show less clear evidence that the epistasis model agrees more with observations compared to the additive model. In fact, the epistasis model produces similar lineage trajectories and slightly slower entropy expansion than is observed. However the advantage of the epistasis model is that, while it produces similar lineage dynamics, it predicts that the specific genotypes that dominate the population will be (Dip+GoF) rather than (Dip + LoF) predicted by the multiple-mutant model.

###### The possibility of further LoF and GoF mutations

This epistasis model shown in Figures 16 - 21 does not permit the the acquisition of more than two mutations. This is clearly unnatural since it is inevitable that further mutations enter at later times and expand. However this simple model appears sufficient to explain the dynamics we observe in the early stages of the experimental evolutions we performed. This suggests that single-mutants with highly fit LoF or GoF mutations have (i) fewer high fitness mutations available to them or (ii) that the fitness effects of new mutations is significantly reduced or (iii) a combination of these. Studying the emergence of multiple-mutants beyond double-mutants, however, and their impact on genetic diversity will likely require re-barcoding techniques as discussed and referenced in the main text as well as longer term whole-population sequencing efforts as in [7,8].

### 5 Lineage abundances and calculation of adaptive lineage diversity

Here we outline how we produced Figure 1 of the main text as well as trajectories for the genetic diversity (entropy) in Figure 3 of the main text.

#### 5.1 Interpolating and extrapolating lineage abundances

To plot Figure 1 of the main text we use all of the time points sequenced to best interpolate and extrapolate the abundance of each lineage in the population. To do this we use the first 104 generations of C-lim and the first 192 generations of N-lim data to estimate the fitness of each lineage using the same methods as outlined in [1], which produce fitness estimates for each lineage as well as the time evolution of the population mean fitness. We then “forecast” the abundance for times later than T104 (C-lim) or T192 (N-lim) by:

1. Using the inferred fitness *x_i_* and the initial mean fitness *x̄*(*t*_0_) to predict the frequencies at the next time point:

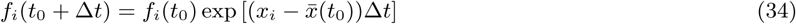
2. Recalculate the new mean fitness at this later time point:

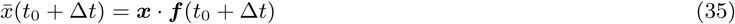

and iterate each generation until T304. The result of this procedure can be validated against the known abundance of the lineage measured from sequencing data at the later time points. Comparisons for how well this procedure works is shown in Figures 22 and 23. **Figure 22:**
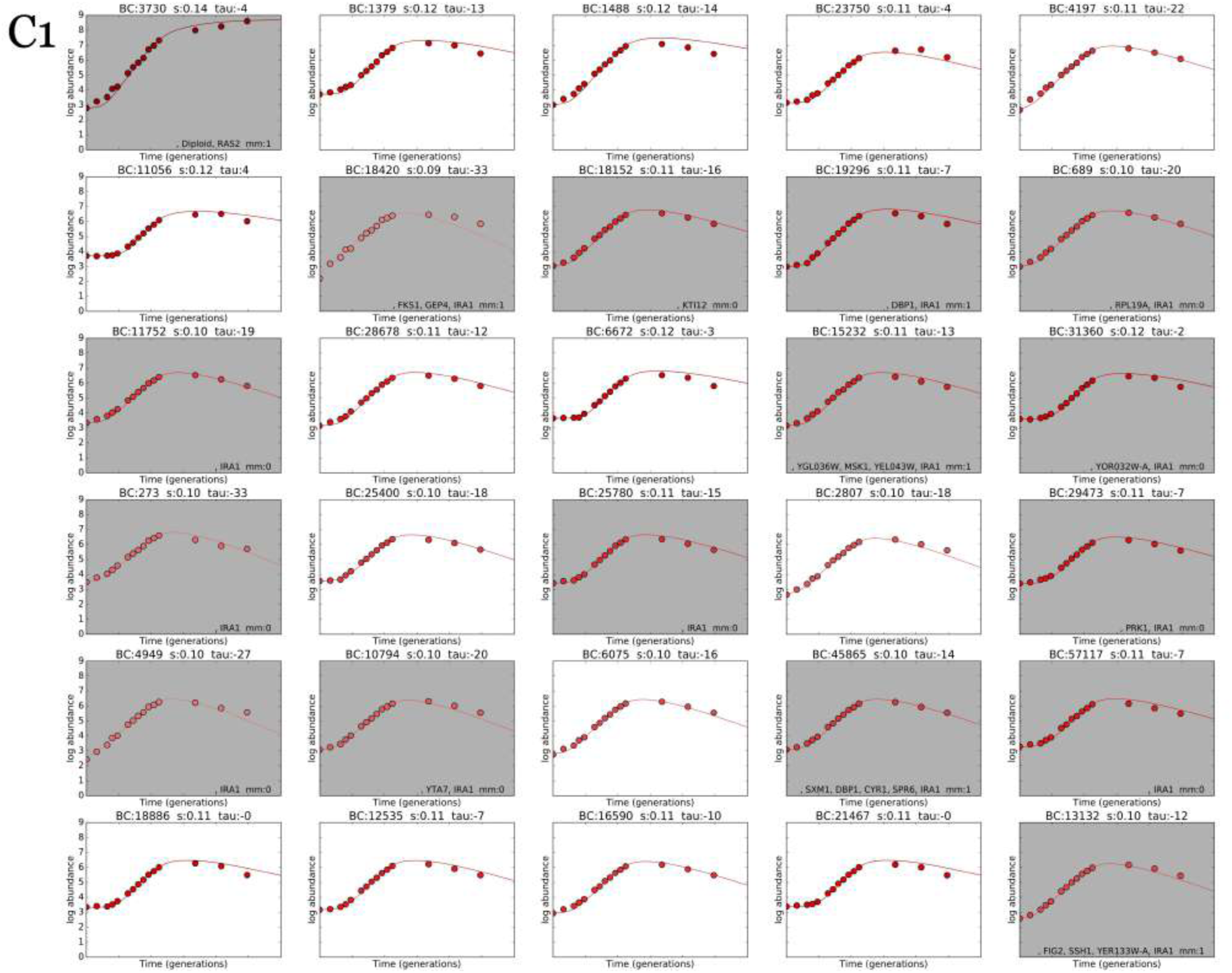
Interpolation and extrapolation of lineage abundance trajectories, C-lim replicate 1. The top 30 barcodes ranked by abundance. Each panel shows a measured barcode trajectory (data points) vs. the predicted trajectory from the extrapolation procedure (solid line). Colors indicate fitness level. Shaded panels are those lineages from which a clone was picked and subsequently whole-genome sequenced.

**Figure 23:**
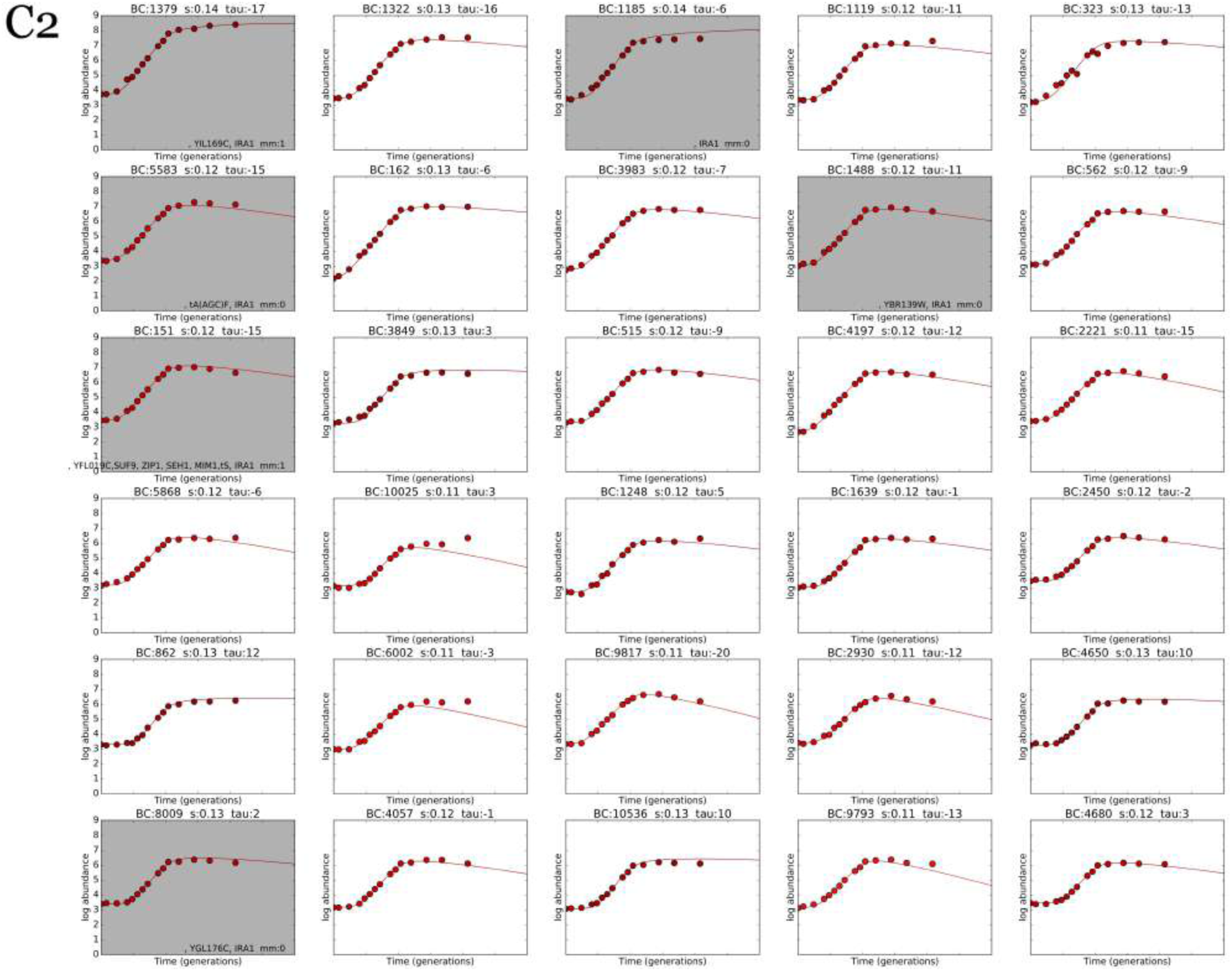
Interpolation and extrapolation of lineage abundance trajectories, C-lim replicate 1. The top 30 barcodes ranked by abundance. Each panel shows a measured barcode trajectory (data points) vs. the predicted trajectory from the extrapolation procedure (solid line). Colors indicate fitness level. Shaded panels are those lineages from which a clone was picked and subsequently whole-genome sequenced.

#### 5.2 Diversity measured via adaptive lineage entropy

To quantify genetic diversity we use the Shannon entropy of adaptive lineage through time defined here as

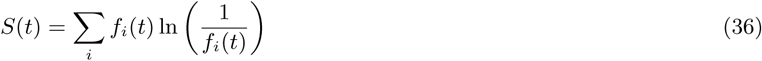

where *f_i_*(*t*) is the frequency of lineage *i* at time *t* and the sum is over all *adaptive* lineages (excluding diploids) identified using the same method as in [1] (using lineage deviation from a neutral expectation). The numerical value of *S* can be interpreted, very roughly, as there being effectively ~ *e^S^* adaptive lineages present in the population. Note that we exclude diploids from our definition because most diploid lineages are genetically identical despite adaptive clones arising in a large number of lineages. SNPs by contrast are typically genetically unique.

##### Other possible measures of diversity

The diversity is really a statement about the distribution of adaptive clones sizes and how this changes through time as shown by the colored lines in the 2nd rows of Figures 16 - 21.

However, if one wants to compare many replicates it becomes difficult unless one chooses a statistic based on the distribution of clone or lineage abundances. The entropy is one such measure. We do not claim that the entropy is the “correct” measure of diversity since there are many alternatives, but it *does* capture the important qualitative distinction between the early vs. late distribution of clone sizes in the populations. Early, there are many small clones of roughly equal size, whereas late there is a very skewed distribution with a small number of very large clones, and the entropy measure captures this difference as discussed in Section 2.5 and in the bottom rows of Figures 16 - 21. Crucially, the entropy of adaptive lineages tracks the entropy of adaptive genotypes into the diversity crash.

One consideration for a diversity measure is how much to weight low frequency lineages. Consider the case where one adaptive lineage is large, at frequency *p* while the *k* ≫ 1 other lineages are at frequency 1 – *p*/*k*. In this case the contribution from the low frequency lineages would be (1 – *p*)log(*k*/(1 – *p*)) which would mean that the low frequency lineages would contribute more to the entropy than the dominant lineage when *p* < 1 – 1/log(*k*). A reasonable alternative measure of diversity, would be to weight variants not with log(*p_i_*) (as one does in the entropy measure), but with a different functional form. For example, if we used a diversity measure:

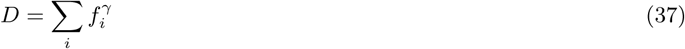

where 0 ≥ *γ* ≤ 1, the contribution from the low frequency lineages would be enhanced over simple linear weighting, (and in the limit of *γ* = 0 it is simply a count of how many are present). Following the same line of argument as above, the contribution of the low frequency lineages would be larger than the single large lineage if 1 – *p* < *k*^1–1/*γ*^.

Other reasonable choices of the diversity measure could be made. One simple example would be a step function counting how many lineages reach above a threshold size *p*_0_ over time. Because many adaptive lineages reach frequencies above 10^−5^, but many of them dip below this frequency at late times, for parameter values 10^−5^ < *p*_0_ < 10^−3^, this measure would be qualitatively similar to the entropy measure, though has the drawback of having to make a somewhat arbitrary choice for *p*_0_.

### 6 Diploid trajectories

#### 6.1 Measuring diploid abundance through time

We suspended cells (from either the carbon or nitrogen limited evolution) from a frozen stock into 1mL of water. We used a coulter counter (Beckman Coulter: 6605700) to quantify cell concentration, then used 5-8 glass beads to spread ~ 400 cells onto Nunc plates (ThermoFisher Cat #: 267060) containing YPD agar (10g yeast extract Fisher: 212750, 20g peptone Fisher: 211677, 20g dextrose Sigma Aldrich: G8270-5KG, 20g agar Fisher: 214010, bring up to 1L with DO water) supplemented with 2x G418 (8mL of 50mg/mL stock into 1L YPD) and allowed them to grow for two days at 30°C. Then we scanned plates at 600 dpi resolution and inputted the resulting images into a custom written MATLAB code (available on request), which infers coordinates for individual colonies based on the scanned image. We then used a ROTOR STINGER robot (Singer Instruments) to pick colonies into 96 well plates (ThermoFisher: 265301) containing YPD with 2x G418. These plates were placed in a 30°C incubator and grown overnight. We next used the ROTOR STINGER robot to replicate the 96 well plates onto a square plate with Benomyl media, and control plate with DMSO (Sigma Aldrich: 472301) media. The Benomyl media was made from a 10mg/ml stock of Benomyl (Sigma Adrich Cat #: 381586-5G) in DMSO by adding 2 mL of stock Benomyl drop-wise while stirring to one liter of YPD that had been allowed to cool to about 55°C. The final concentration of the Benomyl media was 10 *μ*g/ml. The cells replicated onto Benomyl and DMSO plates were grown up for 2 days at 30° C. After 48 hours growth, each plate was scanned. Haploids are able to grow better on Benomyl media while diploids grow more slowly. For each colony that grew on the control plate, we checked the corresponding positions on Benomyl plate and counted how many grew poorly (see Figure 24). Diploid frequencies were then calculated as the fraction of those colonies that grew poorly.

**Figure 24:**
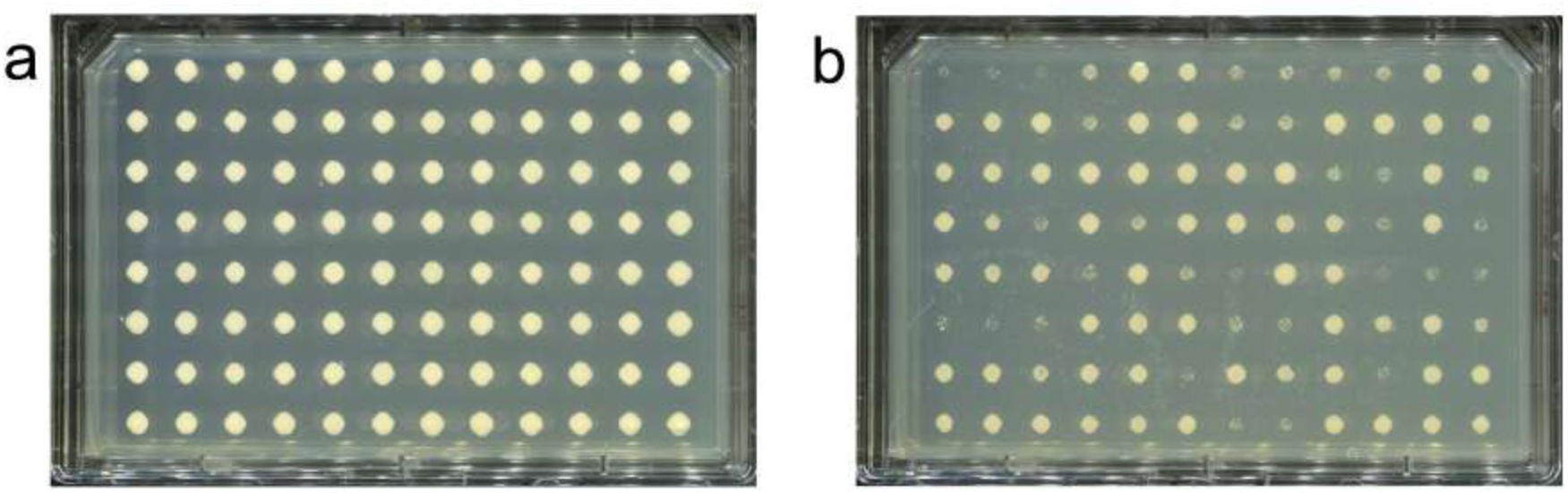
Benomyl assay for quantifying diploid abundances. Example plates showing the growth of 96 randomly sampled colonies from the population on a control plate (YPD) (a) and on a plate containing benomyl (YPD+Benomyl) (b) where the growth of diploids is inhibited.

#### 6.2 Predicted diploid trajectories under the 3 models

To track diploid trajectories, one does not need lineage information. Thus, diploid trajectory simulations were implemented independently since they are far more efficient without the large number of lineage labels that are required in the simulations outlined in Section 4.

The frequency trajectory of diploids in the population though time is then calculated using the following stochastic simulation:

- The initial abundance of diploids is set at 1%, based on direct measurement from the colony growth assay.
- The initial abundance of fitter LoF and GoF mutations is generated stochastically by realizing that their effective starting number will be roughly Poisson distributed with mean *n*_0_ = *NU_LOF_* or *n*_0_ = *NU_GOF_* which then grow deterministically through time as *n*_0_*e^st^*/*s*. (parameter values in the table below). Drift for the single mutants can be safely ignored since *n*_0_ for each of the three classes is a large parameter i.e *NU_LoF_* ≫ 1 and *NU_GoF_* ≫ 1.
- These three expanding classes can each acquire further mutations whose mutation rates and fitness effects are outlined in the tables below. These double mutants change in frequency between time points by generating a Poisson sample the new expected number of cells and normalizing by *N* in order to include the stochasticity of drift: which is crucial for the double-mutants.

Each model is specified by a number of parameters:

**Table 3:** Values used for the simulations of diploid trajectories in C-lim.

| Mutant | Measured fitness | Measured rate | Single $s$ & $U$ | Additive $s$ & $U$ | Epistasis $s$ & $U$ |
| --- | --- | --- | --- | --- | --- |
| *Dip* | 0.04 | $10^{-6}$ | — | — | — |
| *LoF* | 0.10 | $10^{-7}$ | — | — | — |
| *GoF* | 0.10 | $10^{-8}$ | — | — | — |
| *Dip + Dip* | | — | — | $0.08$ & $10^{-6}$ | — |
| *Dip + LoF* | | — | — | $0.14$ & $7.5 \times 10^{-7}$ | — |
| *Dip + GoF* | | — | — | $0.14$ & $7.5 \times 10^{-8}$ | $0.14$ & $10^{-8}$ |
| *LoF + Dip* | | — | — | $0.14$ & $10^{-6}$ | $0.14$ & $10^{-6}$ |
| *LoF + LoF* | | — | — | $0.20$ & $7.5 \times 10^{-7}$ | — |
| *LoF + GoF* | | — | — | $0.20$ & $7.5 \times 10^{-8}$ | — |
| *GoF + Dip* | | — | — | $0.14$ & $10^{-6}$ | $0.14$ & $10^{-6}$ |
| *GoF + LoF* | | — | — | $0.20$ & $7.5 \times 10^{-7}$ | — |
| *GoF + GoF* | | — | — | $0.20$ & $7.5 \times 10^{-8}$ | — |

**Table 4:**
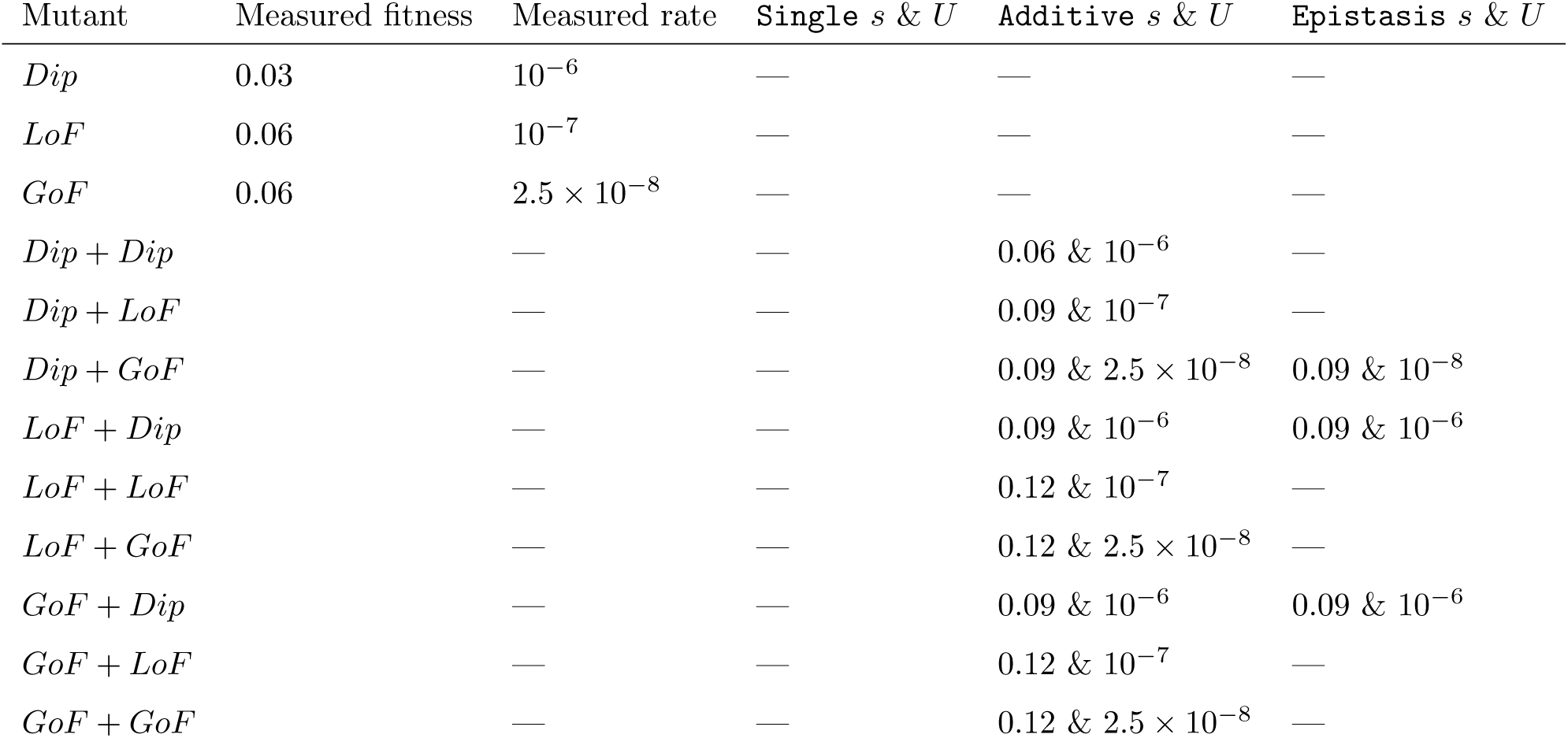
Values used for the simulations of diploid trajectories in N-lim.

**Figure 25:**
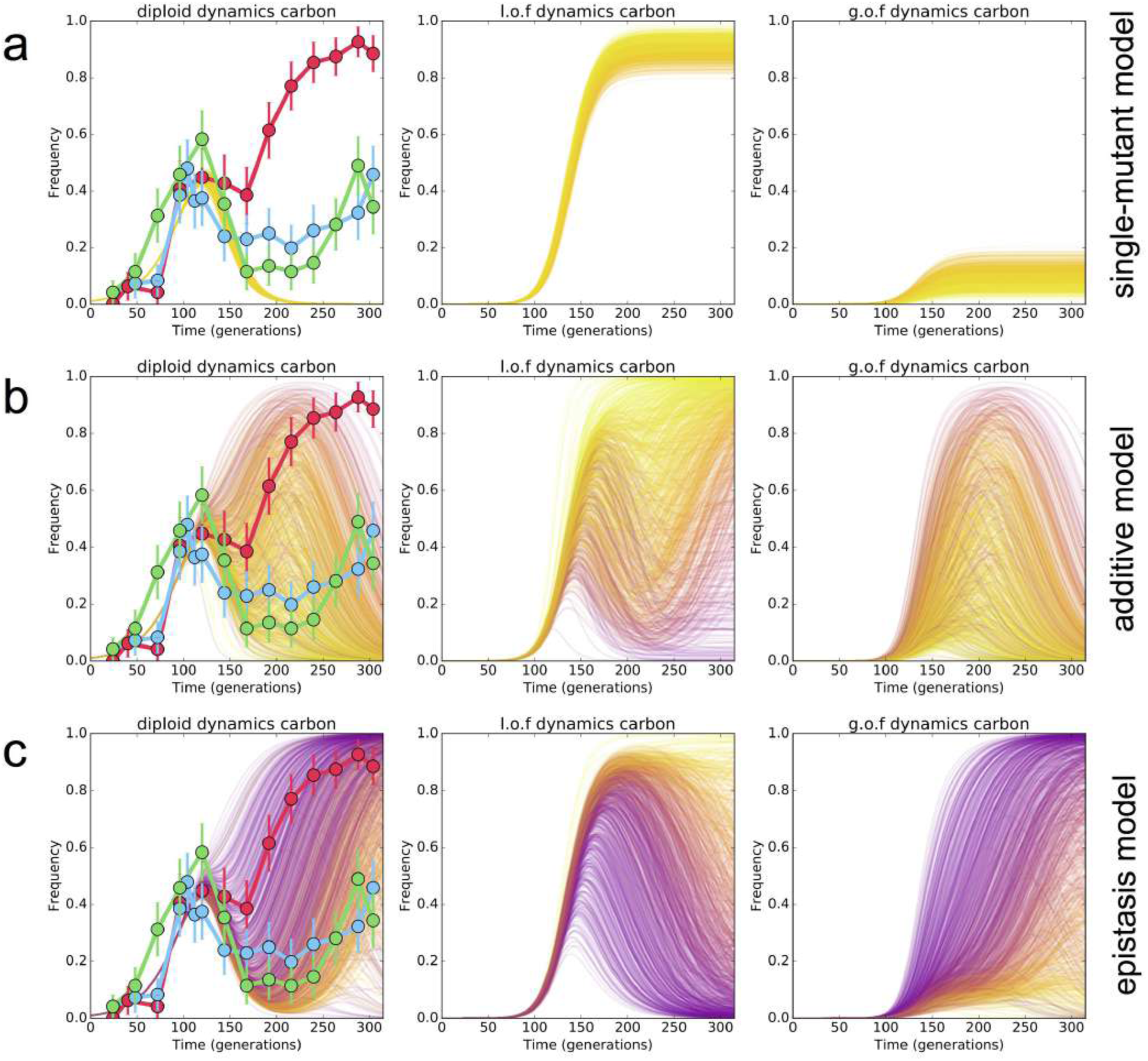
C-lim diploid, LoF and GoF trajectories for 3 models of mutation acquisition. (a) single-mutant model (b) additive model and, (c), epistasis model. The 2nd and 3rd columns show the trajectory of LoF and GoF mutants as classes. Coloring is consistent across the three plots so that a yellow trajectory in one column is the same color across all three columns since it comes from the same simulation.

**Figure 26:**
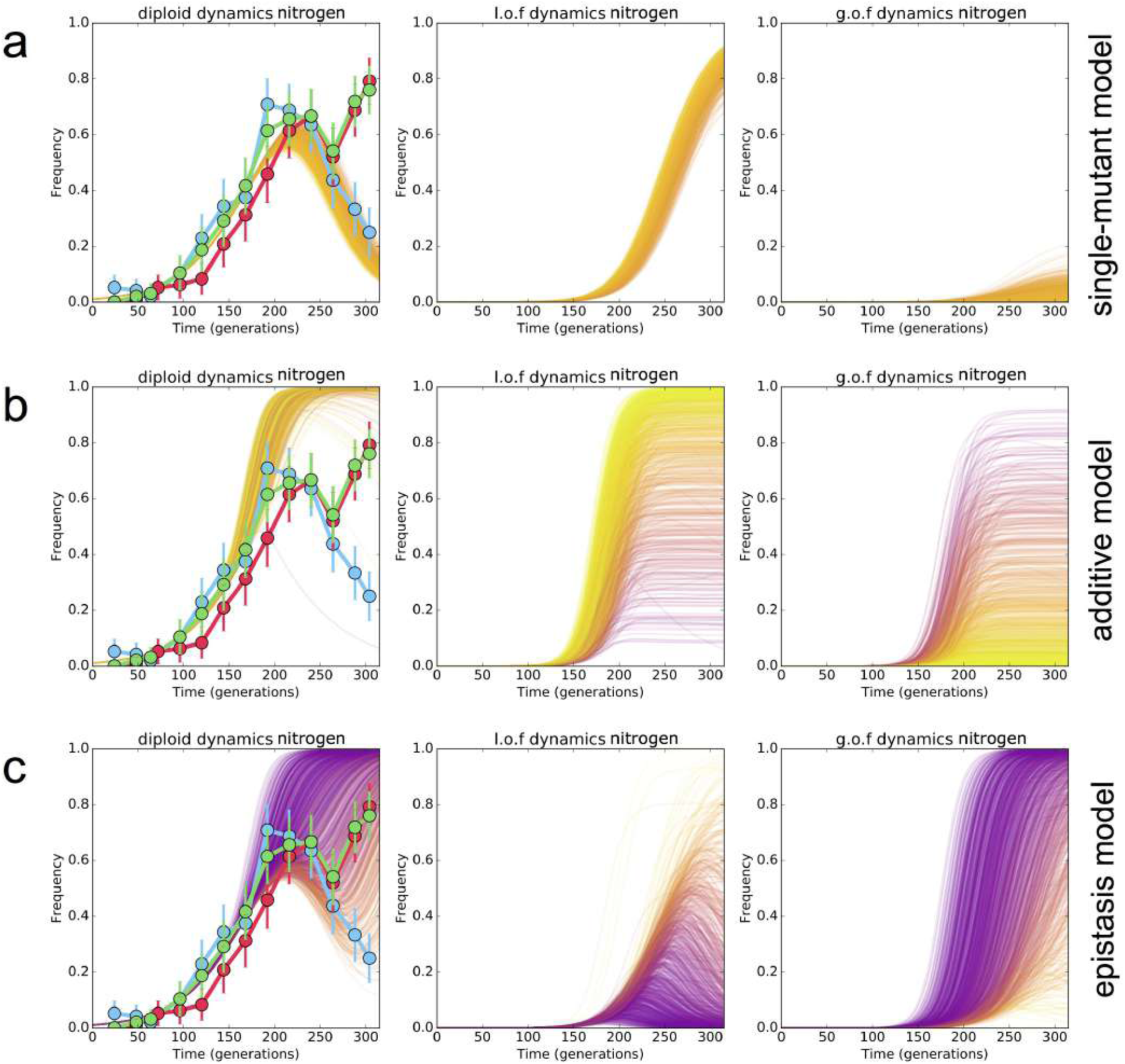
N-lim diploid, LoF and GoF trajectories for 3 models of mutation acquisition. (a) single-mutant model (b) additive model and, (c), epistasis model. The 2nd and 3rd columns show the trajectory of LoF and GoF mutants as classes. Coloring is consistent across the three plots so that a yellow trajectory in one column is the same color across all three columns since it comes from the same simulation.

## References

1. Desai, M. M., Walczak, A. M. & Fisher, D. S. Genetic Diversity and the Structure of Genealogies in Rapidly Adapting Populations. Genetics 193, 565–585 (2013).

2. Neher, R. A. & Hallatschek, O. Genealogies of rapidly adapting populations. PNAS 110, 437–442 (2013).

3. Nowell, P. C. The clonal evolution of tumor cell populations. Science 194, 23–28 (1976).

4. Tenaillon, O. et al. Tempo and mode of genome evolution in a 50,000-generation experiment. Nature 536, 165–170 (2016).

5. Lang, G. I. et al. Pervasive genetic hitchhiking and clonal interference in forty evolving yeast populations. Nature 500, 571–574 (2013).

6. Baym, M. et al. Spatiotemporal microbial evolution on antibiotic landscapes. Science 353, 1147–1151 (2016).

7. Lawrence, M. S. et al. Mutational heterogeneity in cancer and the search for new cancer-associated genes. Nature 499, 214–218 (2013).

8. Landau, D. A. et al. Mutations driving CLL and their evolution in progression and relapse. Nature 526, 525–530 (2011).

9. Gerlinger, M. et al. Intratumor Heterogeneity and Branched Evolution Revealed by Multiregion Sequencing. New England Journal of Medicine 366, 883–892 (2012).

10. Jan, M. et al. Clonal Evolution of Preleukemic Hematopoietic Stem Cells Precedes Human Acute Myeloid Leukemia. Science Translational Medicine 4, 149ra118–149ra118 (2012).

11. Nik-Zainal, S. et al. The Life History of 21 Breast Cancers. Cell 149, 994–1007 (2012).

12. Neher, R. A., Russell, C. A. & Shraiman, B. I. Predicting evolution from the shape of genealogical trees. eLife 3, (2014).

13. Didelot, X., Walker, A. S., Peto, T. E., Crook, D. W. & Wilson, D. J. Within-host evolution of bacterial pathogens. Nat Rev Micro 14, 150–162 (2016).

14. Smith, J. M. & Haigh, J. The hitch-hiking effect of a favourable gene. Genetics Research 23, 23–35 (1974).

15. Messer, P. W. & Neher, R. A. Estimating the Strength of Selective Sweeps from Deep Population Diversity Data. Genetics 191, 593–605 (2012).

16. Gerrish, P. J. & Lenski, R. E. The fate of competing beneficial mutations in an asexual population. Genetica 102–103, 127 (1998).

17. Desai, M. M. & Fisher, D. S. Beneficial Mutation–Selection Balance and the Effect of Linkage on Positive Selection. Genetics 176, 1759–1798 (2007).

18. Kao, K. C. & Sherlock, G. Molecular characterization of clonal interference during adaptive evolution in asexual populations of Saccharomyces cerevisiae. Nat Genet 40, 1499–1504 (2008).

19. Good, B. H., Rouzine, I. M., Balick, D. J., Hallatschek, O. & Desai, M. M. Distribution of fixed beneficial mutations and the rate of adaptation in asexual populations. PNAS 109, 4950–4955 (2012).

20. Buskirk, S. W., Peace, R. E. & Lang, G. I. Hitchhiking and epistasis give rise to cohort dynamics in adapting populations. bioRxiv 106732 (2017). doi:10.1101/106732

21. Lässig, M., Mustonen, V. & Walczak, A. M. Predicting evolution. Nature Ecology & Evolution 1, 77 (2017).

22. Levy, S. F. et al. Quantitative evolutionary dynamics using high-resolution lineage tracking. Nature 519, 181–186 (2015).

23. Venkataram, S. et al. Development of a Comprehensive Genotype-to-Fitness Map of Adaptation-Driving Mutations in Yeast. Cell 166, 1585–1596.e22 (2016).

24. Jost, L. Entropy and diversity. Oikos 113, 363–375 (2006).

25. Luria, S. E. & Delbrück, M. Mutations of Bacteria from Virus Sensitivity to Virus Resistance. Genetics 28, 491–511 (1943).

26. Hughes, D. & Andersson, D. I. Evolutionary consequences of drug resistance: shared principles across diverse targets and organisms. Nat Rev Genet 16, 459–471 (2015).

27. Fusco, D., Gralka, M., Kayser, J., Anderson, A. & Hallatschek, O. Excess of mutational jackpot events in expanding populations revealed by spatial Luria–Delbrück experiments. Nature Communications 7, 12760 (2016).

28. Fisher, D. S. Asexual evolution waves: fluctuations and universality. J. Stat. Mech. 2013, P01011 (2013).

29. Park, S.-C. & Krug, J. Clonal interference in large populations. PNAS 104, 18135–18140 (2007).

30. Jaffe, M., Sherlock, G. & Levy, S. F. iSeq: A New Double-Barcode Method for Detecting Dynamic Genetic Interactions in Yeast. G3: Genes, Genomes, Genetics 7, 143–153 (2017).

31. McKenna, A. et al. Whole-organism lineage tracing by combinatorial and cumulative genome editing. Science 353, aaf7907 (2016).

32. Perli, S. D., Cui, C. H. & Lu, T. K. Continuous genetic recording with self-targeting CRISPR-Cas in human cells. Science 353, aag0511 (2016).

33. Shipman, S. L., Nivala, J., Macklis, J. D. & Church, G. M. Molecular recordings by directed CRISPR spacer acquisition. Science 353, aaf1175 (2016).

## References

[1] Sasha F. Levy, Jamie R. Blundell, Sandeep Venkataram, Dmitri A. Petrov, Daniel S. Fisher, and Gavin Sherlock. Quantitative evolutionary dynamics using high-resolution lineage tracking. <u>Nature</u>, 519(7542):181–186, March 2015. ISSN 0028-0836. URL http://www.nature.com/nature/journal/v519/n7542/abs/nature14279.html.

[2] Cornelis Verduyn, Erik Postma, W. Alexander Scheffers, and Johannes P. Van Dijken. Effect of benzoic acid on metabolic fluxes in yeasts: A continuous-culture study on the regulation of respiration and alcoholic fermentation. <u>Yeast</u>, 8(7): 501–517, July 1992. ISSN 1097-0061. URL http://onlinelibrary.wiley.com/doi/10.1002/yea.320080703/abstract.

[3] Michael M. Desai and Daniel S. Fisher. Beneficial mutation selection Balance and the Effect of Linkage on Positive Selection. <u>Genetics</u>, 176(3):1759–1798, July 2007. ISSN 0016–6731, 1943–2631. URL http://www.genetics.org/content/176/3/1759.

[4] Philipp W. Messer and Richard A. Neher. Estimating the Strength of Selective Sweeps from Deep Population Diversity Data. <u>Genetics</u>, 191(2):593–605, June 2012. ISSN 0016–6731, 1943–2631. URL http://www.genetics.org/content/191/2/593.

[5] Sandeep Venkataram, Barbara Dunn, Yuping Li, Atish Agarwala, Jessica Chang, Emily R. Ebel, Kerry Geiler-Samerotte, Lucas Herrissant, Jamie R. Blundell, Sasha F. Levy, Daniel S. Fisher, Gavin Sherlock, and Dmitri A. Petrov. Development of a Comprehensive Genotype-to-Fitness Map of Adaptation-Driving Mutations in Yeast. <u>Cell</u>, 0(0), September 2016. ISSN 0092–8674, 1097–4172. URL /cell/abstract/S0092-8674(16)31010-8.

[6] Jungeui Hong and David Gresham. Molecular Specificity, Convergence and Constraint Shape Adaptive Evolution in Nutrient-Poor Environments. <u>PLOS Genetics</u>, 10(1):e1004041, January 2014. ISSN 1553–7404. URL http://journals.plos.org/plosgenetics/article?id=10.1371/journal.pgen.1004041.

[7] Gregory I. Lang, Daniel P. Rice, Mark J. Hickman, Erica Sodergren, George M. Weinstock, David Botstein, and Michael M. Desai. Pervasive genetic hitchhiking and clonal interference in forty evolving yeast populations. <u>Nature</u>, 500(7464):571–574, August 2013. ISSN 0028–0836. URL http://www.nature.com/nature/journal/v500/n7464/full/nature12344.html.

[8] Daniel J. Kvitek and Gavin Sherlock. Whole Genome, Whole Population Sequencing Reveals That Loss of Signaling Networks Is the Major Adaptive Strategy in a Constant Environment. <u>PLoS Genet</u>, 9(11):e1003972, November 2013. URL http://dx.doi.org/10.1371/journal.pgen.1003972.

